# An integrated single cell and spatial omics atlas of human prenatal development

**DOI:** 10.64898/2026.03.30.714220

**Authors:** Simone Webb, Antony Rose, Chuan Xu, Lloyd Steele, Masami A Kuri, Emily Stephenson, Kemal Inecik, Daniyal Jafree, April R Foster, Daniela Basurto-Lozada, Nana-Jane Chipampe, Anna V Pournara, Marc-Antoine Jacques, Ken To, Chloe Admane, Efpraxia Kritikaki, Marta A Chroscik, David Horsfall, Julia Foreman, Koen Rademaker, Julia Karjalainen, Anna Laddach, Shaista Madad, John E G Lawrence, Vitalii Kleshchevnikov, Steven Lisgo, Jimmy Tsz Hang Lee, Joséphine Blevinal, Ahlam Alqahtani, Stanislaw Makarchuk, Jake Jackson, Ekin Ucuncu, Teresa P Silva, Valentina Lorenzi, Fereshteh Torabi, Rachel A Botting, Kenny Roberts, Bayanne Olabi, Keerthi P Chakala, Leander Dony, Gabriele Dall’Aglio, Ana-Maria Cujba, Holly J Whitfield, Christine Hale, Justin Engelbert, Tong Li, Elena Prigmore, Nusayhah Hudaa Gopee, Michael W Mather, Ben Talks, Lijiang Fei, Diana Pereira, Corina Moldovan, Rasa Elmentaite, Krzysztof Polanski, Alexander V Predeus, Pasha Mazin, Vicky Rowe, Semih Bayraktar, Vincent R Knight-Schrijver, Menna Clatworthy, Kazumasa Kanemaru, James Cranley, Andrew Guo, Sanjay Sinha, David McDonald, Andrew Filby, Janet Kerwin, Peng He, Inês Sequeira, Roser Vento-Tormo, Malte D. Luecken, Alain Chédotal, Deanne Taylor, Silvia Martin-Almedina, Pia Ostergaard, Sam Behjati, Elizabeth Robertson, Nicola K Wilson, John Marioni, Helen V Firth, Caroline F Wright, Olivier Pourquie, Laure Gambardella, Paula Alexandre, Berthold Gottgens, Kerstin B Meyer, MS Vijayabaskar, Mo Lotfollahi, Sarah A Teichmann, Fabian J Theis, Fränze Progatzky, Omer Ali Bayraktar, Muzlifah Haniffa

## Abstract

Single cell genomics has enabled analysis of human prenatal development at unprecedented resolution. However, most studies have relied on dissociated tissues during restricted windows of development, limiting insights into how spatially distributed networks of cells, and multicellular niches emerge and adapt to distinct organ microenvironments *in situ.* Moreover, existing human developmental atlases have not yet been harmonised, and we thus lack a comprehensive catalogue of known cell types in the developing human body.

Here, we introduce the Human Developmental Cell Atlas (HDCA), a unified structural, cellular and molecular resource for prenatal human development. The HDCA integrates published and unpublished single cell/nucleus RNAseq atlases across prenatal organs, and includes a newly generated, spatially resolved, multimodal cell atlas of intact human embryos. Spanning 4-22 post conceptional weeks, capturing embryonic and early to mid fetal stages, the HDCA contains ∼4.6 million cells/nuclei which resolve into ∼450 cell types, explorable with a bespoke web portal.

For a global overview of the human embryo’s multicellular communities, we applied unsupervised deep learning to our intact human embryo spatial data, charting 114 tissue niches that are structural and signalling hubs for the cellular interactions of the embryo. Guided by these niches, we profiled cellular networks over space and time, not examinable using single-organ atlases. In so doing, we revealed tissue-specific fibroblast patterning from previously undescribed mesenchyme progenitors, early diversification of organ-specific blood capillaries and lymphatic vasculature, emergence of neural crest cell fates, the formation of placode-and neural crest-derived peripheral sensory neurons, and how tissue niches guide peripheral neuron maturation and axonal migration. The HDCA thus serves as a comprehensive step towards a comprehensive understanding of human prenatal development, and a template towards unravelling the biology of congenital disorders.

## Introduction

Human development is complex, spanning fertilisation, blastocyst formation, germ layer emergence, and the coordinated development and maturation of organs, all underpinned by expansive diversification of cell lineages from the initial zygote. *In vivo* studies in model organisms have uncovered key molecular mechanisms guiding developmental events, and together with advances in human stem cell biology, have furthered *in vitro* experimental culture systems that can replicate some aspects of early embryogenesis and human tissue development. However, *in vitro* culture systems and model organisms often do not fully faithfully recapitulate processes in humans. Studies of human embryos^1–4^ have provided anatomical-level understanding of development and demonstrated that congenital disorders arise from anomalies in early developmental processes. However, anatomical information is poorly integrated with high resolution multimodal cellular and molecular information, which is required to thoroughly understand how human development is orchestrated across tissues in anatomical space and gestational time.

Single cell and spatial genomic technologies have transformed our ability to profile human development at unprecedented resolution. Initiatives, such as the Human Cell Atlas (HCA) Development Bionetwork^5^, have mapped many organs, mainly across the first and second trimester of development, identifying temporal dynamics and cell types, e.g., in the brain^6–9^, gut^7,10,11^, heart^7,12–14^, yolk sac^15^, liver^7,16^, bone marrow^17^, kidney^7,16,18,19^, lung^20^, skeleton^21–23^, skin^24,25^, placenta^26^ and thymus^27^. These embryonic and fetal reference atlases have been applied to clinical research, improving our understanding of altered development^17^, the origin of childhood cancers^19,28^, and inflammatory skin diseases^25^. However, the majority of this research profiled individual organs from different donors and with bespoke sampling protocols per lab, posing a challenge for understanding cell type presence and function across the human body. Leveraging large scale computational biology and the infrastructure and principles of the HCA Bionetworks, a new wave of integrative studies have overcome this barrier within tissues (e.g., the adult lung^29^), across complex tissue networks (e.g., developmental^24^ and adult^30^ immune systems), and across the whole body ^31,32^, powering advances in rare cell characterisation, addressing inconsistencies in cell type nomenclature across labs, and even creating frameworks to identify gene programme-disease links^33^. However, integrated and cross-tissue approaches for human development remain limited due to tissue restrictions and scarcity. Consequently, major distributed tissue networks in development such as the stroma, vasculature and peripheral nervous system (PNS) remain under-explored, and given their complex architecture, would gain much from spatial profiling. Whole organism studies can address this challenge, as seen in model organisms^34–38^, but human studies are limited and focus only on very early time points^39–42^. Biological insights have thus remained largely tissue-restricted, and the timing and anatomical location of gene expression implicated in congenital disorders has remained incomplete.

Here, we present the Human Developmental Cell Atlas (HDCA, n=186, k>4.6 million cells/nuclei, 4–22PCW), a unified resource integrating community-generated, organ-focused sc/snRNAseq developmental datasets, alongside complementary single cell data of 6PCW human embryos we generated to fill a critical gap in late embryogenesis. We further produced spatial transcriptomics (Visium, Xenium) data of 6PCW human embryos, which combined with the HDCA, enabled holistic assessment of cellular dynamics across time and space and revealed key insights into: tissue microenvironment influences on early mesenchymal and fibroblast patterning, the timing of organotypic blood capillary endothelial cell specification, the establishment lymphatic vascular molecular heterogeneity, neural crest cell fates, the formation and origins of peripheral sensory neurons, and how tissue niches guide peripheral neuron maturation and axonal migration. Further, the HDCA data and models provide a foundation for future research integrating new developmental datasets, we hope even bridging the gap to the earliest implantation stages. To facilitate continued biological insights and developmental disorder research, and empower users to derive biological insights through their data, we developed an accessible webportal, Cherita, which integrates rare disease patient data from the Developmental Disorders Gene2Phenotype (DDG2P, https://www.ebi.ac.uk/gene2phenotype) dataset^43^.

## Results

### Integrated single cell and spatial omics atlas of human prenatal development

To produce the HDCA, we first compiled published sc/snRNAseq datasets spanning 4-22 post-conception weeks (PCW) and 21 developing organs (n=183 samples, k∼3.6million cells) (**Fig. 1a-b; Supplementary Table 1**). The selected datasets shared consistent sequencing technology (10x Genomics Chromium) and met minimum QC standards for cellular abundance and cell type heterogeneity (see **Methods**). Some tissues and organs were well sampled across gestation (e.g., brain), whereas others exhibited limited temporal span due to tissue sampling (e.g., mesenteric lymph nodes only sampled 16-17PCW), or being restricted to a developmental window (e.g., the yolk sac is absorbed at ∼9PCW). As there was a relative scarcity of cells at embryonic stages <8PCW, we generated complementary new data of 3 whole embryos (WE) dissected into 12 grid-like sections from late human embryogenesis at 6PCW (k∼1million cells) (**Fig. 1a-c; Fig. S1a-h; Supplementary Tables 2-4**). Our grid dissection approach for WE profiling enabled comprehensive sampling across anatomical sites and spatially-informed cell annotation, leading to a powerful resource which allows for comparison across the whole body within the same sample. scVI integration of the community developmental datasets with the WE data produced the consolidated HDCA (n=186, k∼4.6million), providing a harmonized resource that complies with HCA metadata standards (https://data.humancellatlas.org/metadata) (**Fig. S1i; Fig. S2a-c; Supplementary Table 5-8**).

**Figure 1:**
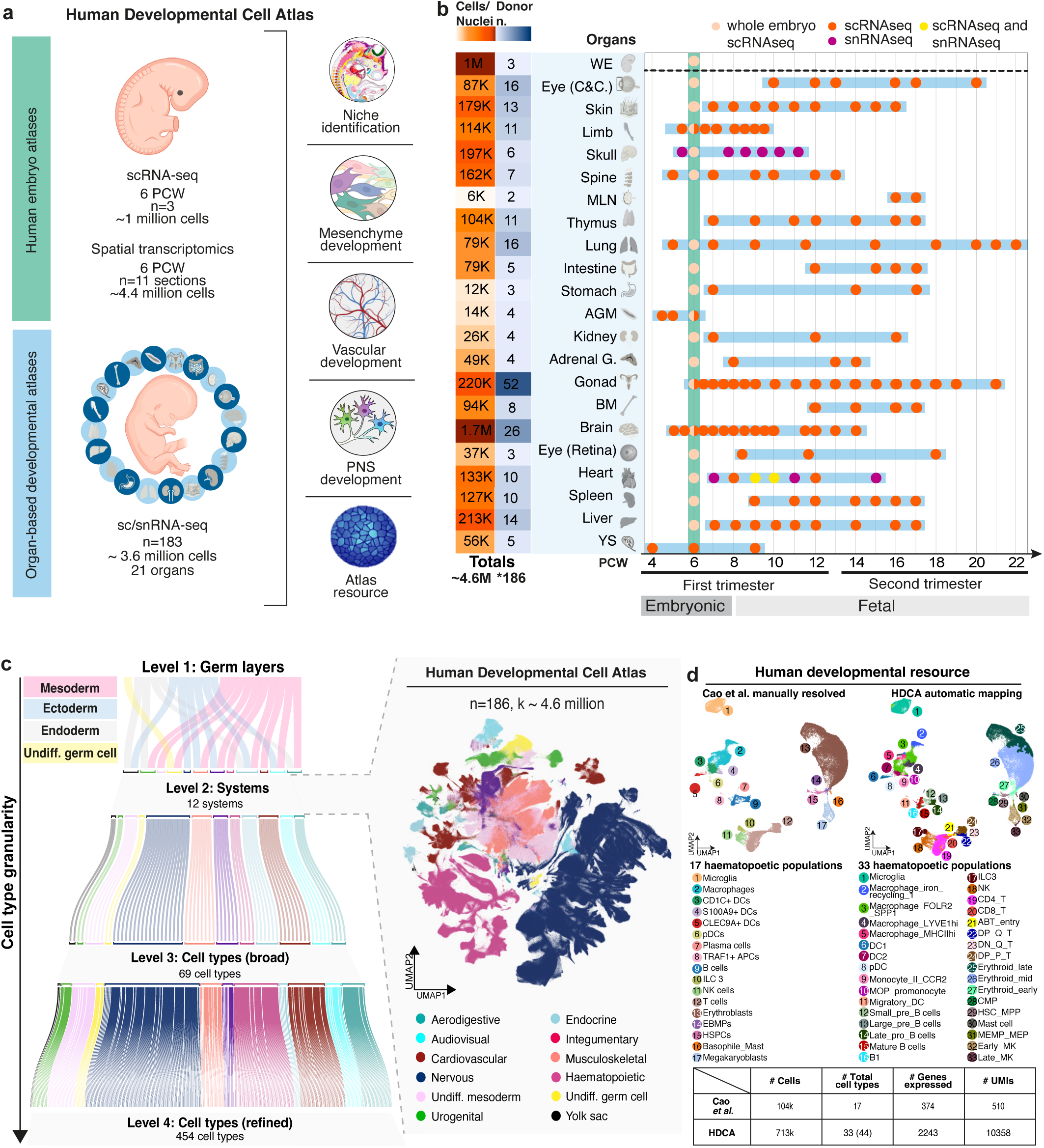
Integrated single cell and spatial omics atlas of human prenatal development. **a)** Schematic of data acquisition and overview of the Human Developmental Cell Atlas (HDCA). Top: *De novo* whole human embryo scRNA-seq data (n=3 samples; k>1million cells) and spatial transcriptomics (left: Visium, right: Xenium 5000-plex, 11 sections, k∼4.4million cells). Bottom: Community acquired sc/snRNA-seq datasets of developmental tissues (n=188; k>3.6 million; 21 organs). **b)** Graphical illustration showcasing the summary of all single cell and nuclei data included in the HDCA atlas. Dot colour indicates data chemistry and public vs *de novo* whole human embryo data. For visualisation purposes, if an age is between 2 integers, a single dot is shown. All ages are outlined in **Supplementary Table 6**. Individual organs were grouped using a hierarchical dendrogram derived from scVI of public data only (as outlined in **Supplementary Table 10**). WE = Whole Embryo, C & C. = Cornea and Conjunctiva, MLN = Mesenteric lymph node, AGM = Aorta-gonad-mesonephros, Adrenal G. = Adrenal gland, BM = Bone marrow, YS = Yolk sac. Black dotted line separates whole human embryo data from public collected data. The first column on the left hand side indicates the number of data points for each organ while the 2nd column indicates the number of tissue samples per organ. The total number of data points (k>4.6million) and individual tissue samples (n=186 total, some contribute to more than 1 organ. **c)** Left: Sankey illustration depicting the level of annotations within the HDCA data increasing in granularity going top down. Right: UMAP visualisation of HDCA data (LVL2 system, n=186; k>4.6million). **d)** UMAPs of hematopoietic cells from Cao et al^47^, colored by author-refined annotations (left) and by cell type labels transferred using the HDCA reference mapping model (right). Tables below summarize the number of cells, total annotated cell types (the number in parentheses denotes the total number of cell types present in the HDCA), and average expressed genes and UMI counts between the two datasets.

After harmonising the cell type annotation labels provided in the public data, we performed expert manual community-driven cell type annotations to: i) incorporate the newly integrated WE data, and ii) further resolve cell types with homogenous labels in organ atlases due to their not being the study focus and assigned a generic label (e.g., Mesenchyme), or undersampled (e.g., Glia) in public data (see **Methods**). This strategy resulted in 4 tiers of annotations in the final HDCA, capturing: germ layers (Level 1), 12 biological systems (Level 2), 69 broad cell types (Level 3), and 454 refined cell types (Level 4) (**Fig. 1c; Fig. S2d; Supplementary Table 9-10**). Here, refined cell types, e.g., ‘CD4 T cell’, were grouped into broad cell types which corresponded generally to lineage, e.g., ‘lymphoid’. Our systems-level annotations included: aerodigestive, audiovisual, cardiovascular, endocrine, haematopoietic, integumentary, musculoskeletal, nervous, undifferentiated germ cell, undifferentiated mesoderm, urogenital, and yolk sac (**Fig. 1c**). Cell type frequency analysis confirmed expected spatial cross-organ cell type specificity, with for example endoderm most abundant in the lung, gut and stomach; ectoderm in brain and retina; and germ cells in gonads (**Fig. S2d**). Visualising the HDCA across time showed that ectodermal neurons and glia were well-captured in the early WE 6PCW data, and mesodermal stromal cell types were present across most public organ datasets across time (**Fig. 1c; Fig. S1i**).

### Benchmarking atlas integration with trajectory fidelity

The HDCA spans 6-22 PCW and thus gives a unique opportunity to model how cellular identities transition over time. However, modelling time in single cell data is complex, and conventional benchmarking metrics such as scIB^44^ for biological conservation and batch correction, do not specifically factor for this (**Fig. S3a**). We therefore developed a custom benchmarking framework focused on assessing how well any identifiable cellular trajectories in the embedding conform to community-consensus ground truth: single cell trajectory representation metrics (scTRAM)^45^, which builds on existing scIB^44^ metrics by additionally assessing if differentiation trajectories remain conserved following integration. scTRAM evaluates trajectory fidelity along 3 axes: topology conservation (whether connectivity and branching match the reference graph), cell-order continuity (whether cells are arranged in the correct progression along directed edges) and trajectory dominance (whether the lineage manifold is preserved and not overwhelmed by batch or other nuisance structure) (**Fig. S3a**). The HDCA public data were pre-processed and integrated using commonly used tools for scRNAseq integration including scVI, scPoli, TarDis, Harmony, Scanorama and PCA, after which embeddings were assessed using both scIB and scTRAM. scVI attained the highest combined score (0.494), followed by scPoli (0.437) and TarDis (0.421), with PCA, Harmony and Scanorama performing sub-optimally (**Fig. S3b**). scVI also achieved the highest mean scTRAM score when stratified by trajectory group (**Fig. S3c**).

To assess performance differences along biologically meaningful developmental paths, we computed per-edge scTRAM scores across the three curated ground-truth lineages (germline, haematopoietic, and peripheral nervous system), and tested for significant differences between scVI and scPoli (**Fig. S3d**). scVI showed higher scores on the majority of informative edges, including late haematopoietic branches, underscoring that trajectory preservation is graded rather than binary and that integration methods introduce systematic biases at specific branch points. For the haematopoietic lineage, UMAP visualisation with scVI exhibited a continuous, correctly oriented path from HSC/MPP and LMPP/MLP through pro-B and pre-B intermediates to mature, cycling and plasma B cells, whereas the scPoli embedding showed less consistent ordering and locally misdirected flow, particularly around late pro-B and immature compartments (**Fig. S3e**). UMAP visualisation of the germline lineage also implies that the atlas yielded a more continuous progression from primordial germ cells through oogonia and pre-oocyte to oocyte and better resolved branching towards pre-spermatogonia and granulosa cells, with smoother connectivity and correctly oriented CellRank-inferred directional flow relative to the García component (**Fig. S3f**). Together, these findings support the selection of scVI for downstream analyses as it provided the strongest balance between trajectory fidelity and batch integration across the developmental atlas.

### A comprehensive, adaptable human developmental resource

To ensure the continued utility of the HDCA as a reusable reference, we provide models to the wider community for data integration (scVI), cell type label transfer (CellTypist, scArches), and niche transfer (NicheCompass) (**Fig. S4a**). We offer two complementary models for cell type label transfer: a deep learning-based scArches^46^ model that enables decentralised reference mapping through updateable, shared latent space, and a CellTypist^30^ model that performs logistic regression-based label transfer directly in gene expression space. These models leverage the fine-grained HDCA annotations to support robust cell type label transfer across different yet complementary analytical frameworks. Using the scArches model, we mapped two independent human developmental datasets spanning different developmental windows^47,48^ onto the HDCA, demonstrating its value as a reference for interpreting users’ data (**Fig. S4a**).

First, we mapped Cao et al^47^, a large human mid-gestation cell atlas (10-18PCW; ∼4.1 million cells and nuclei generated using sci-RNA-seq3) onto the HDCA reference. Cao et al^47^ described 657 cell clusters, yet the use of clustering-like nomenclature hinders systematic alignment with standardized cell ontologies. We therefore utilised their 77 higher-level groupings, which mapped to functional cell type annotations, for downstream analyses. In spite of the substantially greater sequencing depth of the HDCA, we observed robust alignment of cells across datasets within the shared latent space (**Fig. S4b**), and cell type label transfer using both scArches and CellTypist models converged on highly consistent predictions (**Fig. S5a-b**).

Concordant cell types between Cao et al^47^ and the HDCA included erythroid, myeloid, and lymphoid populations, as well as multiple cell types in the vascular, muscle, epithelial, and neural lineages (**Fig. S4b-c; Fig. S5a-b**). Within these shared groups (72 out of the 77 broad cell type groups, excluding placental lineages), the HDCA reference was able to provide increased annotation granularity for 44 cell types (**Supplementary Table 11**). Cao et al^47^ further reclustered and reannotated three major cell compartments, including hematopoietic (∼104k cells), endothelial (∼89k), and epithelial (∼282k), to achieve higher-resolution classifications. We therefore assessed whether HDCA-based automated mapping could recapitulate and potentially surpass these curated annotations. In haematopoietic populations, HDCA mapping recovered the manually defined lymphoid, myeloid, erythroid, and progenitor compartments while further resolving thymocyte states, macrophage subsets, and T and B cell subpopulations **(Fig. 1d**). Similarly, endothelial populations were partitioned beyond broad vascular and lymphatic categories into molecularly and functionally different subtypes, such as refined lymphatic endothelial subpopulations not delineated previously (**Fig. S4d**). Epithelial lineages were likewise resolved into diverse tissue-specific populations (**Fig. S4d**).

Where there was weaker cell type concordance, this generally reflected partial matching of related cell types, differences in developmental timing, or disparities in sampling of rare cell states and tissues between the query data and the HDCA reference (**Fig. S4b-c, S5a-b**). Reassuringly, cell types from tissues unique to the HDCA (e.g., yolk sac), or uniquely sampled in Cao et al^47^ (placenta) lacked reciprocal counterparts (**Fig. S4c; Supplementary Table 11**). While the HDCA reference supports mapping of datasets from comparable developmental stages, alignment to pre-implantation stages dataset^48^ were limited to pluripotent-to-endodermal axis and primordial germ cells and gonads, indicative of embryonic pluripotency programmes persistent in germline populations (**Fig. S6**). Together, these underscore the need to generate comprehensive *in vivo* references for early timepoints or to bridge this gap using *in vitro* embryoid models in the future.

### An unsupervised systematic decomposition of the human embryo into niches

The ability to identify cell types using single cell omics has provided enormous insights into human development, and the application of spatial technologies enables their *in situ* location. However, cells do not exist in isolation but form functional niches with their cellular neighbours, reflecting principles of self organisation^49,50^. To gain insight into cellular niches in tissue, we generated spatial transcriptomics on sagittal sections of an additional 6PCW whole embryo sample, including whole transcriptome (Visium) and 5,000 gene panel (Xenium) and multiplex protein imaging data (RareCyte) to map cell types and gene expression patterns across space and tissue architectures (**Fig. S1f, S7a-e; Supplementary Table 1, 3, 5**). Through collaboration with the Human Developmental Biology Resource (HDBR), macroscopic annotations of nearby haematoxylin and eosin (H&E)-stained tissue sections were incorporated into the Xenium spatial transcriptomic data, enabling multi-modal and multi-scale cross-referencing of histological features with predicted cell type assignments and gene expression profiles (**Fig. S7a**). We then performed niche detection on the Xenium data in an unsupervised manner using NicheCompass^51^, a deep learning tool which generates niche embeddings based on a cell’s gene expression and that of its local neighbourhood at single cell resolution (**Methods**).

### Histologically discernible and novel tissue niches in the human embryo

In total, we identified 114 niche clusters across 8 human embryo Xenium sections (3.1 million cells). We annotated these niche clusters based on HDBR histopathological annotations of nearby section H&Es and niche-level differentially expressed genes (DEGs) (**Fig. 2a-b; Fig. S8a-b; Supplementary Table 12**). 37 niches corresponded to focal anatomical structures, including large organs such as the heart, spine and liver, as well as smaller structures such as the mesonephros, gonadal ridge, otic vesicle, notochord, pancreas, adrenal gland and gallbladder **(Fig. 2b; Fig. S9a-b**). Among these focal structures, 32 niches corresponded to H&E annotated features (which we will refer to as ‘known’), while 5 focal niches were not distinguishable by H&E annotation (hereafter, ‘novel’) (**Fig. 2a-c; Fig. S8a-b, S9a-b)**. 30 niches corresponded to the central nervous system (CNS) (**Appendix A**). The other 47 niches featured across multiple anatomical structures, such as a *Nerve* niche and several *Vascular-rich* niches, in keeping with their presence in distributed networks (**Fig. 2a-c; Fig. S8a-b, S9a-b**). Niches also had varied presence across sagittal tissue sections, e.g., *Rathke’s pouch* and the *Gallbladder* were only captured in specific sections, highlighting the necessity of profiling multiple sagittal sections in order to comprehensively capture distinct midline and lateral embryo structures (**Fig. S8c; Supplementary Table 13**).

**Figure 2:**
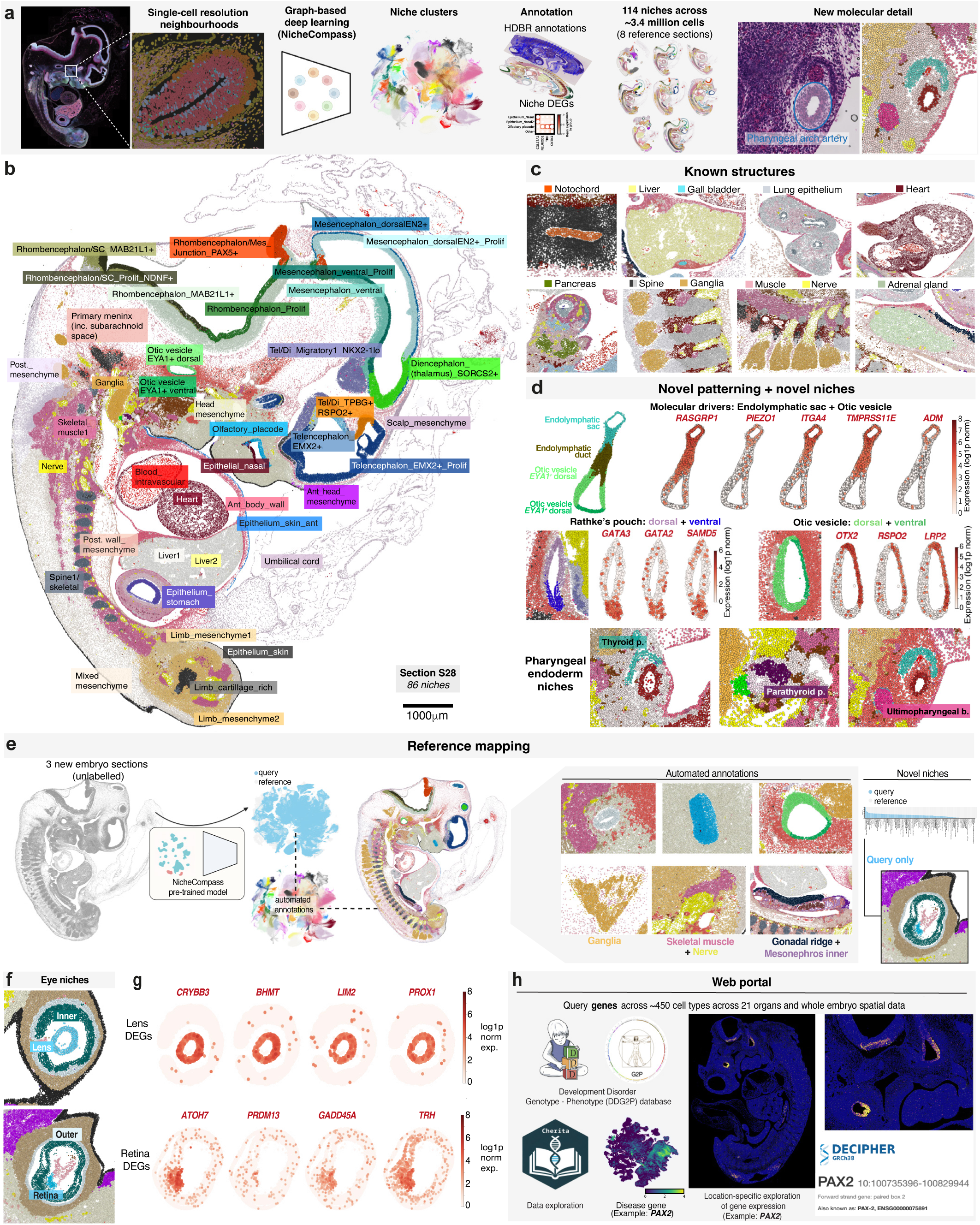
An unsupervised systematic decomposition of human embryos into niches. **a)** Overview of niche identification and annotation methodology. **b)** Xenium-5k profiling of human whole embryo section 28 (6PCW) with each cell coloured by niche assignment, with selected niche labels shown (see **Supplementary Table 13** for niche distribution across all sections).. Full Xenium NicheCompass niche labels shown in **Fig. S7c**. **c)** Focal regions of the whole embryo profiled by Xenium-5k coloured by niche across all embryo sections. We identified focal niches that corresponded to annotated structures from histology, such as the notochord and gallbladder. **d)** Top row: We captured the endolymphatic sac forming from the otic vesicle (S46 shown), including an *Endolymphatic_duct* niche. We queried differentially expressed genes between these structures. Log1p norm expression of example differentially expressed genes between the endolymphatic sac and otic vesicle. Middle row: Within focal niches that captured histological annotations, we identified dorsal-ventral patterning in Rathke’s pouch (S69 shown) and the otic vesicle, (S133 shown) allowing us to query genes driving this difference. Log1p norm expression is shown for selected genes. D: dorsal, V: Ventral. Bottom row: We identified focal niches of pharyngeal endoderm that did not correspond to annotated histological structures beyond ‘body mesenchyme’ (sections S47, S93, and S103). D: dorsal, V: Ventral. **e)** Left: schematic of methodology for obtaining niche embeddings for 3 query sections. We trained a NicheCompass model using our reference data, and then fine-tuned this model with 3 new whole embryo sections. We clustered the k-nearest neighbour graph at high-resolution and named clusters based on the highest contribution from the reference data. To identify novel niches, we queried which clusters were only present in the query (see **Methods**). Section S133 shown. Right: Results from automated annotation, identifying focal niches e.g., the olfactory placode and otic vesicle (and a novel eye retinal niche) in the query without requiring expert histological annotations. **f)** Human eye structure coloured by niche, identifying outer, inner, lens, and retinal components, from sections S36 and S203. **g)** Example DEGs for the eye lens and retinal niches. **h**) Our bespoke Cherita web portal allows for interactive exploration of the HDCA across cell types, age, and genes. Cherita integrates developmental disorder gene information from the DDG2P^43^ database (https://www.ebi.ac.uk/gene2phenotype)^192^ and DECIPHER^193^, a global platform for sharing rare disease patient data and interpreting variants, making the HDCA accessible to clinical and research communities. Information for *PAX2* can be viewed at https://www.deciphergenomics.org/gene/PAX2/expression; DECIPHER^193^ currently lists seven sequence variants in this gene.

Within the known niches, we used DEG analysis to identify candidate drivers of organ formation, confirming previous work with animal studies as well as revealing novel human markers (**Supplementary Table 12)**. We identified various known drivers described in humans, such as *NR5A1* in *Adrenal glands* and *PAX2/8* in the mesonephros and metanephros niches^52,53^, and additionally investigated structures that have been poorly studied. For instance, the endolymphatic sac, which maintains vestibular function by absorbing endolymph, has never been captured in prior whole-embryo human transcriptomic analyses, and drivers of its development are poorly understood. We identified the *Endolymphatic sac* as a distinct niche adjacent to the otic vesicle, from which it is formed, as well as a novel *Endolymphatic duct* niche (**Fig. 2d, Fig. S10a-b**). We found that *RASGRP1, ADM, ITGA4,* and *PIEZO1* were amongst the genes enriched within the endolymphatic sac (**Fig. 2d**; **Fig. S10b-d**). Profiling multiple sections allowed us to confirm these genes in a different tissue section (**Fig. S10c**).

As well as identifying key potential drivers that control the formation of tissue and organ structures, our approach also allowed us to uncover patterning within structures, including patterning of the *Rathke’s pouch* and *Otic vesicle* (**Fig. 2d; Fig. S10a**). This captured known genes, such as *OTX2* expression as a known driver for otic placode patterning^54^, as well as unreported genes including *RSPO2* and *LRP2* (**Fig. 2d**). We revealed finer resolution niche annotation for the mesonephros, demonstrating distinct glomerular (*Mesonephros_inner_glomerular*; *REN*, *FGF1*, *MAFB1*, *CLDN5*) and tubular (*Mesonephros_inner_tubule*; *PDZK1*, *BHMT*, *HNF4A*, *KCNJ15/16*) niches (**Fig. S11a-b**), and captured the *Olfactory placode* (*NEUROD1*, *RPRM*, *TRH*, *CNTN2*) within nasal epithelium in select sections (**Fig. S11c-d**). Collectively, these analyses illustrate how molecularly defined niches link established drivers of organogenesis to classical histological structures, while also uncovering previously unrecognised spatial patterning during human embryonic development.

We were also able to delineate 3 novel *EPCAM*^+^*CDH1*^+^ niches (implicated in mouse pharyngeal endoderm^34,55,56^) located around the pharyngeal artery and with distinct gene expression profiles (**Fig. 2d**), but which were only broadly annotated as body mesenchyme by H&E (**Fig. S12a**). The first niche was consistent with a thyroid progenitor, being characterised by known genes for thyroid development such as *PAX8, FOXE1,* and *NKX2-1* (**Fig. S12b**)^57^. The second niche was consistent with a parathyroid progenitor, being characterised by *PTH* (parathyroid hormone) expression and genes required for parathyroid and/or thymus development, such as *GATA3*, *MAFB, SIX1* and *EYA1*^5859^. Of note, the final pharyngeal endoderm niche represented the transient ultimopharyngeal body (or ultimobranchial body) that gives rise to the calcitonin-producing neuroendocrine cells (C-cells) of the thyroid^60,61^. This niche arises from the 4th pharyngeal pouch in mammals and was present in a similar location to the parathyroid progenitor niche (**Fig. 2c**). C-cells have been profiled in human development but only from 9PCW^62^. Our high-plex profiling of this niche revealed a distinct transcriptomic profile for early embryonic C-cells compared to later timepoints (**Fig. S12b-d)**. This niche expressed genes associated with immature embryonic C-cells in mice (*ISL1*^63^, *FOXA1* and *FOXA2*^61^, *ASCL1*^60^, *NKX2-1*^64^) and other genes associated with neuroendocrine differentiation (*HHEX*, *ENO2, DLL3*)^65^. As expected, expression of more mature C-cell markers was low or absent. For instance, *CALCA*, encoding the alpha isoform for calcitonin, was relatively lowly expressed in this niche, whereas *CALCB*, encoding the beta isoform, was highly expressed (**Fig. S12b-d)**. We thus provide the first detailed characterization of early pharyngeal endoderm niches in human development with spatial resolution, including the previously uncharacterised ultimopharyngeal body. Moreover, this demonstrates that our niche-based approach can resolve distinct molecular structures within tissues that are histologically similar.

Our NicheCompass approach incorporates anatomical information within niche labels, allowing for future community automatic niche transfer and cell type assignment to new embryo tissue sections without domain or histological expertise, even with a small number of query tissue sections. To demonstrate this approach, we mapped a further 3 embryo sections to our core 8-section dataset using reference-query mapping, identifying 90 niches in these new sections (**Fig. 2e; Fig. S13a-d; Methods**). Notably, we identified a retinal niche (*Eye3_innermost_retina*) not present in the reference dataset (**Fig. 2f; Fig. S13**), as well as an *NTN1^+^WNT3^+^GABRB2^+^* niche present within a focal region of *MEIS1^+^HMX2^+^* otic vesicle (**Fig. S10a, c, S13c**). *HMX2* is a marker of dorsolateral otic epithelium^66^, and the focal *NTN1^+^WNT3^+^GABRB2^+^* niche expressed genes that have been associated with semicircular canal development from the dorsolateral otic vesicle in animal models (*WNT3*, *NR4A3*, *NTN1*)^67,68^ (**Fig. S13c**), illustrating the granularity of our niche capture. The human retina has not been studied in early embryogenesis, with prior transcriptomic studies performed at later timepoints (8-9PCW) or in organoid models^69,70^. We showed that the embryonic retina was characterised by expression of *ATOH7*, required for retinal ganglion cell (RGC) development^71^*; PRDM13*, which specifies amacrine cell fate in mouse^72^; *GADD45A*, which characterises early neurogenic cells in mouse retina^73^; and *TRH*, which encodes thyrotropin releasing hormone (**Fig. 2f-g; Fig. S13d**). *TRH* signalling has been reported to specify cone subtypes in human retinal organoids^74^, and thus we provide further evidence in support of a role for *TRH* in the retina.

We finally wanted to confirm whether our niches would be informative for understanding the role of genes implicated in developmental disorders. To illustrate the utility of niches captured, and after identifying the retinal niche above, we investigated reported genes for congenital cataracts^75^ (**Methods**), which is characterised by lens opacity. Consistent with this pathology, congenital cataract genes were enriched specifically in the *Ey*e*1*_*lens* niche (**Fig. 2f-g; Fig. S13e**), including genes encoding crystallins (*CRYAA, CRYBB3, CRYGS, CRYGC*), transcription factors (*MAF, FOXE3*), cytoskeletal proteins (*BFSP1*), and RNA processing proteins (*TDRD7*)^75^. *TDRD7* was initially validated in developing mouse lens^76^, but we illustrate how this gene can now be easily interrogated in the context of human development through query of expression across multiple embryo niches, demonstrating the potential value of our resource for querying disease-associated genes given our comprehensive sagittal profiling and niche capture. Overall, we illustrate how our data can serve as a resource for human embryo biology across diverse niches and generate histologically-informed niche annotations for novel data.

In addition to providing light-weight access to the HDCA through the models described above, our bespoke web portal, Cherita (https://cellatlas.io/studies/hdca/; private until publication), allows interactive exploration of both droplet-based and embryo spatial datasets without requiring computational resource or technical expertise (**Fig. 2h**). Cherita incorporates data from the DDG2P dataset (https://www.ebi.ac.uk/gene2phenotype)^43^, which shares detailed, evidence-based gene-disease models curated from the literature. Given that our HDCA data covers 21 organs and the whole embryo, queried genes can be investigated across different organs and developmental stages to contextualize spatiotemporal patterns. For instance, loss-of-function variants in *PAX2* are known to cause Renal-Coloboma Syndrome [OMIM #120330]^77^, characterised by optic nerve abnormalities, hypodysplastic kidneys, and, in some cases, high-frequency sensorineural hearing loss. Consistent with this phenotype spectrum, we observed that *PAX2* was highly expressed in the eye, metanephros, and otic vesicle niches (**Fig. 2h; Fig. S13f**). We thus provide the first platform for tissue niche-level queries of human developmental tissues to dissect expression profiles across structures, and for comprehensive investigation into developmental disorders linked to pathogenic genetic variants.

### Mesenchymal progenitors in the human embryo

Leveraging the HDCA cross-temporal data and spatial niche identification, we then interrogated the major distributed tissue networks in development which have, to date, have not been investigated in a cross-tissue manner across human gestation: the developing mesenchyme, vasculature, and peripheral nervous system.

Within the developing mesenchyme, current understanding suggests that common fibroblast progenitors arise from the mesoderm and neural crest and contribute to anatomical region-specific fibroblast subsets^78^. Single cell transcriptomics technologies have elucidated novel heterogeneity in adult fibroblasts across tissues, which previously was challenging to resolve due to unreliable surface markers^79^. However, fibroblast diversity within the early stroma of the embryo remains poorly understood as fibroblasts have typically been studied in an organ-specific manner. For instance, *HIC1^+^* and *FRZB^+^* fibroblast progenitors have been described in the skeletal system and skin, respectively^22,80^. Despite recent whole organism profiling efforts^39–42,81,^ the relatively small number of cells and single cell resolution spatial data means it remains unknown how these different progenitor cells compare to each other and whether cross-organ populations.

From a joint integration of ∼3.6 million predominantly mesenchymal cells (**Methods**) from both scRNA-seq and Xenium data, we defined 108 cell types within 12 broad groups (**Fig. 3a**; **Fig. S14-16; Appendix B**). Ten clusters were present at early timepoints (**Fig. 3b)** and remained undefined after expert annotation (**Methods**), raising the possibility that they represented undefined progenitor populations. These clusters expressed *FRZB^+^* +-*HIC1^+^*, and other dermal fibroblast progenitor markers (**Fig. 3c**), indicating broad expression of these previously reported genes for organ-specific progenitors. We then defined the spatial distributions and niche residences of these 10 *FRZB^+^*+-*HIC1^+^* fibroblast progenitor subsets across tissue sections (**Fig. 3d; Fig. S14d, S17a**). The *PDGFRA^+^* progenitor population was non-specific and broadly distributed. The remaining populations demonstrated more focal distributions, including enrichment in the skeletal system (*RUNX2^+^*), limb (*IRX3^+^ISL1^+^*), gut and anterior body (*FOXF1^+^*), and head/jaw and limb (*SATB2^+^PAX3^+^* and *SATB2^+^DCX^+^*). We also delineated 4 *ZIC*^hi^ progenitor populations enriched in the scalp (*ZIC2^+^PCSK9^+^*), primary meninx (*ZIC2^+^ANGPT2^+^*, *ZIC2^+^WNT4^+^*), and the spine/posterior vertebral wall (*ZIC2^+^FIBIN^+^*) (**Fig. 3c-d; Fig. S14d**). We corroborated these distinct locations through interrogation of the anatomical origin of scRNA-seq data (**Fig. S17b**) and PAGA analysis (**Fig. S17c**). These progenitor clusters also showed distinct gene expression signatures that were consistent in scRNA-seq and Xenium data (**Fig. S14c**). Our analysis thus reveals novel regionally-patterned fibroblast progenitor populations that broadly express previously used markers for organ-specific progenitors (*FRZB*, *HIC1*) but which can be distinguished by other marker genes, redefining the stromal landscape in early embryos.

**Figure 3:**
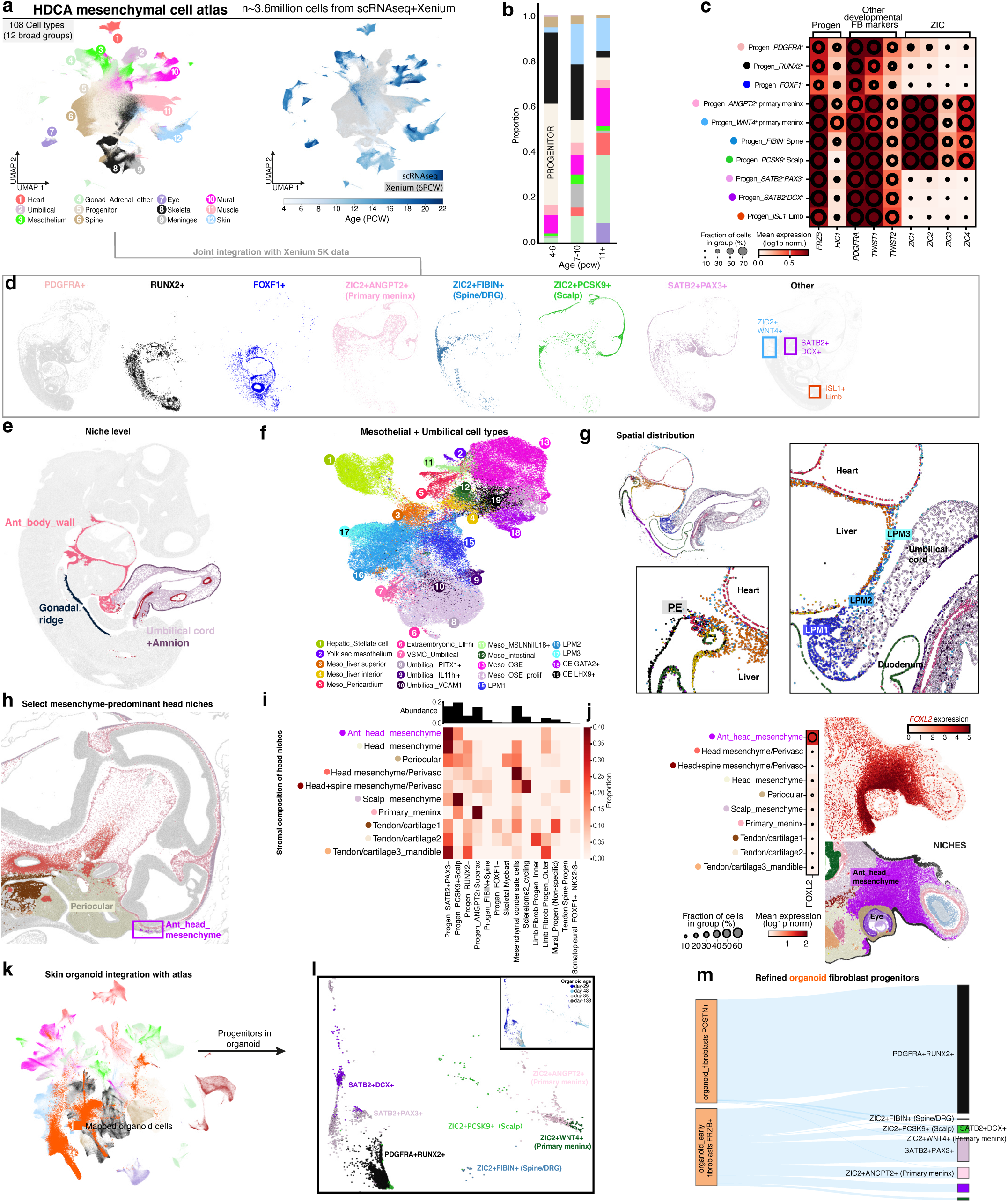
Mesenchymal progenitor heterogeneity in the human embryo. **a)** Left: UMAP embedding of predominantly mesenchymal clusters including scRNA-seq and Xenium-5k data, coloured by their broad cell type group. Across these broad groups we identified 108 cell types. Right: UMAP embedding coloured by age of sample. Xenium-5k samples (all 6 PCW) coloured in grey. PCW: post-conception weeks. **b)** Proportion of cells for each time grouping from each broad group, showing enrichment of progenitors in the early samples. scRNA-seq data only. **c)** Dotplot of expression of broad progenitor markers and *ZIC* genes in undefined human fibroblast progenitors in scRNA-seq data. **d)** Xenium-5k of whole embryo coloured by the undefined progenitor populations, showing distinct localisations (section S28, far right column of rarer populations in S47). **e)** Xenium-5k of whole embryo coloured by niches where mesothelial and umbilical cell populations are predominantly located (S46). **f)** UMAP embedding of mesothelial and umbilical cells coloured by cell type assignment. Hepatic stellate cells were located within the mesothelial group, consistent with a reported origin from septum transversum-derived mesothelium in mice. **g)** Location of mesothelial and umbilical populations in a Xenium-5k whole embryo section (S46). 2 broad clusters of mesothelial cells represented the LPM and PE. The LPM1 population was enriched in the anterior body mesenchyme lining, and LPM2 and LPM3 showed migration from this site. All LPM locations shown in **Fig. S18a**. PE: proepicardial organ. LPM: lateral plate mesoderm. **h)** Xenium-5k of whole embryo head coloured by niches with predominantly mesenchymal cells (S28). Other niches in this region (central nervous system) coloured in grey. **i)** Heatmap showing distribution of fibroblast subtypes by head mesenchymal niche across all Xenium 5k reference sections. **j)** Combined heatmap-dotplot of *FOXL2* expression by head mesenchymal niche. All niches shown in **Fig. S7c**. Example of lateral embryo section coloured by *FOXL2* expression (top), showing enrichment around the eye, and coloured by niche (S193; bottom). **k)** UMAP embedding of organoid cells (orange) compared to mesenchymal atlas coloured by broad cell type group. **l)** UMAP embedding of human organoid cells within the progenitor cluster, showing identification of Progen_SATB2+PAX3+, Progen_ZIC2+PCSK9+(Scalp), Progen_ZIC2+FIBIN+(Spine/DRG), and Progen_ZIC2+ANGPT2+ populations. Dotplots of these progenitor populations in the skin organoid using markers compared to human populations is shown in **Fig. S20b**. **m)** Sankey plot of re-assignment of original organoid labels, showing identification of fine fibroblast populations. Cell populations with a fraction of 0.02 or more are shown.

### Lateral plate mesoderm heterogeneity and mesothelium formation

Embryonic internal organs are embedded within coelomic cavities which are lined by epithelial monolayers termed mesothelial cells, which arise from the lateral plate mesoderm (LPM)^82^. Notably, original author annotations of the HDCA community data had not resolved LPM cells (**Fig. S15a-b**), possibly reflecting the relatively small number of LPM captured when studied in an organ-specific manner, or that mesothelial populations were discarded or lost during tissue dissection. Our spatial and integrative approach captured coelomic cavities in the embryonic anterior body wall niche, including 3 LPM subtypes enriched in anterior body mesenchyme (**Fig. 3e-f; Fig. S18a-c**). The *SFRP5^+^* LPM1 population was enriched in the anterior body mesenchyme, particularly the inner lining of this structure (**Fig. 3g; Fig. S18a-b)**. Notably, *SFRP5* has been reported to be a marker gene for an LPM progenitor in zebrafish^83^. The other 2 populations (LPM2, LPM3) showed a more extensive distribution, with the *DKK1^+^NRN1^+^* LPM3 population particularly observed in the hepatic and cardiac regions (**Fig. 3g; Fig. S18a-b**), consistent with the LPM as a source of mesothelial populations at these sites. In addition, we also observed a cluster of mesothelial cells located posterior to the heart (**Fig. 3g**), consistent with the proepicardial organ.

Organ-specific mesothelium populations have been reported in prenatal lung^20^, heart^14^, liver^84^, and intestine^85^, but similarly to fibroblast progenitors, it is unclear how these different populations are transcriptionally similar or related, or the extent of their organ-specificity. Highlighting the power of our atlas to resolve populations with fine (monolayer) distributions, we captured distinct mesothelial cell populations lining the heart, intestine and inferior and superior liver. We also observed coelomic epithelium subtypes lining the gonadal ridge and mesonephros **(Fig. 3e-g; Fig. S15d, S18c**). As we captured these distinct mesothelial populations from across different organs, we were able to determine which previously reported mesothelial markers for distinct organs were specific (e.g. *CALB2*, *SBSPON*)^86^ and which were shared (e.g., *TNNI*, *BNC1*, *PODXL, WT1*)^14,20^ **Fig. S15c**). Additionally, we defined more comprehensive gene signatures to distinguish these developmental mesothelial subsets (**Fig. S15c**). We thus comprehensively decode the classical LPM structure with single cell spatial resolution, revealing novel LPM subtypes with distinct spatial distributions, and profile the distinct mesothelial populations that they give rise to across organs.

### Extra-embryonic umbilical cord and mesenchymal cell heterogeneity

Prior whole embryo analyses have not resolved extraembryonic mesenchymal populations^40,42^, or have grouped these populations as a single cluster^41^. Here, we revealed novel heterogeneity in the extraembryonic tissue, capturing 4 different umbilical populations and an additional 2 extraembryonic mesenchymal populations (**Fig. 3e**; **Fig. S16a-d)**. A *PITX1^+^* population was the predominant population in the umbilical cord with a broad distribution, with the remaining populations showing more focal distributions. The *SATB2^+^IL11^hi^*population was identified at the periphery of the umbilical cord. The *PITX1^+^VCAM1^+^*population was enriched at the attachment to the embryo. We also profiled umbilical vascular smooth muscle cells (*Umbilical_VSMC*) around the umbilical cord blood vessels (**Fig. S16c-d**), that have been ill-defined to date^87^. The 2 extraembryonic mesenchymal populations, *Extraembryonic_CLIC3^hi^* and *Extraembryonic_LIF1^hi^*, were at the periphery of the umbilical cord (**Fig. S16d**). These extraembryonic populations expressed genes encoding *AFB* and *ALB*, associated with placental cells, but not trophoblast markers (**Fig. S16e**). We thus capture multiple novel cell populations present within extraembryonic tissue in the late embryo, including poorly defined supporting cells around umbilical vessels.

### Decoding cranial embryo niches

Elements of the vertebrate head arise from both mesodermal and neural crest sources^88^. Recent spot-level spatial transcriptomic work has suggested distinct fibroblast subtypes in the head, but this study was limited to only a focal region^40^. We identified 10 head niches formed predominantly of mesenchymal cells (**Fig. 3h; Fig. S19a**). These mesenchymal head niches contained multiple fibroblast subtypes (**Fig. 3i**). In addition to the non-specific *PDGFRA^+^* population, the most abundant progenitor subtypes in the head were the *ZIC2^+^PCSK9^+^* scalp, *ZIC2^+^ANGPT2^+^* primary meninx, and *SATB2^+^PAX3^+^* populations.

Notably, the *Ant_head_mesenchyme* niche showed a distinctive focal location in the anterior face, and was markedly more abundant in association with the periocular niche (**Fig 3h; Fig. S19b**). The *Ant_head_mesenchyme* niche predominantly contained *SATB2^+^PAX3^+^*and *ZIC2^+^PCSK9^+^* fibroblast progenitors (**Fig 3i**), which were not unique to this niche. We therefore queried whether these fibroblasts showed distinct gene expression in different niches. *FOXL2* expression was highly enriched in the *Ant_head_mesenchyme* niche compared to other head niches (**Fig. 3j; Fig. S19c**).We therefore queried *FOXL2* expression within *SATB2^+^PAX3^+^* and *ZIC2^+^PCSK9^+^* fibroblast progenitors across different niches. We observed increased *FOXL2* expression with these progenitors in the *Ant_head_mesenchyme* niche specifically (**Fig. S19d**), suggesting that these fibroblasts exhibit distinct gene expression across different niches. Interestingly, *FOXL2* is implicated in eyelid formation, with loss-of-function mutations causing eyelid abnormalities in blepharophimosis-ptosis-epicanthus inversus syndrome (BPES), in agreement with the enrichment of the *Ant_head_mesenchyme* niche around the *Periocular* niche. Our analysis thus suggests a distinct niche involved in the formation of the eyelid structure that is not identifiable from cell type composition alone.

Overall, within the developing mesenchyme, we reveal novel complexity in fibroblast progenitors across the embryo; comprehensively profile extraembryonic and mesothelial populations at single cell spatial resolution, including migrating LPM; and identify and decode head mesoderm niches, revealing niches contributing to the formation of specific structures, providing the most complete profiling of human developmental fibroblasts to date.

### Novel mesenchymal progenitors are recapitulated in human organoid model

We aimed to determine if our newly-defined human fibroblast progenitor populations were captured and corroborated by an induced pluripotent stem cell (iPS)-derived organoid model containing mesenchymal lineage cells. To this end, we mapped published human iPS-derived skin organoid scRNA-seq data^89^ to our HDCA (**Methods**). We selected human skin organoid as this organoid closely mimics prenatal skin structural composition^80^ and has a proposed anatomical origin (craniofacial region derived from neural crest differentiation)^89^. We observed that skin organoid fibroblasts largely mapped to 3 major mesenchymal groups: progenitors, skin, and skeletal (**Fig. 3k-l; Fig. S20a**). The skeletal population is consistent with a known off-target cartilage population of human skin organoid. Within the progenitor group we identified 4 *ZIC*^hi^ populations (*ZIC2^+^PCSK9^+^_scalp*, *ZIC2^+^ANGPT^+^_primary_meninx*, *ZIC2^+^WNT4^+^_primary_meninx*, *ZIC2^+^FIBIN^+^_spine)* (**Fig. 3l-m**). We had earlier identified these *ZIC*^hi^ progenitor subtypes in the embryo head and spine region (**Fig. 3d, i**), and *ZIC* proteins are linked to neural crest induction as well as meningeal fibroblast precursor identity^90,91^. We also identified *RUNX2^+^* progenitors, linked to the development of skeletal fibroblast populations, and *SATB1^+^* progenitors. These organoid progenitor populations expressed similar gene signatures to the equivalent human embryo populations, with the exception of *SATB1^+^* progenitors (**Fig. S20b)**. The presence of our newly defined progenitors within the human skin organoid model provides further validation of these populations consistent with their anatomic location in human embryos. Finally, the cellular census powered by the HDCA allows resolution of fine cell subtypes within diverse *in vitro* systems. For instance, within the off-target cartilage population, which previously has been broadly labelled as *POSTN^+^* fibroblasts only, we identified *GDF5*-expressing *Interzone cells*, which characterize the initial mesenchymal condensation of synovial joints and give rise to chondrocytes^22^, as well as *Chondrocytes* themselves, *Tenocytes*, *Skeletal myocytes*, *Osteoblasts*, and *Perichondrium* (**Fig. S20c-d**), illustrating the value of our resource to decode unrecognised stromal heterogeneity in organoid models. Overall, we illustrate how our spatially-resolved profiles of developmental human stroma, defined across millions of cells, can be used to define novel complexity in fibroblast progenitors in human organoid models and resolve fine heterogeneity through reference-query mapping. Our HDCA atlas and its associated web portal can thus be used as a powerful resource for interpreting future organoid and organ-specific datasets as well as guiding organoid differentiation.

### Organ-specific blood endothelial specialisation in development

The second distributed system we explored was the nascent vascular system. The vascular endothelium consists of a layer of endothelial cells (EC) lining all vessels, exhibiting remarkable molecular and functional diversity and adapting to the specific demands and microenvironments of distinct tissues and organs^92,93^. Single cell atlases have been used to profile molecular identity across the adult human vasculature, revealing the transcriptional state of arteries and veins in addition to organ-specialised capillary programs underpinning endothelial function and phenotype^94–96^. Studies have also revealed that EC molecular heterogeneity emerges during embryogenesis and is shaped by local tissue cues in mice^97,98^. Developmental EC data in humans however is restricted to individual fetal organ studies^99–101^. Here, we leveraged the HDCA to isolate and analyse 94,681 ECs and provide the first systematic, multi-organ, and temporally resolved atlas of EC states across human gestation.

Unsupervised clustering of the HDCA ECs revealed 37 transcriptionally distinct cell types (**Fig. 4a**). To annotate these subsets, we curated canonical marker genes from an adult human vascular atlas^94^, and cross-referenced them with transcriptional signatures and structured metadata from the public datasets within the HDCA (**Fig. S21a**). All major endothelial compartments were represented, including arterial, venous, capillary, and lymphatic ECs (**Fig. S21b**). Consistent with adult profiling^94^, arterial and venous ECs largely formed organ-agnostic clusters (**Fig. 4b**), with central nervous system and spleen harbouring distinct arteriolar populations from the canonical common arterial EC cluster shared between organs. Alongside a large capillary EC cluster shared by multiple organs (**Fig. S21c**), certain organs, including the heart, lung, central nervous system, and liver, possessed transcriptionally distinct capillary EC subsets, reflecting their fully differentiated adult counterparts^94^.

**Figure 4:**
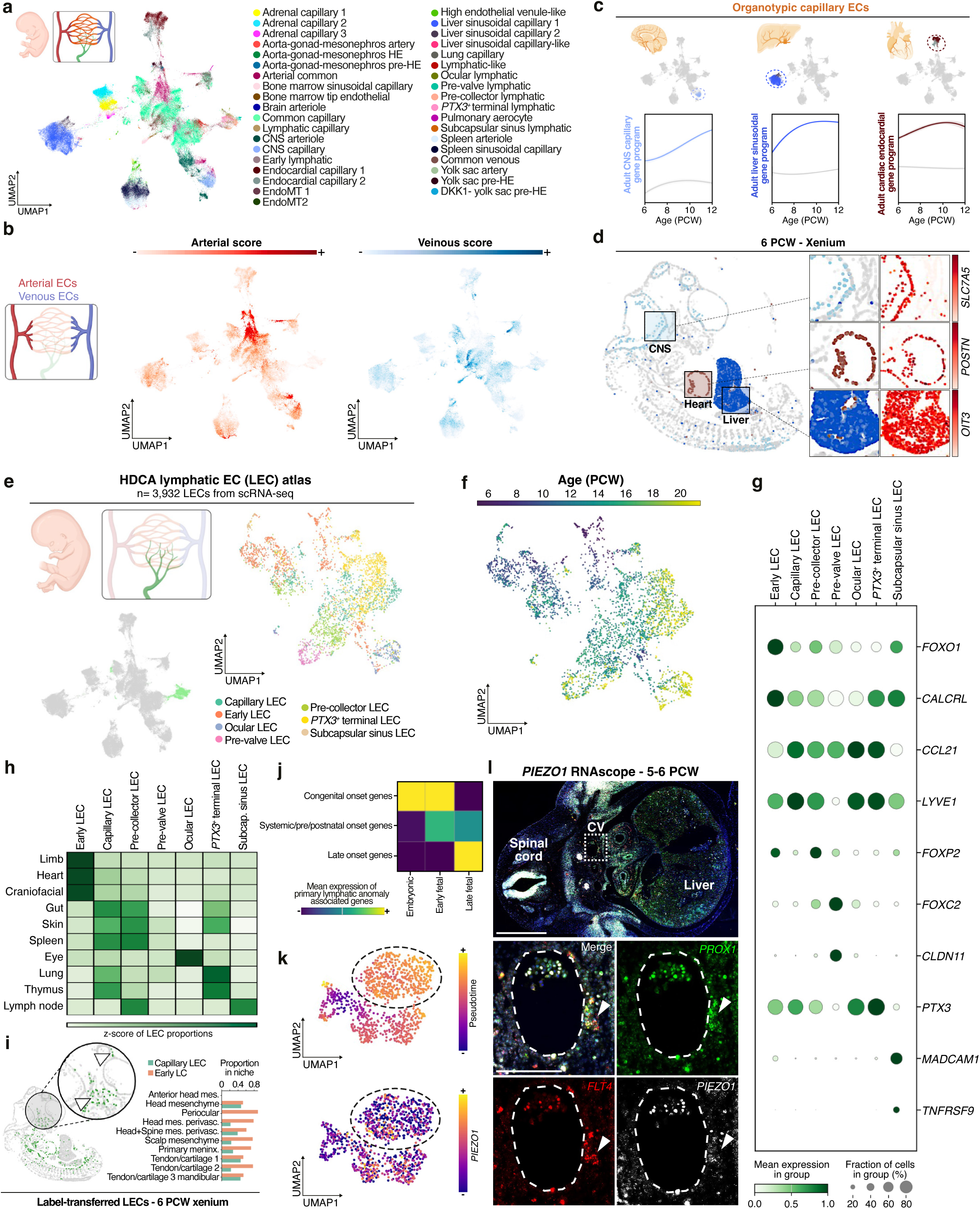
Spatiotemporal analysis of blood vascular and lymphatic endothelial heterogeneity during human development. **a)** UMAP visualisation of HDCA endothelial cells (EC, n=94,681 cells) generated using scRNA-seq, resolving 37 transcriptionally distinct EC types. **b)** Feature plots showing arterial (above, red) and venous (below, blue) gene signatures on the EC UMAP. **c)** Generative additive model (GAM) analysis of organ-specific capillary subsets in the heart (left), liver (middle) and central nervous system (CNS, right), with above UMAPs depicting endocardial cells, liver sinusoidal endothelial cells and CNS capillaries, respectively. Adult gene programs sourced from previously published data were plotted against time in these EC populations (coloured lines) and plotted against randomly sampled ECs from the atlas (grey lines). The faded edge of each line represents the 99% confidence interval. **d)** ECs were isolated from Xenium data, and label transfer and analysis of marker genes was performed. Label transfer from scRNA-seq to Xenium lead to successful labelling of ECs in the CNS (top), heart (middle) and liver (bottom) with corresponding organ-specific capillary-enriched marker genes shown to the right of each subpanel. Section S133 is shown and is representative of three sections analysed. **e)** UMAP visualisation of HDCA lymphatic ECs (LEC, n=3,932 cells), subclustered to resolve 7 transcriptionally distinct LEC types. **f)** UMAP of LECs are plotted here by developmental stage. **g)** Dotplot showing canonical marker genes for each LEC subtype. **h)** Matrix plot with the colour representing z scores for percentage of LEC subtypes across organs. **i)** Label transfer from scRNA-seq to Xenium demonstrates lymphatics to appear in the cranial region from multiple serial sections of a human embryo at ∼6PCW, shown in section S133 representative of three sections analysed. **j)** Calculation of scores for genes associated with primary lymphatic anomalies with congenital onset, systematic / pre- or postnatal onset and late onset, with scores shown over embryonic, early fetal and late fetal stages within LECs of the HDCA. **k)** Pseudotime analysis of early LECs (top, n=995 cells) and featureplot showing *PIEZO1* expression across the trajectory (bottom). Dotted circle indicates early pseudotime. **l)** RNAScope for *PIEZO1* of axial sections of a 5-6 PCW human embryo, alongside the lymphatic markers *PROX1* and *VEGFR3*. The top panel shows low magnification, with the bottom panels showing the boxed region magnified. The white arrowhead shows the co-expression of *PIEZO1, PROX1* and *VEGFR3* in cells within the wall of and adjacent to the cardinal vein (CV). Scale bars 200μm.

To more precisely resolve the timing of EC specialisation, we computed organ-specific gene programs (GPs) using differentially expressed genes from scRNA-seq data of adult ECs across multiple organs^94^. For each developing organ, we generated a subset of the ECs and computed GP scores corresponding to their adult counterparts, before plotting these GPs across developmental time using generalised additive models (GAM) (**Fig. 4c**). In brain, heart and liver, organs with transcriptionally distinct capillary ECs, adult-like gene programs were already present as early as 6PCW. In order to validate this finding, we integrated our suspension data with 163,973 ECs sampled across 11 Xenium sections of the 6PCW human embryo (**Fig. S21d**), and performed label transfer from our HDCA EC atlas to Xenium data (**Fig. 4d**). We additionally detected the expression of canonical organotypic capillary-enriched genes within the Xenium data, including the blood brain barrier endothelial markers *SLC7A5*^+^, which was expressed in ECs lining the maturing CNS^102^, ECs within the endocardium expressing the endocardial marker *POSTN*^+^^103^, and ECs within the embryonic liver which were enriched for the liver sinusoidal endothelial marker, *OIT3*^104^. In contrast, shared capillary ECs exhibited diffuse and organ-agnostic distribution (**Fig. S21e**). These results suggest that organ-specific endothelial transcriptional programs are established during early organogenesis, and that molecular hallmarks of specialised adult ECs are detectable during the first trimester.

### Lymphatic endothelial heterogeneity during human development

We next characterised the lymphatic EC (LEC) compartment, identifying a distinct rare cluster of 3,932 LECs within the HDCA, identified by their expression of canonical lymphatic endothelial markers including *PROX1, FLT4/VEGFR3,* and *PDPN* (**Fig. S22a**). The lymphatic system in adults is highly morphologically specialised^105–108^, and emerging evidence shows LECs possess organ-specific structural, molecular and functional features^109–111^. While scRNAseq has revealed insights into lymphatic development in model organisms^112,113^, studies in human development have been restricted to individual organs^114–116^, and a systematic analysis of the profile and timing of LEC specialisation during human development is required. Unsupervised clustering of LECs revealed 7 transcriptionally distinct subsets (**Fig. 4e**). One cluster was enriched in cells from early embryonic stages (6–10PCW) (**Fig. 4f**), which we termed early LECs. Marker gene analysis showed early LECs expressed *FOXO1* and *CALCRL* (**Fig. 4g**), genes essential for the regulation of lymphatic development in mouse models^117–121^. Conversely, the remaining 6 clusters, containing cells from later developmental stages, showed transcriptional similarities with known LEC subsets in adulthood, including *LYVE1^+^CCL21^+^*capillary^122,123^, *FOXP2^+^*pre-collecting^124^ and *FOXC2^+^CLDN11^+^* pre-valve^125,126^ LECs, ocular LECs^127^, *PTX3^+^* terminal lymphatics^128,129^ and *MADCAM1^+^TNFRSF9^+^* LECs associated with the lymph node subcapsular sinus^109,130^. To investigate organ-specific patterns, we quantified LEC subtype abundance across tissues (**Fig. 4h**). Capillary and pre-collecting LECs were broadly distributed, while ocular and subcapsular sinus LECs were spatially restricted to the eye or lymph node, respectively. *PTX3^+^* terminal LECs, known to interact with innate immune cell subsets^128,129^, were enriched in thymus and mucosal or barrier organs, including gut, skin, and lung. Intriguingly, early LECs were most abundant in the limb buds, heart, and craniofacial regions. Whereas heart lymphatic development has been profiled in mouse studies^131^, and cardiac progenitors with potential for lymphatic differentiation have been identified using lineage tracing experiments^132–134^, the existence of LECs within the human developing cranium contrasts with prior mouse literature. These studies suggest that lymphatics in this region, such as those of the meninges and ocular regions, emerge postnatally in mouse^135–137^. We corroborated these findings with our Xenium data, within which label transferred LECs were detected within the head region (**Fig. 4i**). Leveraging niche annotations to more carefully resolve their anatomical location, capillary LECs and early LECs were detectable in 9 out of 10 head niches, including the primary meninx, the primitive layer of mesenchyme giving rise to the meninges, where central nervous system lymphatics have been reported in human adults^138–142^.

### Spatiotemporal expression of genes associated with primary lymphatic disorders

When genes essential for lymphatic development and function are disrupted, the result is a spectrum of disorders leading to the build-up of interstitial fluid within the tissue extracellular space^143^. Collectively, these are known as primary lymphatic anomalies (PLA)^144^. To assess the relevance of our analysis of human developing LECs to PLA, we profiled expression of nine genes implicated in PLAs, commonly used in clinical variant screening panels (**Fig. S22b; Supplementary Table 14**). Gene expression was stratified by developmental age and grouped into embryonic, early fetal and late fetal stages, with mean expression scores calculated for genes associated with congenital, pre- or postnatal, and late-onset (> 1 year after birth) forms of PLA (**Fig. 4j**)^144^. Genes associated with congenital and early onset PLA were preferentially enriched at embryonic stages, whereas genes linked to late onset PLA were enriched at fetal stages. This temporal stratification is concordant with clinical observations^143^, and suggests that early-onset disorders arise from variants in genes associated with disruption of lymphatic specification, whereas late-onset disease-associated genes may act during later stages of lymphatic maturation.

One gene of particular interest was PIEZO1, which is associated with generalised lymphatic dysplasia, an early onset PLA^145,146^. PIEZO1 encodes a mechanosensitive ion channel, and although biallelic loss of function variants in *PIEZO1* have been associated with fetal hydrops^145,146^, its also implicated in postnatal lymphatic valve formation and maturation, a later phenomenon and currently the most studied role of PIEZO1 in the lymphatic system^147,148^. To investigate this further, we examined PIEZO1 expression in *FOXO1^+^ CALCRL^+^* early LECs. Pseudotime analysis predicted *PIEZO1* expression even at the earliest stages of the trajectory for these early LECs (**Fig. 4k**). As vasculature development is poorly recapitulated in human organoid models^80^, we corroborated our findings in model organisms used to profile lymphatic development. We examined recently published single cell data of LECs and associated ECs at the onset of lymphatic emergence in mice^112^ and zebrafish^113^ embryos (**Fig. S22c-f**). Using the original author cell type annotations, we found that ECs committed to lymphatic endothelial fate were enriched in the expression of the murine and zebrafish orthologs of *PIEZO1*, alongside *FOXO1* and *CALCRL*, markers used to identify early LECs within the HDCA. To validate this finding within a human context, we performed RNAscope in 5–6PCW embryos, demonstrating expression of *PIEZO1* in cells co-expressing the pan-LEC markers, *PROX1* and *FLT4/VEGFR3*. These cells were localised within and immediately adjacent to the cardinal vein (**Fig. 4l**), a major embryonic site of LEC emergence in mammals and zebrafish^108^.

Thus, by leveraging the HDCA, we have defined the timing and molecular heterogeneity underpinning human endothelial diversification during development. We reveal that organ-specific capillary EC transcriptional programs, particularly within the brain, heart and liver, are established as early as the first trimester of human development. Resolving lymphatic endothelial transcriptomes across time and space demonstrated the early establishment of molecular heterogeneity, with disease-associated lymphatic genes such as *PIEZO1* expressed at stages that precede the acquisition of fully differentiated lymphatic cell states.

### Temporal reconstruction of the developing human peripheral nervous system

The third and final distributed system we explored was the peripheral nervous system (PNS). The development of the human PNS, encompassing both sensory and autonomic domains, remains poorly characterised. This is largely due to its anatomical complexity, which spans the entire organism, and its dual developmental origin from two transient multipotent populations arising at the neural plate border: cranial placodes and neural crest cells (NCCs). Although somatic motor neurons contribute to peripheral nerve circuitry, they are neural tube–derived and therefore developmentally CNS in origin, and are not considered further here. Placode- and NCC-derived states have been characterised largely in model organisms^149,150^, while human studies have focused on isolated PNS compartments^151,152^, limiting our understanding of lineage contributions and tissue organisation.

Our single cell and spatial atlas identified the full repertoire of NCC-derivatives (*SOX10^+^* glial and *PRPH^+^* neuronal) across the embryo and placode-derived sensory neurons in cranial regions (*SIX1, EYA1, PRPH*) (**Fig. 5a; S23a-d**). NCCs are predicted to give rise to sensory, autonomic, and mesenchymal lineages, with a pseudotemporal analysis in mice suggesting that sensory specification precedes autonomic and mesenchymal differentiation^149^. In contrast, within the broad NCC *(SOX10, FOXD3)* compartment we observed sensory *(PRDM12)*, autonomic *(PHOX2B)*, and mesenchymal (*PDGFRA*) populations co-existing at 6PCW (**Fig. 5a-b**), precluding inference of a clear temporal ordering between sensory and autonomic specification. To identify the most undifferentiated NCC-derived cell states and characterise lineage differentiation, we applied Cell2fate^153^ as it outperforms existing RNA velocity models in reconstructing known cell fate trajectories and produces interpretable RNA velocity gene modules. This analysis confirmed that sensory and autonomic populations co-exist as transcriptionally active, early lineage-committed states, from which downstream sensory and autonomic trajectories are predicted to emerge (**Fig. S23e**). Multipotent sensory NCCs were predicted to progress through sensory neuronal progenitor states to form sensory neurons (*CNTN2*), while autonomic NCCs were predicted to develop into autonomic neuronal progenitor states (*PHOX2B, ASCL1)* and diverse autonomic neuronal derivatives (enteric (*RET*), sympathetic (*DBH, TH, MAOA*), parasympathetic (*SLC5A7*)), as well as sympathoblasts (*STMN2*) and chromaffin cells (*CHGA*) (**Fig. 5a; Fig. S23e**). These data provide the first comprehensive delineation of human PNS cell types and their lineage relationships, as summarised in **Fig. 5c**.

**Figure 5:**
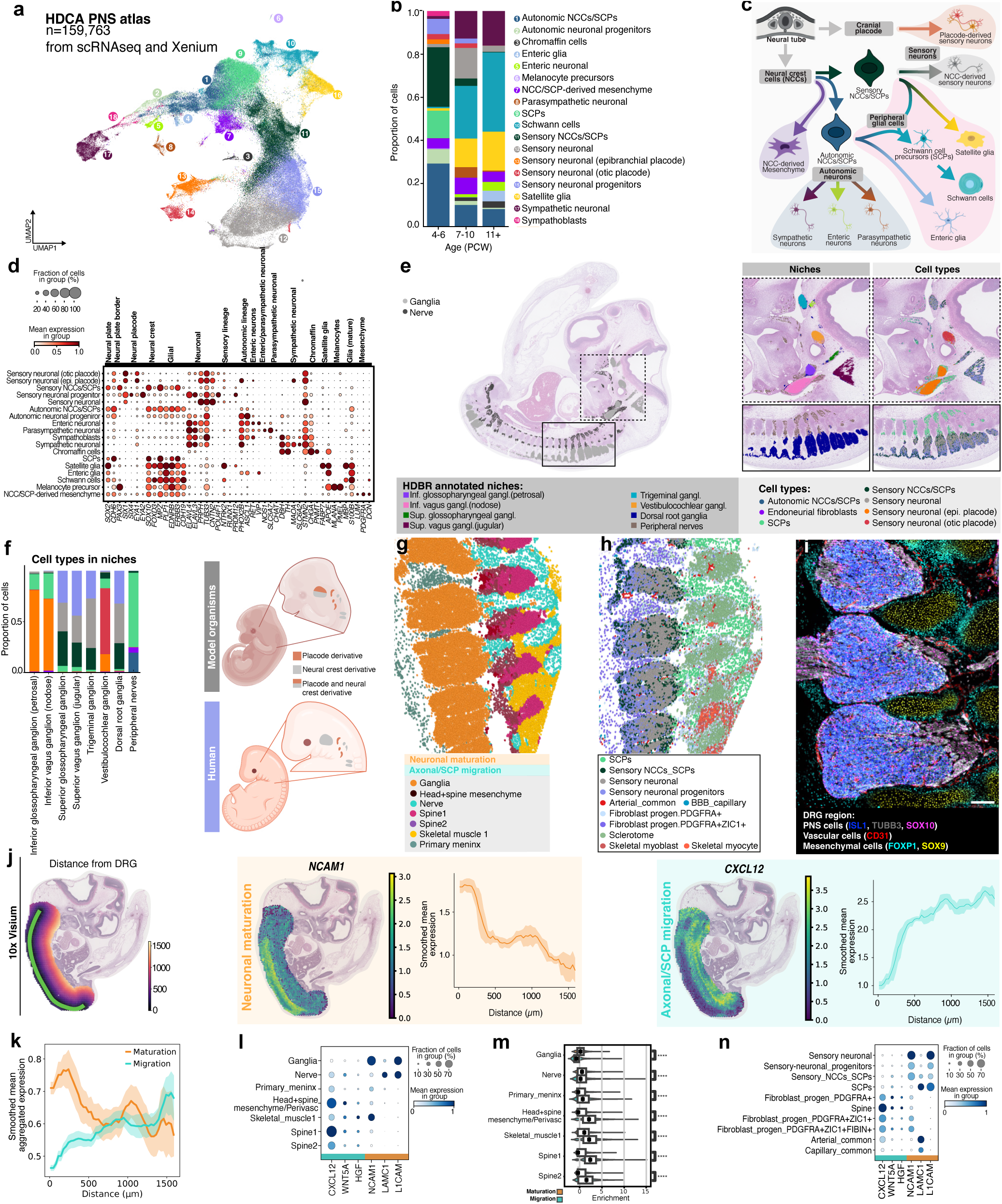
Temporal reconstruction of the developing human peripheral nervous system. **a)** UMAP visualisation of the PNS (n=159,763 cells) from both scRNA-seq and Xenium. Age, technologies, and lineage enrichment are shown in **Fig. S23a**. **b)** Barplot depicting cell type proportions per age. **c)** Schematic illustrating PNS trajectories. The diagram highlights the developmental and spatial progression of PNS lineages, and the emergence and differentiation of sensory and autonomic neuronal populations alongside associated progenitor and glial trajectories. Arrows indicate lineage relationships and transitional states, providing an overview of how distinct PNS cell types arise. **d)** Dotplot of marker gene expression for PNS populations. **e)** Left: Spatial location of the niches covering the PNS overlaid on H&E of a 6PCW embryo (section S133), and a UMAP coloured by cell type (3 slides). Right: Spatial location of the PNS populations of an embryo at 6PCW (left). Zoom into specific PNS areas (right) into head ganglia region (top) and DRG (bottom). H&E, HDBR annotated niches and cell types. **f)** Barplots depicting the proportion of PNS cell types found in each region, and schematic of cell origin per niche compared to model organisms. **g-h)** Xenium 10x niches (g) and cell types (h) in the DRG region of a 6PCW embryo (section S133) highlighting: neuronal maturation (left) and axonal/SCP migration (right). **i)** Section of human embryo (7PCW) showing DRG region immunostained for vascular (CD31), PNS (ISL1, TUBB3, SOX10) and mesenchymal cells (FOXP1, SOX9). Scale bar 200μm. **j)** Expression of *CXCL12* and *NCAM1* by distance from the DRG (proximal to distal planes) using Visium (section_121). Colour scale overlaid on H&E representative images indicate the normalised gene expression. Line plots depict the gene expression by distance from the DRG. **k)** Line plots of the mean gene expression by distance of genes associated with neuronal maturation and axonal/SCP migration (see **S21c** for individual ligands). **l)** Dotplot with the gene expression of ligands associated with axonal/SCP migration and neuronal maturation in niches in the DRG region of a 6PCW embryo (Xenium section S133). **m)** Violin plots depicting the enrichment of the gene signatures for neuronal maturation and axonal/SCP migration for the niches found in the DRG region of a 6PCW embryo (Xenium section S133). **n)** Dotplot of gene expression of ligands associated with axonal/SCP migration and neuronal maturation in the cell types within the DRG of 6PCW embryo (Xenium section S133). Dot size represents the proportion of cells within each group expressing a given gene. Color intensity indicates the scaled mean expression level. Gene expression values were standardised across genes to facilitate comparison. Statistical significance was determined by two-sided, Wilcoxon rank sum test with Bonferroni multiple testing correction. p < ****: p <= 1.00e-04.

NCCs are classically described as a progenitor population that is molecularly distinct from schwann cell precursors (SCPs), with each population defined by partially non-overlapping marker sets. However, recent studies in mice have proposed the existence of a multipotent NCC/SCP ‘hub’ state characterised by co-expression of NCC and SCP markers^154^. Consistent with this model, both sensory and autonomic NCC populations (*SOX10, FOXD3*) in the HDCA data universally co-expressed SCP markers (*MPZ*, *PLP1*) from the earliest stages examined (**Fig. 5d**). Despite the uniform co-expression of NCC and SCP markers, it was still possible that an NCC-only state exists transiently and is difficult to capture in human embryos. We thus leveraged the iPS-derived skin organoid system in which NCC-like populations are induced and can be interrogated from the earliest stages of differentiation^80,89^. The skin organoid dataset^89^ (**Fig. S23f-g**) similarly corroborated early co-expression of NCC and SCP markers, with no clearly separable NCC-only state detected, suggesting that such a state is either extremely transient or that prenatal human NCCs broadly adopt an SCP-like transcriptional identity.

The population of NCC-derived mesenchyme in our dataset co-expressed canonical NCC (S*OX10, FOXD3*) and mesenchymal markers (*PDGFRA, LUM, DCN*) (**Fig. 5a, c-d**). Notably, this population expressed *TWIST1*, a transcription factor shown to be critical for enabling trunk NCCs, which are primarily neuralgenic with limited mesenchymal potential compared to cranial NCCs^155^, to acquire ectomesenchymal potential in mice^149^. We also detected *FRZB* and *ZIC* gene expression (**Fig. S23h**), consistent with the above described *ZIC*^hi^ mesenchymal populations (**Fig. 3d, i**), which are also characterised by high *TWIST1* expression but no longer express NCC markers (**Fig. S23h**). The presence of these markers within the NCC-derived mesenchyme supports the above interpretation that *ZIC*^hi^ mesenchymal lineages are indeed NCC-derived and suggests that the NCC-derived mesenchyme population captured here represents a progenitor state giving rise to these cells. Consistent with this model, we detected an early mesenchymal population in the skin organoid NCC^89^ that co-expressed an NCC transcriptional signature alongside mesenchymal markers *FRZB*, *TWIST1*, and *ZIC*s at d6 (**Fig. S23g**) before the emergence of the 3 *ZIC*^hi^ populations at d26 (**Fig. 3l**). Notably, we found that the organoid NCCs, which possess a cranial identity^80,89^, gave rise to a greater abundance of mesenchymal cells relative to neural derivatives over time (**Fig. S23i**), consistent with the enhanced mesenchymal potential of cranial NCCs *in vivo*^155^. Together, these findings support a conserved NCC-to-ectomesenchyme transition across *in vivo* and organoid systems.

The NCC compartment in our dataset also encompassed a diverse range of peripheral glia (Schwann cells, satellite and enteric glia) and melanocytes (**Fig. 5a-d**). Temporal analysis revealed that neuronal specification occurs around 6PCW, while glial cells (**Fig. 5a-b, Fig. S23a**), as well as a glial maturation signature (**Fig. S23j**), emerged at later developmental stages. Thus consistent with model organisms^154,156^ and as previously shown for human DRG development^151^, overall, human peripheral nervous system development exhibits a temporal hierarchy, with neurogenesis preceding gliogenesis.

### Neural crest- and placode-derived sensory neurons

Sensory ganglia have been shown to arise from placodal and neural crest lineages in model organisms^149,150,157,158^; however, it is unknown whether the same developmental origins apply in humans. Transcription factors associated with placode- and neural crest–derived sensory lineages have been well characterised in model organisms and are frequently used as lineage markers; however, the extent to which these markers reliably distinguish placode-versus NCC-derived populations during human development remains unclear. We found that placode-associated transcription factors *EYA1/2* and *SIX1/4*^157,158^ were not restricted to placode-derived populations in human development, but were transiently expressed in NCC-derived sensory neuron progenitors, consistent with reports from human dorsal root ganglion (DRG) studies^151,159^ (**Fig. 5d**). In contrast, *SOX10* and *FOXD3* specifically marked NCC-derived populations, while *PRDM12* identified the sensory NCC-derived lineage, enabling clear discrimination between placode- and NCC-derived sensory neurons (**Fig. 5d**). These distinctions were further supported by 6PCW single nucleus multiome analysis showing that placode- and NCC-derived cells clustered separately (**Fig. S23k**), with chromatin accessibility at placode-associated genes broadly shared, whereas accessibility at NCC-associated genes, including *PRDM12*, was specific to NCC-derived sensory neurons (**Fig. S23l).**

We next mapped these populations spatially within PNS niches identified by NicheCompass (**Fig. 2**) across three Xenium tissue sections in which PNS niches were abundant (**Fig. 5e**). Ganglia niches corresponded to sensory ganglia at 6PCW and were enriched for neuronal transcripts (**Fig. 5e**). Of note, we only detected sensory ganglia, not parasympathetic, sympathetic or enteric ganglia, suggesting that although autonomic and sensory NCC/SCPs co-exist (**Fig. 5a, Fig. S23a**), sensory neuronal differentiation and ganglia maturation precedes autonomic. Leveraging these spatial and transcriptional data, we showed evidence for the developmental origins of human sensory ganglia for the first time (**Fig. 5f**). Consistent with animal studies^160,161^, we identified both placode-derived (vestibulocochlear, nodose and petrosal) and NCC-derived (dorsal root, jugular and superior glossopharyngeal) sensory ganglia in humans, each with distinct molecular signatures (**Fig. S23m**). Notably, we found that the human trigeminal ganglion consisted purely of NCC-derivatives, in contrast to the mixed origins reported in model organisms^162^. Together, these analyses provide the first human-specific map of sensory ganglia origins, defining placodal and neural crest derivation, revealing key differences from model organisms.

### The PNS development is guided by tissue niche interactions

Following ganglia formation, PNS development proceeds through axonal outgrowth toward target tissues, accompanied by the migration of SCPs along extending axons, where they support axon guidance and subsequent glial differentiation. To identify those axonal structures, we examined the second PNS cell–rich niche identified by NicheCompass across 6PCW Xenium embryos, which was distinct from ganglia. We termed this niche *Nerve* (**Fig. 2**), as it spatially mapped to peripheral nerve structures on H&E sections, both within the spinal region and along axonal extensions emerging from cranial sensory ganglia, and was enriched for NCC/SCP and SCP populations (**Fig. 5e**). In addition to axons, the *Nerve* niche also identified PNS compartments not discernible by H&E, such as the inferior hypogastric plexus (**Fig. S24a**), underscoring the power of our niche approach to resolve distinct gene expression profiles within histologically featureless tissues and across systems (beyond the pharyngeal endoderm shown in **Fig. 2**). NCC-derived mesenchymal populations also localised along axons, as well as around ganglia and in regions overlapping with *ZIC^+^* mesenchymal populations in the primary meninx, spine/DRG and scalp regions, again suggesting a direct lineage-relationship (**Fig. S24b**).

Using the NCC/SCP-rich *Nerve* niche as a proxy for tissue innervation, we found that, at 6PCW, PNS cells have not yet colonised organ parenchyma, in contrast to the mesenchymal and vascular compartments, which we have shown already display organ-specific identities at this stage (**Figs. 3-4**), indicating that mesenchymal and vascular niches are established prior to PNS innervation. The only exception was the gut, where autonomic NCC/SCP populations were detected that start forming the enteric nervous system (**Fig. S24a**), even earlier than previously reported^163^. In line with their later developmental emergence, mature enteric neurons^163^, enteric glia^163^, Schwann cells^151,159^, satellite glia^151,159^, and chromaffin cells^152^ were not observed in the 6PCW Xenium data (**Fig. 5b**).

To investigate how tissue niche interactions shape PNS development, we focused on the DRG and spinal nerve region, where sensory neurons undergo maturation and axonal outgrowth within a region composed of spatially adjacent PNS, mesenchymal and vascular niches and cells (**Fig. 5g-i**). We analysed spatially resolved ligand–receptor signalling across the previously identified niches within this region (**Fig. 5i-j**) using a literature- and Gene Ontology-curated set of signalling pathways implicated in neuronal maturation and axonal/SCP migration^164–166^. We first mapped ligand expression onto a tissue section using Visium to examine their spatial distribution (**Fig. 5j**). This analysis revealed opposing dorsoventral patterns: ligands associated with SCP migration and axon-associated guidance, such as CXCL12^167^, showed a ventrally enriched, gradual increase consistent with gradient-based, chemoattractive signalling, whereas neuronal maturation ligands, including *NCAM1*^168^, exhibited an inverse dorsally enriched distribution consistent with locally acting, short-range signalling (**Fig. 5j-k; Fig. S24c**). To identify the niche and cellular sources of these ligands, we analysed their expression across both niches and constituent cell types (**Fig. 5l-n**). Consistent with sensory neuron maturation occurring within ganglia, maturation-associated ligands (including *NCAM1*, *LAMC1* and *NTF3*; **Fig. 5l**) were enriched in the ganglia niche (**Fig. 5m**) and PNS-derived populations (**Fig. 5n**), although expression was not restricted to this compartment (**Fig. 5l)**. In contrast, cues associated with axonal outgrowth and SCP migration (e.g. *CXCL12*, *WNT5A* and *HGF*) showed higher relative expression within the Nerve niche and adjacent mesenchymal and vascular niches surrounding extending axons (**Fig. 5l-n**). Analysis of distance-aware cell–cell interaction and downstream signaling activity further confirmed that maturation-associated ligand–receptor pairs scored highest at short intercellular distances, supporting a local signalling mode, whereas migration-associated interactions were less distance constrained (**Fig. S24d-e**). These findings reveal spatially organised niche signalling programmes that coordinate PNS migration, differentiation and integration into embryonic tissues.

Notably, we detected expression of key regulators of sensory ganglia formation within the NGF signalling pathway, with *NGF* and *NTF3* expressed by mesenchymal cells and the receptor *NTRK1* expressed by sensory neurons (**Fig. S24f**). Given that genetic loss of these components in mice results in hypoplastic or absent DRG, extensive sensory neuron apoptosis, and impaired peripheral axon outgrowth^169–171^, these data suggest that NGF signalling pathways implicated in congenital peripheral neuropathies are active at this developmental stage. Together, our integrated atlas defines conserved and human-specific principles of PNS lineage specification, spatial organisation and intersystem communication during prenatal development.

## Discussion

This study presents the Human Developmental Cell Atlas (HDCA, n=186, k>4.6 million cells/nuclei, 4–22PCW), a unified resource integrating community-generated, organ-focused single cell/nucleus developmental datasets, alongside our complementary single cell and spatial transcriptomics (Xenium, Visium) data of 6PCW human embryos to fill a critical gap in late embryogenesis. Our HDCA data serves as a resource to contextualise human cells in development and disease, and those recapitulated within *in vitro* organoid systems. We’ve ensured the HDCA’s future reusability and continued biological insights by making data and models (scVI, CellTypist, scArches, NicheCompass) available, and creating a bespoke web portal for equitable access (https://cellatlas.io/studies/hdca/; private until publication).

The HDCA provides a temporally and spatially resolved view of how the human body is built. Using NicheCompass, we provide a comprehensive decomposition of the human embryo into 114 distinct tissue niches sampled across 11 sagittal whole embryo Xenium sections, representing a step-change in how cell types have been assessed and contextualised in human developmental single cell atlases to date. Niche decomposition allowed us to identify potential drivers of tissue patterning, and to define histologically featureless niches with distinct gene expression profiles, such as within the *Ant_head_mesenchyme* niche, which would not have been possible with cell types and spatial data alone. We make our NicheCompass models publicly available, enabling future community embryo niche identification both without expert histopathological annotation and even within a small number of sections.

We show how fibroblast progenitors diversify early to establish tissue niche architecture. Whether these tissue-specific embryonic fibroblast progenitors mature *in situ*, or migrate, remains an important open question. The HDCA also allowed us to resolve complexity within rare mesothelial populations (epithelial monolayers surrounding the outside of visceral organs) and the LPM populations from which they are derived. We identify shared and unique gene expression patterns between mesothelial cells across the whole embryo body, and detail, for example, LPM subtypes with distinct transcriptomic and spatial profiles. Detailing mesothelial ontogeny in development has implications for our understanding of aberrant programs in disease, such as mesotheliomas. Such detailed interrogation of mesothelium and fibroblasts have only been made possible through the pooling of these respectively scarce and abundant cells across datasets. Further, our integrated analysis across organs and systems enabled us to delineate NCC-derived mesenchymal populations, in human and across *in vivo* and organoid models.

Secondly, within the vascular system, we provide a comprehensive prenatal, cross-tissue atlas of human endothelial cells, encompassing arterial, venous, capillary and lymphatic compartments across development. Our analyses revealed that endothelial diversification occurs early, preceding the establishment of fully differentiated and functional organ systems^172^. We demonstrate early and rapid diversification of specialised vascular beds within the developing heart, central nervous system and liver, and that organ-specific capillary transcriptional programs^94,95^ are detectable as early as the mid-first trimester. This timing is in contrast to recent scRNA-seq studies of murine vascular development, in which organ-specific EC identity has been reported to emerge mid-gestation in mouse embryos^173^. The early emergence of these adult-like capillary programs in humans suggest that endothelial specialisation is coupled to early tissue patterning^98^, or intrinsic cues^174,175^, rather than arising as an adaptation to organ function in later development. Similarly, lymphatic endothelial heterogeneity was established early, including a distinct subset of early lymphatic cells in humans, enriched for genes with evolutionarily conserved roles in lymphatic development such as *FOXO1*, *CALCRL* and *PIEZO1*^117–121,145,147,148^. Notably, transcriptional signatures corresponding to adult lymphatic subtypes, including *PTX3-*expressing terminal capillary lymphatics^128,129^, were already detectable during human fetal development, indicating that lymphatic diversification precedes the acquisition of mature lymphatic patterning and organ-specific molecular phenotypes^111^. In contrast to mouse models, in which cranial lymphatics emerge postnatally^135–137^, we detect evidence for lymphatic cells in the human craniofacial region as early as 6PCW. Altogether, these findings support a model in which organ-specific capillaries and lymphatic endothelial heterogeneity are specified early in human gestation, providing a molecular substrate upon which later vascular remodelling, structural specialisation and functional maturation are built. These findings also have clinical implications, as we illustrate that the developmental timing of lymphatic gene expression has concordance with the timing of onset of primary lymphatic anomalies^143^.

Finally, within the PNS, we present a spatially resolved framework of human PNS ontogeny, capturing a broad repertoire of likely placode- and neural crest cell (NCC)-derived states, including sensory and autonomic neurons, peripheral glia, NCC-derived mesenchyme, sympathoblasts, chromaffin cells and melanocytes. We resolved lineage contributions across distinct ganglia niches and directly assessed placodal versus NCC origins of human sensory ganglia *in situ*. Overall, the inferred developmental origins of human sensory ganglia were largely consistent with those of model organisms^160,161^, with one notable exception: in contrast to the mixed placodal and NCC contributions reported in animal models^162^, the human trigeminal ganglion was composed exclusively of NCC-derived sensory neurons. Although we cannot formally exclude a transient placodal contribution at stages earlier than those sampled, the absence of a placodal signature in the human trigeminal ganglion (despite clear and distinguishable placode-derived populations in the petrosal, nodose and vestibulocochlear ganglia) argues against a substantial placodal input. This raises the possibility that, unlike in model organisms^162^, placodal input is functionally dispensable for trigeminal sensory roles in humans, or that NCC-derived sensory neurons can functionally substitute by adopting molecular and functional programs typically associated with placodal lineages, noting that lineage-specific roles within the trigeminal ganglion remain poorly defined even in model organisms^157,158,176^ and motivating future lineage-tracing and functional studies. Beyond lineage specification, our spatial analyses indicate that human PNS development involves temporally coordinated interactions between neighbouring tissue niches. Analysis of spatially resolved, known ligand-receptor interactions^164–166^ revealed a spatial division of labour in which mesenchymal and vascular compartments, established prior to PNS innervation, form an early scaffold that supports axonal outgrowth and Schwann cell precursor migration. Together, these findings underscore the ability of an holistic atlas such as the HDCA to reveal intersystem cross-talk events required for niche and tissue development.

The HDCA spans tissues and distributed tissue networks, and can therefore act as a springboard for researchers with diverse interests to explore prenatal cell types of interest. Decades of model organism developmental biology research provided crucial literature to contextualise many of the cell types and lineages in this study, however, these were never detailed in humans before, or at such scale. We report both human-specific aspects of development and those which are conserved with model organisms, with the hope that this study can more widely act as a human developmental cell reference for understanding developmental disorders in model systems. Organ biologists can use the HCDA to understand how cellular composition and tissue architecture forms across time and space, and researchers interested in inter-system cross-talk can leverage this unique resource to comprehensively evaluate such interactions at a niche-level, as here in the context of the PNS-mesenchyme. Our data and models provide a foundation for future research integrating new developmental datasets, we hope even bridging the gap to the earliest implantation stages.

Looking toward the utility of a future HDCAv2^5^, important challenges to tackle include data generation from undersampled tissues, such as from the endocrine system and the time points between implantation and 4PCW. Spatial profiling at sequential timepoints, e.g., 8PCW and 10PCW (but especially those from less represented embryonic and late fetal timepoints) combined with niche inference, would be a powerful approach to capture transient tissue microenvironments, especially given that our fixed sampling at only 6PCW was able to resolve embryonic tissue drivers, granular niche capture and cellular composition of pharyngeal niches and hitherto undescribed cranial niches. While we provide the most comprehensive assessment of niche capture to date from 11 sagittal sections and >4 million cells, more comprehensive profiling is required to fully capture niche composition across the whole embryo. More widely, inferring molecular information from H&E is an open problem in computational and clinical research^177^, and niche decomposition provides one method for tackling this open problem at organism-scale by discretisation into functionally relevant histological groupings. The translational applications of the HDCA are wide-ranging, particularly with our Cherita webportal integration with the DDG2P dataset (https://www.ebi.ac.uk/gene2phenotype)^43^ allowing for both gene-cell and gene-niche investigation into developmental disorders for which genetic variants have been identified.

In summary, this study represents a major milestone toward a unified and complete HDCA that will be a transformative resource for research and clinical communities. Together, by enabling development to be analysed through spatially and temporally defined tissue niches, our HDCA shows that, similar to architecture, form follows function as tissue structure emerges from coordinated interactions between cells, tissues and systems.

## Methods

The Human Developmental Cell Atlas (HDCA) was constructed by integrating 2 complementary sources of single cell RNA-sequencing data: (i) a whole-embryo (WE) scRNA-seq dataset generated de novo in this study, and (ii) a large collection of publicly available community datasets (Community data). The annotations for the community datasets were informed by the original author-provided labels and subsequently harmonised across studies. In contrast, the WE dataset lacked pre-existing cell type annotations and extensively manually curated through literature informed gene markers and expert input, augmented by machine learning models generated using the community data. The final HDCA was generated by jointly integrating and reconciling information from both datasets, leveraging the strengths of de novo annotation in the WE data and the diversity of community-derived annotations. In this study we specifically focussed on: mesenchyme, endothelial cells and the peripheral nervous system. Below, we describe the strategies for separate sample preprocessing, quality control, integration, clustering and cell type annotation of WE and Community data and how they informed annotation of HDCA. All code supporting the methods and analyses presented in this study will be made available at https://github.com/haniffalab/hdca-paper upon publication.

### Community data: Public human developmental single cell data acquisition and preprocessing

We collected 21 community scRNAseq datasets, comprising 18 publicly available ones and 3 obtained through internal communications, representing diverse tissues across the entire body of the human embryo and/or fetus (**Supplementary Table 1, 5, 7**). The selected datasets were accessed in 2022, and fulfilled the following 3 criteria: 1) consistent technology (10x Genomics Chromium), 2) minimum 750 cells post-QC per organ (see below for QC thresholds), 3) the majority of annotation labels provided were granular, i.e., to the cell type level ‘monocyte’ rather than lineage ‘myeloid’. To render comparability between pre-existing, pre-processed single-cell RNA-sequencing (scRNA-seq) studies, together comprising over 3.6 billion cells, we implemented a scalable unification pipeline that systematically harmonised the feature space, standardised metadata, and concatenated the expression matrices into a single analytic object. For each dataset, the raw count matrix was downloaded along with the accompanying cell meta information. After removing trivial cell types annotated by the original studies (for example, “Doublets”), we used a loose quality control criterion by requiring a minimum number of expressed genes per cell > 200, and obtained a total of 3,652,263 cells representing all major cell populations in the human embryo (**Supplementary Table 1**). Thirty-one curated metadata fields, including biological attributes (organ, age), donor-specific identifiers (integration_donor), and technical descriptors such as sequencing_platform, were harmonised and appended to the cell-level annotation table, thereby establishing a consistent observational schema. Comprehensive provenance information, including dataset identifiers, original publication details, accession codes, and acquisition metadata, is catalogued in **Supplementary Table 5**. For datasets obtained through restricted-access agreements or pre-publication data-sharing arrangements, relevant ethical approvals and consent frameworks are documented in the original source publications.

### Community data: Realignment and gene space harmonisation

We used the preexisting data from the original publications. To ensure all datasets were aligned to GRCh38, Miller et al 2020^178^, Sridhar et al 2020^179^, and Yu et al 2021^180^ were remapped to GRCh38 2020-A as outlined in **Supplementary Table 1**. To resolve assay- and release-specific disparities in gene nomenclature for external datasets, we employed the pre-release IDTrack algorithm^181^ (see https://github.com/theislab/idtrack), a graph-based identifier harmonization framework designed to reconcile discrepancies in gene nomenclature across large-scale transcriptomic datasets. Variability in gene identifiers arises not only from the use of distinct annotation systems such as Ensembl and RefSeq, but also from differences between annotation releases, which evolve over time and contribute to inconsistencies in gene symbol conventions. In our integration effort, datasets derived from various genome builds and annotation versions were standardized by mapping all gene identifiers to a unified target space defined by HUGO Gene Nomenclature Committee (HGNC) gene symbols corresponding to Ensembl release 107. Gene symbols that mapped unambiguously to a single Ensembl identifier were retained without modification, whereas one-to-many relations were adjudicated by prioritising biologically verified target identifiers, and genes exhibiting irreconcilable or failed conversions were removed. Dataset-level statistics, such as the numbers of genes whose mapping changed only because of one-to-many conflicts or because of alternative, equally valid one-to-one targets, were logged to enable traceability, and the full conversion table was stored within the unified object for subsequent audit. When combined with WE scRNA-seq data to form the complete HDCA data, only genes which intersected with the genes post IDTrack were retained.

### Community data: Highly variable genes selection pipeline

To enable robust, scalable, and biologically meaningful downstream analyses across millions of single cells, we developed a comprehensive feature selection pipeline designed to eliminate confounding technical signals, enforce rigorous quality control, and identify highly variable genes (HVGs) in a batch-aware manner. Beginning with a consolidated expression matrix comprising 3,611,521 single-cell transcriptomes profiled across 57,009 genes, we removed 516 genes annotated to the cell-cycle process, 400 immunoglobulin heavy- and light-chain genes encoding B-cell receptors, and 227 αβ or γδ T-cell receptor genes, whereas mitochondrial and cytoplasmic ribosomal transcripts were retained to preserve markers of cellular fitness and translational capacity; this operation reduced the data matrix to 3,609,265 cells by 55,866 genes. Subsequent quality control removed 2,256 cells (0.06 % of the total) with fewer than 200 detected genes, and filtered 11,490 genes supported by fewer than 5 UMI counts across the entire cohort, yielding a high-fidelity dataset of 3,609,265 cells and 44,376 genes that served as the input for variable-gene selection step.

Raw gene counts were normalised using *sc.pp.normalise_total* function with target sum as 10^4^ and then a shifted log transformation was applied using the *sc.pp.log1p* function. The batch_key was defined by the Cartesian product of 4 covariates —donor identity, biological unit, sample status, and sequencing platform, resulting in 192 batches. For each gene, mean expression was constrained to the interval 0.05 ≤ μ ≤ 6 to exclude extremely rare or constitutively expressed features, after which 219 genes falling outside the specified expression bounds were discarded. Normalized dispersion values were computed independently for each batch and also aggregated globally. From the global rankings, the top 12,288 genes (2¹⁴ – 2¹²) exhibiting the highest dispersion were retained as candidate HVGs for downstream analysis. Concordance analysis revealed a median Spearman correlation between global and batch-specific dispersion rankings that exceeded that expected by chance, and the intersection between the top 10,000 HVGs derived from the global model and the per-batch selections reached 75.4 % across batches, attesting to the stability of the procedure. Moreover, a Wilcoxon rank-sum test comparing dispersion profiles obtained with and without batch correction yielded P < 10⁻¹⁵, underscoring the statistical benefit of the batch-aware strategy.

The final unified dataset, comprising 57,009 harmonised genes and 3,611,521 quality-controlled cells contains (i) gene-level annotations that include harmonised ENSEMBL names and intersection flags, (ii) complete provenance records listing the unification_id, preprocessing_id, IDTrack parameters, and per-study conversion logs in adata.uns, and (iii) the full set of 31 standardised metadata columns. By enforcing stringent quality and formatting standards, this uniformization strategy ensures that all downstream analyses operate on a reproducible and harmonized data foundation, thereby enabling rigorous and unbiased inference in single cell transcriptomic studies.

### Community data: Integration with scVI

To address the multifaceted challenge of integrating single cell RNA sequencing data spanning multiple organs, developmental time points, and assays, we implemented a cell-type-agnostic training and validation strategy based on the single cell Variational Inference (scVI)^182^ framework. An extensive hyperparameter search was conducted in 2 phases to identify a suitable network architecture and training scheme for large-scale data integration as described below. This approach was designed to accommodate the inherently high-dimensional and sparse nature of early developmental atlases (i.e., genes × cells × time × organ) and to mitigate the numerous technical confounders frequently encountered in large-scale integration, including unbalanced sampling across organs and time points, protocol-specific batch effects (e.g., fresh versus frozen, nuclei versus whole cell, 5′ versus 3′ sequencing). In addition to scVI, we evaluated other cell-type-agnostic methods, including scPoli (deployed without prototype initialization), TarDis (using batch labels only as the disentanglement target and did not incorporate any auxiliary covariates like cell-type annotations), Scanorama, Harmony, and principal components analysis (PCA), each subjected to a 40-run hyperparameter tuning process.

In the first phase of model training, an extensive hyperparameter search spanning more than 4 thousand configurations of scVI was conducted to identify a suitable network architecture and training scheme for large-scale data integration. Model hyperparameters, encompassing the dimensionality of hidden layers, the timing and magnitude of Kullback–Leibler regularization warm-up, dropout rates, and learning rates, as well as dataset-level considerations such as the strategy for selecting highly variable genes (union versus intersection across datasets) and the number of input features were systematically varied. For each configuration, multiple batch definitions were tested to ascertain whether biological attributes or technical factors (e.g., donor identity, sequencing platform) should be explicitly encoded as batch variables, reflecting the heterogeneity intrinsic to multi-organ, multi-time-point studies. A preliminary benchmarking pipeline computed R² scores on differentially expressed genes, along with a rapid subset of integration quality metrics from the scIB suite, thereby enabling high-throughput evaluation across many candidate models. Early convergence patterns revealed pronounced sensitivity to parameters controlling regularisation strength—particularly weight decay and the duration of KL warm-up, and to architectural depth, whereas broader sampling of activation functions and optimiser variants yielded diminishing returns. Accordingly, search density was adaptively shifted toward the high-impact subspace, with occasional manual refinements applied when intermediate checkpoints indicated latent spaces with superior topology preservation but suboptimal reconstruction. Empirical validation of these early trials led to a narrowed set of hyperparameters that balanced model expressiveness and training stability while mitigating overfitting.

To further refine integration performance, an additional 40 training configurations of scVI were also explored to refine the integration pipeline and corroborate the hyperparameter settings derived from the initial large-scale optimization. We also evaluated other cell-type-agnostic methods, scPoli, TarDis, Scanorama, Harmony, and principal components analysis (PCA), following the 40-run hyperparameter tuning process. Importantly, the knowledge gained during the preliminary 4,000-configuration scVI sweep guided parameter selection for subsequent runs of autoencoder based models (scVI, scPoli^183^ and TarDis^184^), thereby aligning each model’s architecture and training strategy with empirically validated best practices while retaining sufficient variation to identify the most robust approach. Scanorama and Harmony were executed with their default parameters. Benchmarking of the models were performed using scIB and scTRAM^45^ metrics to quantify the balance between batch removal, biological conservation, and trajectory representation fidelity.

To establish a baseline for evaluating integration performance, we computed differentially expressed genes (DEGs) using the Wilcoxon rank-sum test implemented in scanpy‘s method, ‘tl.rank_genes_groups’. Genes were ranked by log-fold changes, adjusted p-values (padj<0.05), and significance thresholds, with results stored in the AnnData object’s unstructured (uns) metadata. This preliminary analysis enabled post hoc validation of cell-type-specific markers preserved during integration.

A cell-type-agnostic modelling paradigm was adopted for several reasons rooted in the intrinsic complexity and partial annotation of large-scale developmental atlases. Because emerging cell populations in early embryonic or fetal stages may lack definitive markers, strictly label-based strategies risk propagating bias from uncertain or incomplete annotations. Furthermore, dynamic cell definitions can shift abruptly across time points or organs, rendering static category assignments unreliable. By avoiding explicit cell-type priors, we aimed to preserve flexibility in uncovering rare or transitional states, as well as accommodate divergent nomenclature across datasets without prematurely imposing a fixed annotation framework. This approach thereby emphasized the inherent structure of the data over potentially erroneous or heterogeneous cell-type labels.

### Community data: Benchmarking integration and trajectories with scIB and scTRAM

To systematically evaluate how effectively different integration methods and model configurations preserve both batch correction and developmental biology in the multi-organ human developmental cell atlas, we implemented a dual-framework benchmarking strategy combining the established scIB^44^ protocol with scTRAM^45^, a trajectory-fidelity metric suite specifically designed to assess preservation of developmental progressions in integrated representations. Cell embeddings with different hyperparameters were obtained using scVI, scPoli, TarDis, Harmony, Scanorama, and principal components analysis (PCA). The scVI, scPoli, and TarDis models were trained with architectures and hyperparameters derived from the extensive 4,000-configuration optimization described in the *Data Integration* section, ensuring each autoencoder-based method operated under empirically validated best practices. Harmony and Scanorama were executed with their default parameters as specified in their respective implementations. For all methods, the batch_key was defined as mentioned above in the selection of highly variable genes section.

### scIB: Conventional integration quality assessment

The scIB benchmarking framework quantifies integration performance through a comprehensive suite of complementary metrics conceptually partitioned into 2 categories: biological conservation and batch removal. Biological conservation metrics assess the extent to which intrinsic cellular structure and lineage relationships are maintained following integration, encompassing measures such as cell-type average silhouette width, normalized mutual information, adjusted rand index, isolated label F1 score, and local inverse Simpson’s index computed with respect to cell-type labels (cLISI). These metrics collectively evaluate whether the integrated representation preserves discrete cell-type identities, maintains neighbourhood composition, and prevents unwarranted amalgamation of biologically distinct populations. Batch removal metrics quantify the effectiveness with which technical covariates are suppressed, including graph connectivity, batch average silhouette width, k-nearest-neighbour batch effect test, principal component regression, and local inverse Simpson’s index computed with respect to batch labels (iLISI). These measures assess whether batch-associated structure has been adequately mitigated without introducing artificial discontinuities or over-correction artefacts.

For each integration method, individual metrics were computed on latent representations using hierarchical cell-type annotations at both broad cell level and fine-grained subtypes as ground-truth references. Metric values were min–max scaled within their respective categories, directionalities harmonized such that higher values uniformly indicate superior performance, and subsequently averaged within biological conservation and batch removal groups before computing a composite aggregate score. This normalization procedure enabled direct ranking of integration strategies according to the joint criterion of biological fidelity and technical robustness, thereby facilitating systematic comparison across methods with heterogeneous performance profiles.

### scTRAM: Trajectory-specific fidelity evaluation

While scIB effectively quantifies batch correction and cluster alignment, it does not capture whether embeddings preserve the differential-geometric properties of continuous cellular transitions that underpin lineage commitment and differentiation dynamics. To address this limitation, we applied scTRAM, single cell trajectory representation metrics, a principled benchmarking framework that systematically evaluates how well integrated representations retain ground-truth developmental trajectories across complementary failure modes.

Developmental trajectories were formalized as directed acyclic graphs G_ref_ = (V, E), where vertices V correspond to discrete cell types and directed edges E represent validated state transitions between cell types. We curated 3 exemplar reference systems with distinct developmental architectures: hematopoiesis (57 cell types, 58 directed connections), germline development (8 cell types, 7 connections), and the peripheral nervous system (12 cell types, 11 connections). These reference graphs captured diverse trajectory topologies ranging from hierarchical branching structures in hematopoiesis to linear progressions in germline development, ensuring comprehensive evaluation across multiple scales of developmental complexity. Ground-truth trajectories and corresponding cell-type annotations for each system are presented in **Supplementary Figure 7**.

scTRAM evaluates trajectory fidelity by comparing inferred developmental structures in the latent representation against the curated reference lineage graph G_ref_ through 3 complementary metric groups that decompose trajectory preservation into distinct performance axes. Topology conservation metrics (adjacency-based) assess whether the branching architecture and connectivity patterns of the reference graph are maintained in the integrated representation, quantifying structural similarity through measures including graph edit distance, spectral distance, and topological data analysis approaches, among others. Cell-order continuity metrics (pseudotime-based) evaluate whether cells are arranged in the correct developmental progression along directed edges, testing temporal monotonicity through rank correlations such as Spearman’s ρ, dynamic time warping distance, and concordance indices, alongside additional ordering-based measures. Trajectory dominance metrics (embedding-based) determine whether the developmental manifold remains the primary source of variation in the latent space rather than being obscured by batch effects or other nuisance structure, employing manifold preservation measures such as neighbourhood conservation, trustworthiness, and geodesic stress, supplemented by directionality and continuity assessments. For each integration method, diffusion-based pseudotime and cell-type connectivity graphs were inferred from the latent representation, then compared against reference pseudotimes and adjacencies derived from G_ref_ using the full battery of metrics within each group. Individual metric scores were subsequently aggregated within their respective categories to produce topology conservation, cell-order continuity, and trajectory dominance summary scores for systematic comparison across integration methods.

### Edge-resolved and lineage-stratified analysis

To enable fine-grained performance assessment across developmental transitions, we partitioned each reference lineage graph into constituent trajectory segments using an automated decomposition algorithm. This algorithm identified hierarchical functional modules by extracting induced subgraphs over all branch points and their descendants, enumerated branch-to-leaf paths from the trajectory root to all terminal cell types, traced linear chains from each bifurcation point’s immediate successors until subsequent branch points, and computed the longest path through the developmental hierarchy via dynamic programming. Candidate subgraphs were retained only if they contained at least 3 nodes, exhibited weak connectivity, and possessed exactly one root node. This multi-resolution decomposition yielded hundreds of trajectory segments spanning diverse developmental transitions from stem cell activation to terminal differentiation, ensuring balanced coverage of distinct developmental pathways and enabling localized performance analysis that aggregate metrics would obscure.

To localize performance differences along developmental paths, we computed per-edge scTRAM scores across the 3 curated lineage graphs. For each directed edge in G_ref_, we identified all trajectory segments containing that edge and calculated the mean performance rank across all applicable metrics within those segments, thereby deriving edge-specific fidelity scores for each integration method. Differences between methods were assessed using two-sided paired comparisons after multiple-testing correction via the Benjamini–Hochberg procedure at a 5% false discovery rate threshold. This edge-resolved analysis revealed non-uniform, stage-specific behaviour wherein fidelity varied continuously along developmental progressions and models selectively preserved or distorted particular developmental segments.

In parallel, trajectory-specific analysis was performed by grouping the decomposed segments according to major lineage classes (myeloid, lymphoid, megakaryocyte-erythroid-mast, and stem/progenitor), computing mean ranks within each group, and evaluating whether methods exhibited systematic biases in handling different branching complexities or developmental contexts. Performance was additionally stratified by scTRAM metric group (pseudotime-based, adjacency-based, embedding-trajectory) to assess whether models optimized for preserving topological connectivity sacrificed fidelity in pseudotemporal ordering or vice versa.

### Integrated benchmarking framework

For each latent representation, scIB and scTRAM metrics were computed independently and subsequently combined to provide a comprehensive evaluation of integration quality. Individual scTRAM metrics were min–max scaled and averaged within their respective groups (topology conservation, cell-order continuity, trajectory dominance) before computing a trajectory representation aggregate score. Integration methods were ranked according to both conventional scIB criteria (biological conservation and batch removal) and trajectory-specific scTRAM criteria, with particular attention directed toward identifying trade-offs between batch correction and lineage preservation. Qualitative visual inspections of lineage-restricted subsets complemented quantitative assessments, ascertaining preservation of rare transitional states and detecting unwarranted amalgamation of biologically distinct populations. Particular scrutiny was directed toward the retention of developmental trajectories across organs and time points and toward the attenuation of artefacts attributable to protocol heterogeneity, most notably differences between fresh and frozen preparations, nuclei-versus whole cell captures, and 5′-versus 3′-biased library chemistries.

This dual-framework approach enabled rigorous model selection that balanced conventional integration objectives with trajectory-specific requirements, ensuring that the chosen representation not only achieved effective batch correction and cluster purity but also faithfully preserved the continuous manifold structures essential for downstream analyses of developmental progression, lineage commitment, and transcriptional dynamics along differentiation paths.

### Community data: Cell type annotation

In the first iteration, we adopted an automated method for annotating the community data based on the author provided annotations for each constituent dataset, followed by manual curation for standardising cell type names. We trained a logistic regression-based model for each previously published dataset using CellTypist^30^ (version 1.4.0), publicly available at https://www.celltypist.org/models, with the function *celltypist.train* (tag-value pairs: max_iter=1000, feature_selection=True, top_genes=300). We cross-predicted cell identities for each pair of datasets using these models with the function *celltypist.annotate*, and based on these predictions we standardised cell type names across datasets to achieve a common naming scheme. Specifically, we collapsed cell types from different datasets only if they could be reciprocally predicted and thus corresponded to a single cell type (e.g., alignment between “*Multiciliated precursor*” from Miller et al. 2020^178^ and “*Deuterosomal*” from He et al. 2022^20^). Moreover, we located and subsequently refined a coarse cell type only if it was divisible into subtypes from another dataset and the latter could be uniquely mapped back (e.g., subdivision of “*pro_B*” from Garcia-Alonso et al. 2022^185^ into “*PRO_B*” and “*LARGE_PRE_B*” from Suo et al. 2022^24^). The resulting cell types underwent another round of manual curation, including inspection of marker gene expression and renaming of cell types, which resulted in a final list of 536 cell types curated from 766 author-defined original cell annotations (https://www.celltypist.org/encyclopedia/Human_Whole_Embryo/v1).

To test the performance of this model, we split the entire data set into a set of training (90%) and the remaining set of test dataset (10%). A CellTypist model was then trained from the training set and used to predict cell types for the test dataset. Specifically, we defined one iteration as one mini-batch training, with each batch containing 1,000 cells randomly selected from the training dataset. A CellTypist model was derived after each iteration and was then used to predict cell types for the test dataset. This process was repeated over 30 epochs, with each epoch consisting of 100 mini-batches; in other words, a total of 3,000 iterations were performed (**Fig. S1e**). The model performance was assessed for each cell type separately using 3 metrics: precision (“*sklearn.metrics.precision_score*”), recall (“*sklearn.metrics.recall_score*”), and F1 score (“*sklearn.metrics.f1_score*”). Lastly, we averaged these scores across cell types to obtain the performance curves, showing the model performance as a function of training iterations (**Fig. S1e**). Second, we tested the performance of the final model (i.e., the model trained from 100% of the data) on an independent dataset profiling the human embryo using sciRNA-seq3 (Cao et al. 2020)^47^, and visualised the cell type correspondence between the model and the tested dataset using *celltypist.dotplot* (**Fig. S1f**). In the second iteration the resultant cell types were manually crosschecked with original author annotations per leiden cluster (resolution=3), considering organ and lineage diversity to annotate the cells for the community data. This model is used to increase the power of manual annotations of the de novo Whole Embryo dataset. In the second iteration the resultant cell types were manually crosschecked with original author annotations per leiden cluster (resolution=3), considering organ and lineage diversity to annotate the cells for the community data.

### Whole embryo: sample acquisition and processing

Human embryos for scRNAseq/multiome (samples F137, F147, F158) and for spatial Xenium, Visium, RareCyte (samples SOB25, SOB26) were collected by the MRC– Wellcome Trust-funded Human Developmental Biology Resource (HDBR; http://www.hdbr.org; MR/R006237/1, MR/X008304/1 and 226202/Z/22/Z) with written consent and approval from the Newcastle and North Tyneside NHS Health Authority Joint Ethics Committee (23/NE/0135) (**Supplementary Table 1**). One human embryo for spatial MACSIMA (sample SM74P7) was provided by the HuDeCA Biobank (Inserm) following French ethical and legal guidelines. Donors provided written informed consent prior to sample procurement. Ethical approval for the use of these materials was granted by the Agence de la Biomédecine (Saint-Denis La Plaine, France; Authorization N° PFS19-012) and the INSERM Institutional Review Board (IRB00003888).

### Whole embryo: scRNAseq sample processing, library preparation and sequencing

Each of the three whole human embryo samples for scRNAseq/multiome were dissected into twelve anatomical regions using a scalpel (**Supplementary Table 2; Fig. S1a-e**). Unfortunately, Embryo 1 (Haniffa Lab sample ID F137) was received with one lower limb missing. Each tissue region was stored in HypoThermosol® Preservation Media (STEMCELL Technologies) for a maximum of sixteen hours refrigerated. Each tissue was then washed with phosphate buffered saline (PBS, Sigma-Aldrich) before mincing with a scalpel. The minced tissue was digested with 1.6 mg/ml type IV collagenase (Worthington) in RPMI (Sigma-Aldrich) at 37°C for 30 min with intermittent agitation. Digested tissue was then passed through a 100μm cell strainer and 0.25% (w/v) trypsin (Sigma-Aldrich) was added to any undigested on the strainer mesh for 5 min before adding RPMI supplemented with 10% (v/v) heat inactivated fetal bovine serum (FBS), 1% (v/v) penicillin streptomycin and 1% (v/v) L-glutamine. Cells were then collected by centrifugation (500 g for 5 min at 4°C) and manually counted using an Improved Neubauer hemocytometer. Cells from each tissue region were stained with DAPI (Sigma-Aldrich) at a final concentration of 3μM and passed through a 35µm filter (Falcon) before FAC-sorting to select for only live, single cells (**Fig. S1c**).

Immediately after sorting, cells were then manually counted using an Improved Neubauer hemocytometer. Where possible, 4 channels of cells from each tissue region were loaded onto the 10x Genomics Chromium Controller to target 10,000 cells per channel. Embryo 1 (Haniffa Lab sample ID F137) was processed with the 10x Genomics Chromium Single Cell 5’ Reagent Kit v1 and embryos 2 and 3 (Haniffa Lab sample ID F147 and F158) were processed with the 10x Genomics Chromium Next-GEM Single Cell 5’ Reagent Kit v2, as per the manufacturer’s instructions. Sequencing libraries were also processed with the 10x Genomics proprietary reagents. Libraries were then sequenced using Illumina NovaSeq S4 Flow Cells to target a minimum sequencing depth of 50,000 UMIs per cell.

### Whole embryo: scRNAseq alignment and quality control

Whole embryo (WE) scRNAseq data was mapped to the human reference genome (GRCh38-2020-A) using CellRanger (v6.0.1). Empty drops were excluded from analysis through use of filtered CellRanger matrices. 2 low quality lanes were excluded: WS_wEMB10202360 had low fraction of valid barcodes; WS_wEMB11031943 had low number of cells detected. This produced an initial count matrix of size 1,183,081 cells x 36,601 genes).

Filtering was performed to select for cells expressing >1,000 UMIs and >=500 genes, and genes expressed in >=5 cells (QC step 1/2), leaving a count matrix of size 1,138,964 cells x 34,584 genes. Low quality cells as determined by high mitochondrial content were calculated and flagged for later evaluation alongside annotation information. We aimed to use the strictest threshold available while ensuring the majority of any annotated cell type category was not excluded from downstream analysis. Cells with mitochondrial reads <5%, 10%, 15% and 20% were evaluated for selection, and cells with <10% mitochondrial reads fulfilled criteria above and were selected for downstream analysis. To assess doublet contribution to the scRNAseq data, Scrublet (v0.2.3) was applied to each lane independently, obtaining per cell doublet scores. An exclusion threshold of >(median + (3 * median absolute deviation)) Scrublet score was applied on a per cell basis, as previously described^16,17^.

To investigate the potential of maternal contamination we followed a similar protocol outlined in Dann and Suo et al^24^. Every 10x lane from each sample was put through Souporcell (v2.4) with genotype clusters k=1/2/3 models to represent the likelihood of single or multiple genotypes (2nd genotype represents maternal, 3rd represents control and artefacts). Optimal model selection was determined using BIC (bayesian information criterion). Most 10x lanes were determined to be a singular genotype. The optimal model for lane WS_wEMB10202336 was k=3, but given no technical replicates were available for this sample:section (F137 torso section 1) these data were retained. Of note, 2 genotypes were detected for lanes WS_wEMB10202351, WS_wEMB10202353 and WS_wEMB10202356 (from sample F137 torso sections 5 and 6). The cells contributing to the smaller genotype cluster were flagged as potential maternal cells and recorded in metadata prior to annotation, then removed after annotation was complete.

Following removal of cells based on high mitochondrial content, doublet status, and maternal contaminant status (QC step 2/2), the count matrix was of size 1,029,326 cells x 34,584 genes.

To investigate the level of ambient RNA within the scRNAseq data, Cellbender (v0.2.0) was run for each high quality lane. The resultant soup-corrected CellBender filtered matrix summed to 1,363,423 cells x 36,601 genes. A ‘soup’ matrix was then generated by taking the difference between CellBender-soup-corrected counts and the non-corrected CellRanger counts. Given the more permissive cell calling employed by CellBender vs CellRanger, we had to access the equivalent counts for the non-corrected 1,363,423 cells through use of unfiltered CellRanger matrices. CellBender was run on a per sample basis with the following parameters: expected yield of 10,000 cells per lane, total-droplets-included=20,000, fpr=0.01, epochs=150. Parameters were adapted by lane based on CellRanger barcode rank plot: WS_wEMB10202336 expected cells parameter was lowered to 5000; WS_wEMB11031950 total-droplets-included was set to 60,000; WS_wEMB12142148 total-droplets-included was set to 35,000; WS_wEMB12142131 failed due to insufficient background, therefore no corrected raw counts were obtained. These soup matrices then facilitated investigations into ambient RNA contribution across each sample. The top 200 gene counts from each sample’s soup matrix were recorded (**Supplementary Table 4**) and the top 50 were recorded in the data for reference in downstream annotation and analysis. Due to overall low contribution of ambient RNA across all samples, CellRanger filtered counts were used throughout preprocessing and analysis.

Preprocessing for the scRNAseq data followed similar logic outlined in the Scanpy (v1.7.2) workflow tutorial^186^. Raw gene counts were normalised using *sc.pp.normalise_per_cell* function with counts_per_cell_after=1e4 and then a ln(x+1) transformation was applied using the *sc.pp.log1p* function. For any independent analysis shown, expression values reported are normalised then log-transformed as above.

### Whole embryo: scRNAseq integration, batch correction and visualisation

For WE scRNA-seq integration, cell cycling genes were obtained from an public integrative fetal scRNAseq atlas^24^, and the 542 intersecting cell cycle genes present in our dataset were excluded. Data were then log-normalised as described in ‘*scRNAseq gene expression visualisation using dotplots and heatmaps*’. Highly variable gene (HVG) selection was performed using the *sc.pp.highly_variable_genes* function from Scanpy (*min_mean=0.0125, max_mean=3, batch_key=’haniffa_ID’*), and the top 10,000 highly variable genes HVGs ranked by *‘dispersions_norm’* were flagged. For dimensionality reduction and integration, we used the scVI module from scvi-tools (v0.19.0). The top 10,000 HVGs identified above were input into scVI as raw expression counts. The scVI workflow^182^ was then followed, with biological replicate and 10x chemistry taken as categorical covariates, and 256 nodes per hidden layer used. Following scVI integration, latent spaces were passed in the *sc.pp.neighbors* (*n_neighbors=30*) and *sc.tl.umap* functions from Scanpy. In order to evaluate optimum scVI model and number of neighbourhoods, a varying number of latent spaces (10, 20, 30), batch sizes (1,024, 3,000, 5,000) and number of neighbours (15, 20, 25, 30) were scored for purity, homogeneity (*sklearn.metrics.homogeneity_score*), completeness (*sklearn.metrics.cluster.completeness_score*), normalised mutual information (*sklearn.metrics.cluster.normalized_mutual_info_score*) and global silhouette distance (*sklearn.metrics.silhouette_score*) for both biological replicate and chemistry. All metric outcomes were screened but global silhouette distance for both biological replicate and chemistry informed the final model and neighbourhood decision (20 latent spaces, 30 neighbours, batch size of 3,000). Finally, we utilised the uniform manifold approximation (UMAP) algorithm from the *sc.tl.umap* function from Scanpy for visualisation.

For annotation purposes, dimensionality reduction and integration as described above was independently re-run for cells and genes from each of 12 embryo grid-like anatomical tissue sections, and with the following differences: cell cycle genes were not removed, only top 6,000 HVG by dispersion were retained for scVI integration, default scVI parameters were utilised. For section-wise dimensional reduction, the optimised number of neighbours (between 15, 20, 25, and 30) were assessed using *sklearn.metrics* for silhouette scoring, with the best-performing number of neighbours (NNs) selected as follows: 15NNs for sections 2, 7, 11; 20NNs for sections 3, 6, 9; 25NNs for sections 1, 4, 10; 30NNs for sections 5, 8, 12.

### Whole embryo: scRNAseq manual annotation - clustering and differential expression testing

For manual annotation, clustering was performed independently on each of the 12 section-wise embryo scRNAseq objects using the Scanpy *sc.tl.leiden* function with a resolution parameter of *res=3.* Subclusters were then generated within each major cluster with a resolution parameter of *res=1*. Manual annotation was performed by a combination of literature markers, differentially expressed genes (DEGs) between major clusters using the *sc.tl.rank_genes_groups* function from Scanpy for a two-sided Wilcoxon rank-sum test (*corr_method=‘benjamini-hochberg’*) and information from the community data CellTypist model. DEG output was filtered for significant adjusted p-values (<0.05) and the top 100 significant DEGs per category were then ranked by log fold change. The resultant manual annotations were used for evaluation of QC metrics as described above (i.e., low quality cells according to mitochondrial reads, doublets, and maternal contaminants were removed), then these annotations were refined through the construction of a reference human embryonic dataset (described next).

### HDCA: Integration and annotation harmonisation of the final integrated community and whole embryo data

The HDCA dataset is the combination of the Whole Embryo and the Community data consisting of 4,679,782 cells, split into 12 level 2 systems based on the initial annotated cell types (**Fig. 1c**). For each of the systems, initial round of annotation harmonisation of public data and embryo data were performed as follows: 4,000 most highly variable genes (HVGs) were identified using *scanpy.pp.highly_variable_genes* with a concatenated variable of study, sample state (fresh or frozen), single cell or single-nuclei, and sequence technology as the batch key (*batch_key*). Batch correction was performed using scVI with default parameters using the above-mentioned batch key. kNN graphs were constructed using *scanpy.pp.neighbors* with k=15 and the scVI latent space. The underlying kNN was used to cluster the cells using Leiden community detection using *scanpy.tl.leiden* (*resolution* = 1.0).

For the aerodigestive, audiovisual, endocrine, cardiovascular, hematopoietic, integumentary, undifferentiated germ cells, yolk sac and urogenital systems, we adopted a cluster centric embryo-community label harmonisation approach. The 6PCW embryo cell types at level 3 (broad) and lvl4 (fine) were predicted using a distance weighted kNN classifier (*sklearn.neighbors.KNeighborsClassifier*) with k=10 nearest neighbours from the community data. Clusters that were largely dominated by cells from community data, the whole embryo cells of those clusters were assigned the kNN-classifier predicted cell types and clusters where cells were predominantly embryo data, the community data cells were assigned the whole embryo cell type. For clusters where both community and embryo data were proportionally well represented, we adopted a manual approach with expert guidance for cell type harmonisation.

Cells of the three distributed systems namely the mesenchyme (**Fig 3**), vascular and lymphatic endothelium (**Fig 4**) and peripheral nervous system (**Fig 5**) were of particular focus in our manuscript. Cells were assigned to these systems based on the initial round of refined cell type annotations derived from both community datasets and de novo data. Harmonisation of the annotations across the community data and the embryo data were then performed with expert guidance. For each of these three groupings the HVGs and cluster resolutions were not treated as a fixed or standard parameter as for other systems above, but instead were selected in a data-driven manner based on whether clusters represented biologically meaningful and sufficiently pure cell populations (res; vasculature = 4.0, peripheral nervous system = 1.0, mesenchyme = 3.0). For HVG used per system, please see **Supplementary Table 15**. Cell type identities were assigned using canonical marker genes (**Supplementary Table 9, 16**) and also informed by integrating the suspension and Xenium spatial data described below in section *Whole embryo: Xenium preprocessing and analysis*. After initially clustering and annotating each system independently, the refined cell types were collated and iteratively refined to assess and merge equivalent populations across systems. System-specific clustering also enabled us to more accurately resolve cells that had previously been assigned to broader categories. For example, within the mesenchymal system, higher-resolution clustering and annotation identified a subset of cells exhibiting vascular endothelial transcriptional profiles, which were subsequently reassigned to the vascular endothelial system. Clusters with less than 10 cells after refinement were assigned to k nearest neighbour’s celltype. For celltypes from the other systems, the initial unbiased clustering framework provided a common reference structure for harmonising the community datasets with the de novo Whole Embryo data and for evaluating concordance between existing annotations across studies. During cross-system refinement, previously published annotations were retained where they were unambiguous and provided higher-resolution labels than those obtained from unbiased clustering. For example, in the hematopoietic system, a myeloid lineage cluster in which the majority of cells were annotated as Macrophage_LYVE1_High by Suo Dann et al., but which also contained cells with more generic annotations such as Macrophage from Zhang et al., was annotated as MACROPHAGE_LYVE1_HIGH. Central nervous system cells retained the original fine-grained annotations from Braun et al., as these annotations were informed by spatial context, a key variable that would not otherwise be captured in the suspension transcriptomic latent space. The final annotation spans across 12 systems, 69 broad cell types and 454 refined cell types and are provided in **Supplementary Table 9.** Post harmonisation and cross system refinement, concordance between the HDCA refined cell type annotations and the original author-provided annotations was assessed using Normalised Mutual Information (NMI; *sklearn.metrics.normalized_mutual_info_score*). This yielded an NMI score of 0.85, indicating strong agreement between the two annotation schemes. The top 50 differentially expressed genes ranked by z-scores and padj < 0.05, for each of the refined cell types is provided in **Supplementary Table 16**. The refined cell types that were expert annotated in the three distributed systems (the mesenchyme, vascular and lymphatic endothelium and peripheral nervous system) are listed in **Supplementary Table 17.**

To integrate the entire public and embryo data after per-system annotation harmonisation as above, we used the above-mentioned *batch_key* to identify 12,288 HVGs. Highly variable genes were selected in a batch-aware manner using *scanpy.pp.highly_variable_genes*, with dispersion normalized within each batch and aggregated globally to mitigate batch-driven effects. The top 12,288 genes ranked by global normalized dispersion were retained for downstream analysis (**Supplementary Table 15**). We trained scVI using a two-layer network (2,048 hidden units per layer), a 28-dimensional latent space, a dropout rate of 0.1, and a zero-inflated negative binomial (ZINB) gene likelihood with gene-wise dispersion, observed library size, and a normal latent prior. Batch effects were accounted for by including the *batch_key* as a covariate in scVI training, to generate the latent space. The above parameters were selected following a limited hyperparameter search constrained by the size of the dataset. We evaluated combinations of hidden layer sizes (512, 1024, and 2048 units), latent dimensionality (26, 28, and 30), number of hidden layers (2 and 3), and likelihood models (negative binomial and zero-inflated negative binomial), using a dropout rate of 0.1. Model performance was assessed using scIB metrics computed on the initially predicted cell type annotations. The latent space was used to generate the kNN graph and this graph was used to generate the UMAP.

### scRNAseq gene expression and UMAP visualisation

To visualise gene expression across dotplots, heatmaps or UMAPs, raw gene expression count matrices were subset to cell types of interest, and were then normalised using the Scanpy *sc.pp.normalise_per_cell* function with counts_per_cell_after=1e4. A ln(x+1) transformation was then applied to the normalised counts using the *sc.pp.log1p* function. Gene expression was then visualised with either the *sc.pl.dotplot*, *sc.pl.heatmap* or *sc.pl.umap/sc.pl.embeddings* functions. To increase interpretability of plots, the *vmax* argument was sometimes used to apply an upper limit to the color scale or scaled using *standard_scale=’var’* (where used, noted within GitHub scripts).

### scRNAseq cluster entropy calculation

To assess donor diversity across the atlas (**Fig. S2b**), within each high-resolution leiden cluster (resolution = 3), we calculated a donor entropy score. Clusters with a greater mix of donors have higher entropy scores, while clusters dominated by a single donor have lower scores. Entropy was computed using the Shannon entropy formula (base-2 logarithm) based on the proportion of cells contributed by each donor within a cluster. Entropy scores were normalized to range from 0 to 1 for comparability across clusters with different numbers of donors. The normalized entropy scores were then mapped back to individual cells for visualization.

### scRNAseq cell-cell communication analysis

For the cross-system analysis within the Dorsal Root Ganglion (DRG), cell–cell communication analysis was performed using LIANA+^187,188^ v1.5.0 on the spatially resolved Xenium dataset restricted to DRG cells found in **Fig. S24d**. Ligand–receptor interactions were inferred using the *rank_aggregate* function, which aggregates evidence across the consensus resource from cells grouped by cell types. Only ligands and receptors expressed in at least 10% of cells within each group were considered (*expr_prop = 0.1*), with a minimum of 5 cells per cell type and 100 permutations used to assess interaction specificity; raw expression values were retained. To incorporate spatial information, pairwise mean Euclidean distances between all cell-type pairs were calculated from a precomputed cell–cell distance matrix. Interactions were further filtered to retain only those with statistically significant CellPhoneDB-derived p-values (≤ 0.05).

### scRNAseq gene set enrichment analysis

Gene set enrichment scores for the PNS system were computed using the Univariate Linear Model (ULM) method implemented in the decoupleR^189^ Python package (v2.1.1) in **Fig. 5m**, and **Fig. S23f, S23j, S24e**. For the enrichment of neural crest cells in skin organoids, curated marker gene signatures were used for NCC (*SOX10*, *FOXD3*), glial (*MPZ*, *PLP1*, *EDNRB*, *ERBB3*, *CDH19*), and neuronal (*ELAVL4*, *ELAVL3*, *PRPH*, *TUBB3*) transcriptional programs. To further resolve Schwann cell states, glial enrichment analyses additionally incorporated subtype-specific signatures for non-myelinating Schwann cells (*SCN7A*, *GFRA3*, *NCAM1*, *NRN1*) and myelinating Schwann cells (*MBP*, *PRX*, *PMP22*, *MAG*). To assess downstream receptor signalling activity, pathway gene sets were compiled from KEGG MEDICUS, Reactome, Gene Ontology Biological Process (GOBP), and BioCarta databases, including pathways associated with ‘WNT5A–ROR signalling’, ‘NCAM-mediated neurite outgrowth’, ‘L1CAM interactions’, ‘ERBB2 signalling’, ‘CXCR4 signalling’, and ‘HGF-MET-PI3K signalling’. In addition, ligand enrichment per niche was assessed using curated gene sets derived from GOBP and WikiPathways annotations, including ‘Neuron migration’, ‘Neuron maturation, Neural crest cell migration during development’, ‘Sympathetic ganglion development’ and ‘Peripheral nervous system neuron differentiation’. Enrichment scores were calculated at single-cell resolution to enable comparative analysis of lineage-specific programs and niche-derived ligand activity.

Arterial and venous module scores (**Fig. 4b**) were computed using the *sc.tl.score_genes* function in Scanpy. Arterial scores were calculated based on the gene set *CXCL12, GJA5, EFNB2, HEY1, DLL4, SOX17, NOTCH4*, and *NRP1*, whereas venous scores were derived from *TLL1, ADAMTS18, SELP, NR2F2, EPHB4, APLNR, VCAM1,* and *NTRK2*. Adult gene program (**Fig. 4c**) scores were computed using the same approach, using genes derived from lists of DEGs from Supplementary Table 6 of a previously published scRNA-seq atlas of human adult endothelial cells^94^. To ensure robustness, we performed 20 randomised subsampling iterations. In each iteration, cells were randomly sampled from vascular and lymphatic cell types excluding the query population, matching the total number of cells in the query group. Scores were recalculated for each subsample to assess stability and minimise sampling bias. Linear Generalised Additive Model (GAM) with four splines was used to fit the scores.

### Whole embryo: multiome sample acquisition and processing

Single nuclei were isolated from the remaining frozen cells from F137, F147 and F158 embryos that were not used for scRNAseq (https://www.protocols.io/view/nuclei-isolation-from-frozen-tissue-8epv5x6wjg1b/v1). Briefly, frozen cells were homogenised in homogenisation buffer (250 mM sucrose, 25 mM KCl, 5 mM MgCl2, 10 mM Tris-HCl, 1 μM dithiothreitol, 1× protease inhibitor, 0.4 U μl−1 RNaseIn, 0.2 U μl−1 SUPERaseIn and 0.1% Triton X-100 in nuclease-free water) using a pre-chilled Dounce homogeniser. Cells were left for 5 min on ice and then they were initially homogenised with a loose-fitting pestle “A” (4-8 strokes) followed by a tight-fitting pestle “B” (2-4 strokes). Homogenate was filtered through a 40μm cell strainer (CellTrics, Sysmex) and centrifuged for 6 min at 500 g (acceleration 0, deceleration 0) to obtain nuclei pellets. Pellets were resuspended in 500 ml of wash buffer (1× PBS, 5% BSA and 40 U μl−1 Protector RNaseIn) for 2 minutes on ice. After visualisation and assessment for viability under the microscope, nuclei suspensions were centrifuged again for 3 min at 500 g and the supernatant was carefully removed. Nuclei were manually counted and concentrations were estimated using an Improved Neubauer hemocytometer. All suspensions were maintained on ice until the transposition step.

### Whole embryo: multiome library preparation, sequencing and alignment

Nucleus suspensions were loaded at a concentration targeting 10,000 nuclei per channel. Nuclei were subjected to transposition before GEM generation and barcoding. All embryos were processed using the Chromium Next GEM Single Cell Multiome ATAC + Gene Expression kit (10x Genomics) according to the manufacturer’s instructions. Gene expression and ATAC libraries were prepared using proprietary 10x Genomics reagents and sequenced on the Illumina NovaSeq6000 S4, targeting a minimum depth of 50,000 UMIs per cell. Multiome data (sn-RNA-seq and sn-ATAC-seq) were aligned to GRCh38-2020-A reference using cellranger-arc v2.0.2 pipeline with the default parameters from 10X CellRanger suite.

### Whole embryo: multiome snRNAseq QC preprocessing, annotation and visualisation

Raw counts from whole embryo multiome single nuclear RNA-seq (sn-RNA-seq) samples (**Supplementary Table 5**) with 467,575 cells and 36,601 genes were processed using Scanpy v1.11.2^186^ (anndata v0.12.0) framework. Quality control metrics were calculated per cell, including total counts, and percentage of mitochondrial, ribosomal and hemoglobin counts. Cells were filtered to remove low-quality and doublets using the following thresholds:cells with >12,500, <200 detected genes and mitochondrial fractions > 5% were excluded. Genes expressed in fewer than 3 cells were removed. Counts were library-size normalised (shifted algorithm) using pp.normalize_total (target_sum=None), followed by log-transformation. Highly Variable Genes (HVGs) were identified using the Seurat v3 flavor with embryo_ID specified as the batch key, and the top 2,000 HGVs were retained. Dimensionality reduction was performed by PCA, followed by computing neighbours, UMAP and Leiden clustering (resolution parameter=1). Cells annotated as glia, neurons, neuronal progenitors were extracted based on established marker gene expression. The dataset was subset to these cells encompassing central nervous system (CNS) and Peripheral nervous system cells (PNS) populations for further subclustering. 3 Clusters (10, 12 and 13) were identified as PNS populations using known PNS markers and some CNS markers as negative markers (see Github) and were extracted for detailed annotation (**Fig. S8**). The PNS cells were subclustered in a new embedding space and leiden clustering of resolution = 0.5 identified 9 clusters corresponding to: Autonomic NCCs (Neuronal Crest cells)/ Schwann cell precursors (SCPs), Sensory NCCs/SCPs, sensory neurons, sensory neuronal progenitors, sensory neurons (epibranchial placode), sensory neurons (otic placode), Autonomic neuronal progenitors, NCC/SCP-derived progenitors, NCC/SCP-derived mesenchyme, Melanocyte precursors. Cluster identities were assigned using known lineage markers and differential expression analysis (**Fig. S8c**; **Supplementary Table 9**). Final cell type labels were retained for cross-modality integration with snATAC-seq data, enabled by shared barcodes from the 10x Genomics multiome technology.

### Whole embryo: multiome ATAC-seq preprocessing, peak calling and visualisation

snATAC-seq data were processed using cellranger-arc analytical suite and further filtered, and analysed downstream using SnapATAC2^190^ v2.8.0 framework. Cells were retained if they met the following criteria, >1,000 unique fragments and TSS enrichment score >=7 (stricter than the default threshold of 5). In order to investigate the dataset further and assess the batch effect across embryos in the ATAC modality we processed to feature selections and dimensionality reduction. Feature selection was performed using 50,000 genomics features prior to performing dimensionality reduction. For dimensionality reduction we used Laplacian Eigenmaps via spectral embedding implemented in the SnapATAC2 framework (snap.tl.spectral), followed by UMAP computation and visualisation with default parameters. After preprocessing and filtering, the whole embryo snATAC-seq dataset comprised 521,423 cells and 606,219 features. Since 10X multiome technology provides matched RNA and ATAC profiles from the same nuclei, snRNA-seq-derived annotations were directly transferred in ATAC modality using shared cell barcodes. From the full snATAC-seq dataset, 22,063 cells overlapped with PNS-annotated cell types defined in the snRNA-seq and were selected for downstream chromatin accessibility analysis. To ensure comparable peak detection across PNS cell populations, prior to peak calling cells were downsampled to a maximum of 2,000 cells per cell type. Cell types with less than 2,000 cells were not downsampled. This approach ensures more comparable and accurate peak identification across cell populations. Peak calling was performed using the MACS3^191^ method implemented in the snapATAC2. After peak calling, coverage tracks were generated and exported in bigWig format using snap.ex.export_coverage function. The genomic tracks were then visualised in the Interactive Genomics Viewer (IGV) to examine chromatin accessibility patterns at specific marker gene loci across PNS populations (**Fig. S8**).

### Whole embryo: multiome cell2fate analysis

We quantified spliced and unspliced counts with Velocyto (version v0.17.17) using GRCh38 v3. RNA velocity modules were separately modelled for PNS sensory and autonomic lineages using level 3 cell type annotations with cell2fate^153^ (v.1.0.0, https://github.com/BayraktarLab/cell2fate) for **Fig. S23e**. Cell types outside lineages or with low cell numbers (parasympathetic neuronal, enteric glia, chromaffin cells, melanocyte precursors) were not considered for analysis. We used 3,000 highly-variable protein-coding genes (19,273 genes, GRCh38, HUGO Gene Nomenclature Committee, 2025-10-02 version) with at least 10 detected counts, implementing the default dynamical model with embryo as batch key (“haniffa_ID”) and 8 modules. Model training was performed on an NVIDIA A100-SXM4-80GB GPU for 200 epochs until convergence with a default learning rate of 0,01. Module time estimates and activation/repression states were sampled from the posterior distribution (default 30 samples), and module-specific top genes, TFs, and GO-terms were calculated using *get_module_top_features* function.

### Whole embryo: Visium sample processing

A single embryo at Carnegie stage 17 (6 post conception weeks; PCW), sample SOB26, was formalin fixed and embedded in a paraffin (FFPE) block (**Supplementary Tables 1, 5**). One section at 4μm thickness was taken along the sagittal plane through the centre of the embryo (section_121) and through the left upper and lower limbs (section_221) using a microtome and collected on Superfrost plus microscope slides. 2 slides (section_121, section_221) were then dried for 3 hours at 42°C then placed in desiccant to avoid moisture. First, deparaffinization and Haematoxylin and Eosin staining were performed according to the standard 10X CytAssist workflow (CG000520). Slides were then imaged at 40X magnification on a Hamamatsu Nanozoomer. These slides were then taken forward for Visium analysis by destaining and decrosslinking the stained sections according to the standard 10X CytAssist workflow (CG000520) then proceeding immediately to perform the Visium CytAssist Spatial Gene Expression according to the standard 10X CytAssist workflow (CG000495). In brief this includes, probe hybridisation and probe ligation followed by CytAssist enabled probe release and extension before pre-amplification and cleanup. For these samples, the Visium_Human_Transcriptome_Probe_Set_v2.0_GRCh38-2020-A was used. Next, the samples were taken forward for probe-based library construction and quality control was performed using the Agilent Bioanalyzer High Sensitivity chip. 4 libraries were pooled at a time and sequenced down one lane of an Illumina Novaseq SP flow cell with the following run parameters: read 1: 28 cycles; i7 index: 10 cycles; i5 index: 10 cycles; read 2S: 50 cycles.

### Whole embryo: manual annotation of H&E images, and visualisation on Visium spatial data

Manual annotations of H&E section_121 from embryo SOB26 were performed by the Human Developmental Biology Resource (HDBR) team using the OMERO.iviewer (v0.16.0) platform. The original high-resolution images were hosted on the OMERO Plus server (https://omeroplus.sanger.ac.uk/) provided by Glencoe Software and maintained by the CellGenIT group at the Wellcome Sanger Institute. Regions of interest (ROIs) were outlined using the Draw Polygon tool within OMERO.iviewer to assign labels to known anatomical features.

The resulting annotations were subsequently integrated with Visium spatial transcriptomics data using the ann2SR.py script from the OMERO_tools repository (https://github.com/cellgeni/OMERO_tools). This script retrieves the polygon coordinates of annotated regions from the OMERO server and combines them with the spatial coordinates of Visium spots. Based on geometric overlap, each Visium spot is assigned to annotated anatomical regions in both a categorical manner (discrete region label) and a continuous manner (proportional overlap with annotated regions) (**Fig. S7b**). The script outputs a CSV file containing spot (cell) identifiers together with the corresponding annotation assignments.

### Whole embryo: Visium gene expression analysis

Spatial transcriptomics Visium data (section_121) was mapped using SpaceRanger (v2.1.0) using GRCh38-2020-A reference. Spots were filtered to retain spots with total RNA counts >2000 and <200,000. Low count genes were then removed using the *cell2location.utils.filtering.filter_genes* function with the following parameters: cell_count_cutoff=5, cell_percentage_cutoff2=0.15, nonz_mean_cutoff=1.11. Genes were filtered to the set shared with scRNAseq data. This yielded Visium data comprising 2 sections (section_121 and section_221) with 9,043 spots, and 17,824 genes. Only section_121 was used for downstream analysis comprising 7,778 spots. For plotting gene expression within the DRG region per Visium voxel (**Fig. 5j**) the spots corresponding to the DRG region were manually drawn as a region of interest (ROI) using the 10x Genomics Loupe Browser and the spatial coordinates of selected DRG spots were exported as CSV files. From the selected spots, the x and y centroid coordinates of each spot in pixel units (high resolution) were converted to micrometres (µm) using the Visium platform scale factor to represent true physical distance in tissue space.For the distance calculation from the DRG spots, the spatial proximity between spots was calculated by constructing a *k*-dimensional tree (k-d tree) using the full resolution spatial coordinates of DRG spots and *scipy.spatial.cKDTree(k=1) function*. The nearest Euclidean distance between each spot in the section_121 to its nearest DRG_spots was computed by querying each against the constructed KDTree and resulting distance metric for each spot (in µm) was stored as a per-spot metadata variable. To visualise the results, mean expression of each gene set was aggregated per cell and binned by distance from the DRG across a fixed number of equally spaced bins (n=10); per-bin means and standard errors were computed for each sample, then smoothed using a centered rolling window (window size=5) to produce continuous expression-by-distance profiles with 95% confidence intervals.

### Whole embryo: Xenium sample acquisition and processing

FFPE tissue samples from sample SOB25 (Carnegie stage 16) and SOB26 (Carnegie stage 17) were sectioned onto Xenium slides (10x Genomics 3000941) at a thickness of 4µm and stored at RT in desiccant (**Supplementary Table 1, 5**). These were then processed with the Xenium Prime workflow as per the manufacturer’s protocols: CG000580 for pretreatment including deparaffinization and decrosslinking; and CG000760 for probe hybridisation, rolling circle product generation, segmentation marker staining, autofluorescence quenching, and nuclear counterstaining. Slides were loaded onto a Xenium Analyzer for automated cyclic fluorescent labelling and imaging as per the manufacturer’s protocol CG000584, with Xenium Onboard Analysis software v3.0.0.15 or v3.1.0.4. Data were exported in the vendor format of ome.tif images (segmentation stain images including DAPI-stained nuclei), .h5 cell-gene count matrices, and .csv cell and transcript data.

### Whole embryo: Xenium preprocessing and analysis

We used default 10x cell segmentation, utilising segmentation marker staining. 10x Genomics Xenium data was filtered to exclude cells with <10 genes per cell. We then used SCANVI (number of layers: 1; number of latent dimensions: 10) using labelled scRNA-seq and unlabelled Xenium data, restricted to ∼5000 genes found in both datasets. Batch-aware HVG selection was then performed by setting the batch key to sample ID. We selected 2,000 HVGs as features for the integration. The batch key was sample id and a categorical covariate of technology (scRNA vs spatial) was used. We used this initial integration to broadly label groups in Xenium data (e.g. vascular, peripheral nervous system). We then repeated feature (HVG) selection and integration by lineage. For each lineage, we calculated the kNN graph and performed Leiden clustering. We labelled clusters based on a combination of scRNA-seq annotations and spatial location, with spatial positions helping to refine names for populations. We defined *FRZB*^+^ progenitors as populations that had an uncertain identity after expert annotation. In contrast, other progenitors, such as dermal fibroblast progenitors and limb fibroblast progenitors, had been assigned a clear identity after expert annotation. For the predominantly mesenchymal cluster, we repeated the by-lineage integration separately for each group due to the increased complexity of this group (>3 million cells overall). These joint integrations, and the same marker genes used for scRNA-seq and Xenium-5k data, are shown in **Appendix B**.

### Whole embryo: Niche identification analysis

We ran NicheCompass^51^ (v0.3.0), a graph neural network-based approach, to obtain niche embeddings. We split whole embryo sections into 8 reference and 3 query sections (**Supplementary Table 5)**, to illustrate how our reference can be used to obtain niche embeddings for future datasets. Spatial neighbourhood graphs were constructed using Squidpy (8 nearest neighbours), and curated gene programs were assembled using NicheCompass. NicheCompass was trained on the reference dataset with default parameters and batch key as sample_id for 100 epochs. Niche annotations were based on histopathological annotations of whole embryos and differentially expressed genes (**Supplementary Table 12**), calculated after log1p normalisation using Scanpy’s rank_genes_groups function with a t-test. We also used niche cell compositions predicted from SCANVI. For query data, the trained model was subsequently fine-tuned on the query data and both datasets were integrated into a shared latent space. All weights were frozen except categorical covariates embedder weights (unfreeze_cat_covariates_embedder_weights == True). Fine-tuning was otherwise performed using the same training objective and hyperparameters as reference training. Low-dimensional embeddings were computed using UMAP after kNN construction (k=15). For querying disease genes across niches, we used a reported gene list for Mendelian genes for congenital cataracts^75^.

### HDCA mapping to skin organoid for mesenchyme analysis

We jointly integrated human skin organoid (unlabelled for integration) with our HDCA using scanVI. We included only populations labelled as fibroblast: organoid_Fibroblasts POSTN+, organoid_Pre-dermal condensate, organoid_Early fibroblasts FRZB+, organoid_Early fibroblasts HOXC5+, organoid_Dermal condensate, organoid_Dermal papilla, organoid_Adipocytes, organoid_Neuron progenitors, organoid_Neuron progenitors SPP1+, organoid_Neuroepithelial cells, organoid_Smooth muscle/Pericytes, organoid_Myoblasts, organoid_Myocytes, organoid_Early myocytes. We included samples for HVG selection (∼2000 target genes in a batch-aware manner, using sample id as batch key) if they contain >1000 cells. Genes associated with mitochondrial, ribosomal, hemoglobin, cell-cycle, hypoxia, and stress responses were removed. kNN graph construction (k=20) was performed followed by Leiden clustering. We identified Leiden clusters enriched for progenitor populations, subset the dataset to these clusters, and recomputed the k-nearest neighbor graph in the SCANVI latent space. UMAP embeddings were initialized using the PAGA graph to preserve global trajectory structure. SCANVI was used to transfer new annotations from reference atlases to organoid data, which we validated using the same marker genes used for the HDCA populations

### Whole embryo: RNAscope in situ hybridization imaging

Human embryonic tissues for RNAScope were firstly fixed in 10% PFA overnight at room temperature, and then secondly fixed in Methacarn before they were processed using automated Excelsior AS (Thermo Scientific, A823-1005) to generate FFPE blocks. Sections were cut from the blocks at 8 µm using a manual rotary microtome (Leica microsystems RM2235, UK). Simultaneous RNA *in situ* hybridization was carried out using RNAscope Multiplex Fluorescent Reagent Kit v2 (Cat. No. 323100 and 323270, ACD) following the manufacturer’s protocol for FFPE tissue mounted on slides, with pre-treatment steps of Hydrogen Peroxide and Protease reagents (Cat. No. 322381, ACD) and target retrieval reagents for 20mins at 95°C (Cat. No. 322000, ACD). The following RNAscope probes were designed and purchased from ACD: Hs-PIEZO1 (Cat No. 485101-C3), Hs-PROX1 (Cat No. 530241) and VEGFR3 (Hs-FLT4) (Cat No. 552441-C2). The probes were fluorescently labelled using OPAL 520 (Cat. No. FP1487001KT, PerkinElmer), OPAL 570 (Cat. No. FP1488001KT, PerkinElmer) and OPAL 650 dyes (Cat. No. FP1496001KT, PerkinElmer) for direct visualization under automated laser-scanning microscopy (Zeiss CellDiscoverer7, Carl Zeiss, Germany) (**Fig. 4l**).

### Whole embryo: RareCyte immunofluorescence staining and imaging

The following steps were performed at room temperature unless otherwise noted. Formalin fixed tissue from embryo sample SOB26 (**Supplementary Table 1**) embedded in paraffin (FFPE) were sectioned at 4μm thickness with a microtome (Leica RM2235) and mounted onto superfrost slides (Fisher Scientific 12312148) before drying at 60°C for 60 minutes to ensure adhesion. The mounted sections were then deparaffinized and antigen retrieval was performed using the BioGenex EZ-Retriever system for 95℃ for 5 minutes then 107℃ for 5 minutes. Slides were then bleached with AF Quench Buffer (4.5% H2O2 / 24 mM NaOH in PBS) to remove autofluorescence. Quenching was performed using a UV transilluminator using the high setting for 60 minutes with a strong white light exposure followed by 30 minutes using the 365 nm high setting. Slides were then rinsed with 1X PBS before incubating in 300 µl of Image-iT™ FX Signal Enhancer (Thermo Fisher I36933) for 15 min and rinsed again. The primary labelled antibodies Ki67 (RareCyte, 52-1013-501, fluorophore ArgoFluor 555L, clone D3B5) and Pan-CK (RareCyte, 52-1015-801, fluorophore ArgoFluor 845, clone C11/AE1/AE3) were prediluted according to manufacturer’s recommendations and added to the tissue then incubated for 120 minutes in the dark on a humidity tray. The antibodies used are provided in **Supplementary Table 3**. Slides were then washed with a surfactant wash buffer then 300 µl of nuclear staining in goat diluent was added to the slide, before incubating in the dark for 30 minutes on a humidity tray. Slides were washed and placed in 1X PBS before adding a coverslip with ArgoFluor mount media and left in the dark at room temperature overnight to dry. For imaging, the RareCyte Orion microscope was used with a 20X objective lens (**Fig. S1f**). Scans were performed using Imager and relevant acquisition settings were applied using the software Artemis. Slides were subsequently transferred to −20°C for extended storage.

### Whole embryo: MACSIMA immunofluorescence staining and imaging

For MACSIMA immunofluorescence imaging of the dorsal root ganglion (**Fig. 5i; Supplementary Table 1**), a 7PCW human embryonic (sample SM74P7) was cryoprotected in a solution of 10% sucrose at 4°C for 3 days. It was then embedded in a mixture of 7.5% gelatin, 10% sucrose, and 0.12 M phosphate buffer, followed by freezing in isopentane at −55°C and storage at −80°C until needed. The sample was cut into 10μm sections using a cryostat (Leica CM3050) and placed on Superfrost slides for frozen sections. Immunofluorescence staining was performed using the MACSima™ Imaging Platform. This imaging system operates by iterative immunofluorescent staining, sample washing, imaging, and signal bleaching, using 3 fluorochrome-conjugated antibodies per cycle. The antibodies used are provided in **Supplementary Table 3**. Image datasets (stack of tiff images) were analyzed using the MACS iQ View Software Version 1.3.2.

### HDCA web portal integration with developmental disorders database

We downloaded the DDG2P^43^ database from the G2P website (https://www.ebi.ac.uk/gene2phenotype, accessed on 16/10/2024). DDG2P gene-disease records are annotated with a confidence level which reflects the level of evidence supporting the association. Records with ‘limited’ confidence were removed, keeping only associations with ‘definitive’, ‘strong’ and ‘moderate’ confidence. We then stored gene-disease pairs in a relational database and used the open-source software Strapi to expose a web API to allow our Cherita web portal to query and retrieve the data dynamically by either gene or disease as per user exploration. Cherita, derived from the Malay word for “story”, embodies the core purpose of the tool: enabling users to derive biological insights through their data. For more detailed information about its functionality and applications, please refer to the GitHub repository at https://github.com/haniffalab/cherita-react.

## Acknowledgements

We are grateful to the donors and donor families for granting access to the tissue samples. We thank the Newcastle University Flow Cytometry Core Facility, Newcastle University Genomics Facility, Sanger Institute Cellular Genetics IT, CRUK CI Genomics Core Facility, and Newcastle upon Tyne NHS Trust NovoPath. We thank Mr Mohi Miah for wet-lab tissue processing assistance. We thank Dr Aidan Maartens and Dr Vicki Moignard for their writing contributions and suggestions. We also thank Martin Prete for his technical guidance on the use of the OMERO platform for anatomical annotations of H&Es.

## Author contributions

Conceptualisation: M.H. Project Administration: S.W, MS.V.B, V.R. Formal Analysis: MS.V.B, S.W, A.R, C.X, L.S, M.A.K, K.I, D.J, A.V.Po, MA.J, K.T, K.Ra, S.Mad, J.T.H.L, S. Mak, J.J, K.P.C, L.D, G.DA. Data Curation: MS.V.B, S.W, A.R, C.X, L.S, M.A.K, K.I, D.J, K. Ra, A.V. Po, E.K, M.A.C, J.J. Investigation: E.S, NJ.C, J.B, A.A, S.MA, R.A.B, K.Ro, B.O, C.H, J.E, E.P, N.H.G, D.M. Visualisation: E.S, A.R.F, D.B.L, C.A, E.K, M.A.C, H.V.F, C.F.W. Data Interpretation: S.W, A.R, C.X, L.S, M.A.K, E.S, K.I, D.J, A.R.F, NJ.C, A.V.Po, MA.J, K.T, K.Ra, J.K, A.L, J.E.G.L, V.K, S.L, J.T.H.L, A.A, E.U, T.P.S, V.L, F.T, R.A.B, B.O, A.M.C, H.J.W, N.H.G, P.H, M.W.M, B.T, L.F, D.P, C.M, R.E, S. Ba, V.R.K.S, K.K, J.C, S.S, I.S, M.D.L, S.MA, P.O, O.P, L.G, P.A, V.B.MS, F.J.T, F.P. Software: D.B.L, D.H, A.R, C.X, L.S, J.F, T.L, K.P, A.V.Pr, P.M. M.L. Resources: S.L, M.C, A.G, A.F, J.K, R.V.T, A.C, D.T, N.K.W, J.M. Funding Acquisition: M.H, O.A.B, S.A.T, K.B.M, B.G, E.R, S.Be. Supervision: M.H, O.A.B, F.P, F.J.T, S.A.T, K.B.M. Writing – Original Draft Preparation: S.W, C.X, L.S, D.J, F.P. Writing - Review & Editing: S.W, M.H. All authors reviewed and approved the manuscript.

## Data and code availability

### Data

Raw de novo whole embryo data will be available on EMBL-EBI ArrayExpress and BioImage Archive. All data accessions are currently private, and will be made available upon publication:

- scRNAseq raw FASTQ and counts (embryo F137), E-MTAB-11504
- scRNAseq raw FASTQ and counts (embryo F147), E-MTAB-11520
- scRNAseq raw FASTQ and counts (embryo F158), E-MTAB-11911
- Multiome raw FASTQ and counts (embryos F137, F147, F158), E-MTAB-16757
- Spatial transcriptomics Visium raw FASTQs, counts, images (embryo SOB26), E-MTAB-14785
- Spatial transcriptomics Xenium raw cell x gene counts post Xenium cell segmentation, images (embryos SOB25, SOB26), BioImage Archive, S-BIAD2878

The HDCA atlas will be accessible via an interactive web portal at https://cellatlas.io/studies/hdca/. This webportal is currently private and password protected, and will be made available upon publication. Through this portal, users will be able to visualise the dataset and download both raw and processed count matrices, as well as associated metadata and models.

### Code

All code will be made publicly available at https://github.com/haniffalab/hdca-paper; the repository is currently private, and will be made available upon publication.

Additional repositories associated with this study include https://github.com/theislab/idtrack^181^ for biological identifier mapping and harmonization across heterogeneous data sources, and https://github.com/theislab/sctram^45^, which provides the trajectory-fidelity benchmarking framework introduced in this study.

