## Appendix A for "An integrated single cell and spatial omics atlas of human prenatal development" (from another version of this preprint: version 1, 2026-04-01)

### Appendix A – Central nervous system (CNS) niches

**Appendix A Fig. 1 Central nervous system niche overview.** a) Dot plot of marker genes for CNS niches. b) Whole embryo sections coloured by CNS niches for reference and query sections (Xenium 5k profiling).

**Appendix B Fig. 2 – Central nervous system niches in section 28.** a) Whole embryo section 28 coloured by CNS niches. b) Log1p norm gene expression for selected marker genes.

**Appendix B Fig. 3 – Central nervous system niches in section 47.** a) Whole embryo section 47 coloured by CNS niches. b) Log1p norm gene expression for selected marker genes.

**Appendix B Fig. 4 – Central nervous system niches in section 69.** a) Whole embryo section 69 coloured by CNS niches. b) Log1p norm gene expression for selected marker genes.

**Appendix B Fig. 5 – Central nervous system niches in section 103.** a) Whole embryo section 103 coloured by CNS niches. b) Log1p norm gene expression for selected marker genes.

Note: Here CNS niches are defined in an unsupervised manner based on an index cell and its nearest neighbours using NicheCompass. This is unsupervised and does not use cell type labels or H&E annotations when generating niche embeddings.

H&E annotations and differentially expressed genes for niches were used to aid annotation (assignment of labels to niche clusters). Notably, there were some differences of unsupervised niches with H&E annotations. For instance, the telencephalon was often annotated as a single complete structure, but for unsupervised niches we observed *FOXG1*$^{lo}$ regions more consistent with diencephalon +/or migratory neurones. We show these patterns across multiple sections, showing robustness of this finding. Additionally, we also show gene expression spatial maps. For instance, for the telencephalon region that was suggested to be a distinct niche, we show *FOXG1*$^{lo}$ regions. We also show expression of genes in consistent patterns (e.g. FEZF1, SORCS2, PAX6, EMX2) around the migratory region between telencephalon and diencephalon. We thus propose the niches as an additional way of understanding CNS regions in addition to H&E. These niches can be queried for specific gene expression across ~5000 genes in early human development.

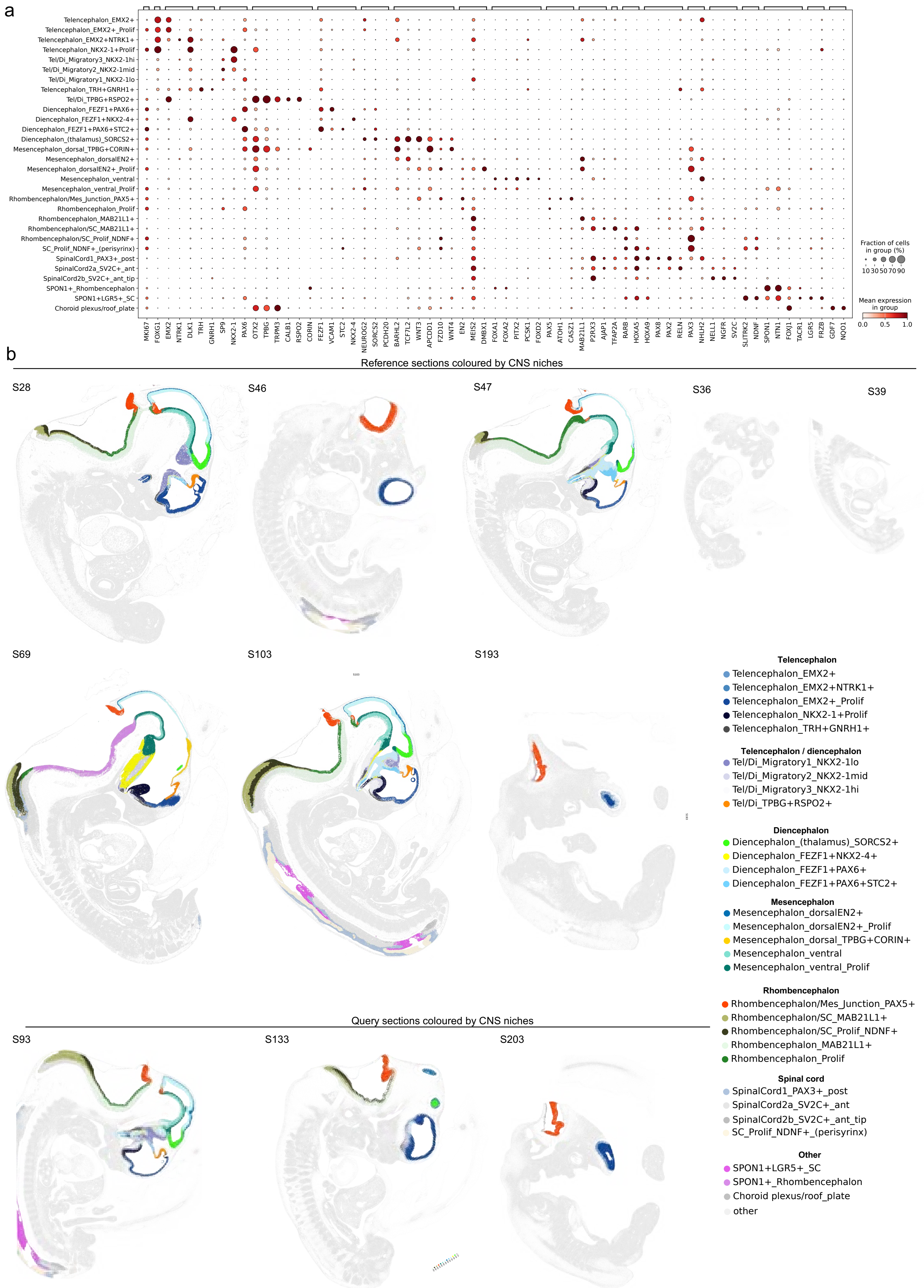

a
Telencephalon_EMX2+
Telencephalon_EMX2+_Prolif
Telencephalon_EMX2+NTRK1+
Telencephalon_NKX2-1+Prolif
Tel/Di_Migratory3_NKX2-1hi
Tel/Di_Migratory2_NKX2-1mid
Tel/Di_Migratory1_NKX2-1lo
Telencephalon_TRH+GNRH1+
Tel/Di_TPBG+RSPO2+
Diencephalon_FEZF1+PAX6+
Diencephalon_FEZF1+NKX2-4+
Diencephalon_FEZF1+PAX6+STC2+
Diencephalon_(thalamus)_SORCS2+
Mesencephalon_dorsal_TPBG+CORIN+
Mesencephalon_dorsalEN2+
Mesencephalon_dorsalEN2+_Prolif
Mesencephalon_ventral
Mesencephalon_ventral_Prolif
Rhombencephalon/Mes_Junction_PAX5+
Rhombencephalon_Prolif
Rhombencephalon_MAB21L1+
Rhombencephalon/SC_MAB21L1+
Rhombencephalon/SC_Prolif_NDNF+
SC_Prolif_NDNF+_(perisyrinx)
SpinalCord1_PAX3+_post
SpinalCord2a_SV2C+_ant
SpinalCord2b_SV2C+_ant_tip
SPON1+_Rhombencephalon
SPON1+LGR5+_SC
Choroid plexus/roof_plate
MKI67 FOXG1 EMX2 NTRK1 DLK1 TRH GNRH1 SP9 NKX2-1 PAX6 OTX2 TPBG TRPM3 CALB1 RSPO2 CORIN FEZF1 VCAM1 STC2 NKX2-4 NEUROG2 SORCS2 PCDH20 BARHL2 TCF7L2 WNT3 APCDD1 FZD10 WNT4 EN2 MEIS2 DMBX1 FOXA1 FOXA2 PITX2 PCSK1 FOXD2 PAX5 ATOH1 CASZ1 MAB21L1 P2RX3 AJAP1 TFAP2A RARB HOXA5 HOXA9 PAX8 PAX2 RELN PAX3 NHLH2 NELL1 NGFR SV2C SLITRK2 NDNF SPON1 NTN1 FOXJ1 TACR1 LGR5 FRZB GDF7 NQO1
Fraction of cells in group (%)
10 30 50 70 90
Mean expression in group
0.0 0.5 1.0
b
Reference sections coloured by CNS niches
S28
S46
S47
S36
S39
S69
S103
S193
Query sections coloured by CNS niches
S93
S133
S203
Telencephalon
Telencephalon_EMX2+
Telencephalon_EMX2+NTRK1+
Telencephalon_EMX2+_Prolif
Telencephalon_NKX2-1+Prolif
Telencephalon_TRH+GNRH1+
Telencephalon / diencephalon
Tel/Di_Migratory1_NKX2-1lo
Tel/Di_Migratory2_NKX2-1mid
Tel/Di_Migratory3_NKX2-1hi
Tel/Di_TPBG+RSPO2+
Diencephalon
Diencephalon_(thalamus)_SORCS2+
Diencephalon_FEZF1+NKX2-4+
Diencephalon_FEZF1+PAX6+
Diencephalon_FEZF1+PAX6+STC2+
Mesencephalon
Mesencephalon_dorsalEN2+
Mesencephalon_dorsalEN2+_Prolif
Mesencephalon_dorsal_TPBG+CORIN+
Mesencephalon_ventral
Mesencephalon_ventral_Prolif
Rhombencephalon
Rhombencephalon/Mes_Junction_PAX5+
Rhombencephalon/SC_MAB21L1+
Rhombencephalon/SC_Prolif_NDNF+
Rhombencephalon_MAB21L1+
Rhombencephalon_Prolif
Spinal cord
SpinalCord1_PAX3+_post
SpinalCord2a_SV2C+_ant
SpinalCord2b_SV2C+_ant_tip
SC_Prolif_NDNF+_(perisyrinx)
Other
SPON1+LGR5+_SC
SPON1+_Rhombencephalon
Choroid plexus/roof_plate
other

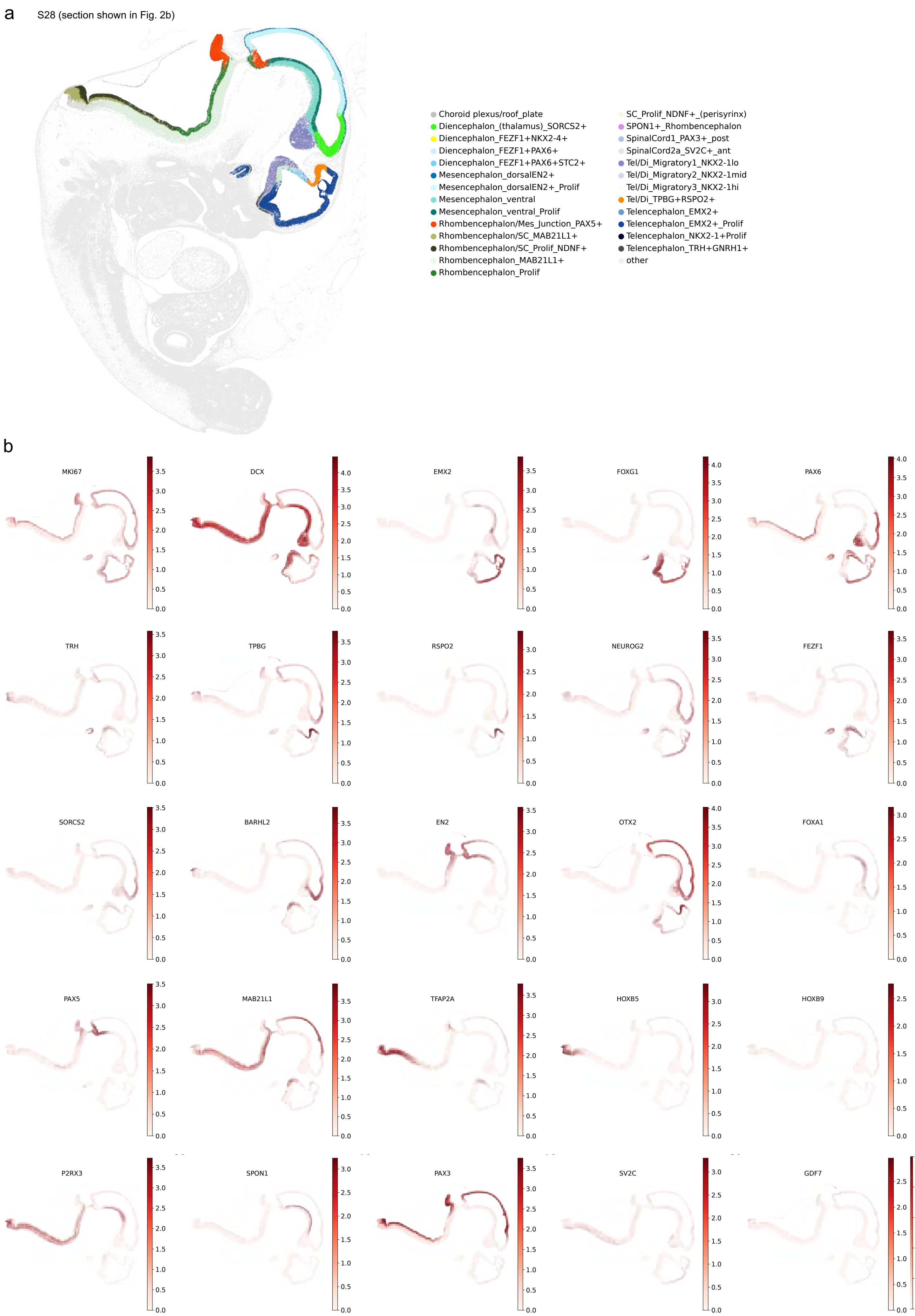
a
S28 (section shown in Fig. 2b)
Choroid plexus/roof_plate
Diencephalon_(thalamus)_SORCS2+
Diencephalon_FEZF1+NKX2-4+
Diencephalon_FEZF1+PAX6+
Diencephalon_FEZF1+PAX6+STC2+
Mesencephalon_dorsalEN2+
Mesencephalon_dorsalEN2+_Prolif
Mesencephalon_ventral
Mesencephalon_ventral_Prolif
Rhombencephalon/Mes_Junction_PAX5+
Rhombencephalon/SC_MAB21L1+
Rhombencephalon/SC_Prolif_NDNF+
Rhombencephalon_MAB21L1+
Rhombencephalon_Prolif
SC_Prolif_NDNF+_(perisyrinx)
SPON1+_Rhombencephalon
SpinalCord1_PAX3+_post
SpinalCord2a_SV2C+_ant
Tel/Di_Migratory1_NKX2-1lo
Tel/Di_Migratory2_NKX2-1mid
Tel/Di_Migratory3_NKX2-1hi
Tel/Di_TPBG+RSPO2+
Telencephalon_EMX2+
Telencephalon_EMX2+_Prolif
Telencephalon_NKX2-1+Prolif
Telencephalon_TRH+GNRH1+
other
b
MKI67
DCX
EMX2
FOXG1
PAX6
TRH
TPBG
RSPO2
NEUROG2
FEZF1
SORCS2
BARHL2
EN2
OTX2
FOXA1
PAX5
MAB21L1
TFAP2A
HOXB5
HOXB9
P2RX3
SPON1
PAX3
SV2C
GDF7

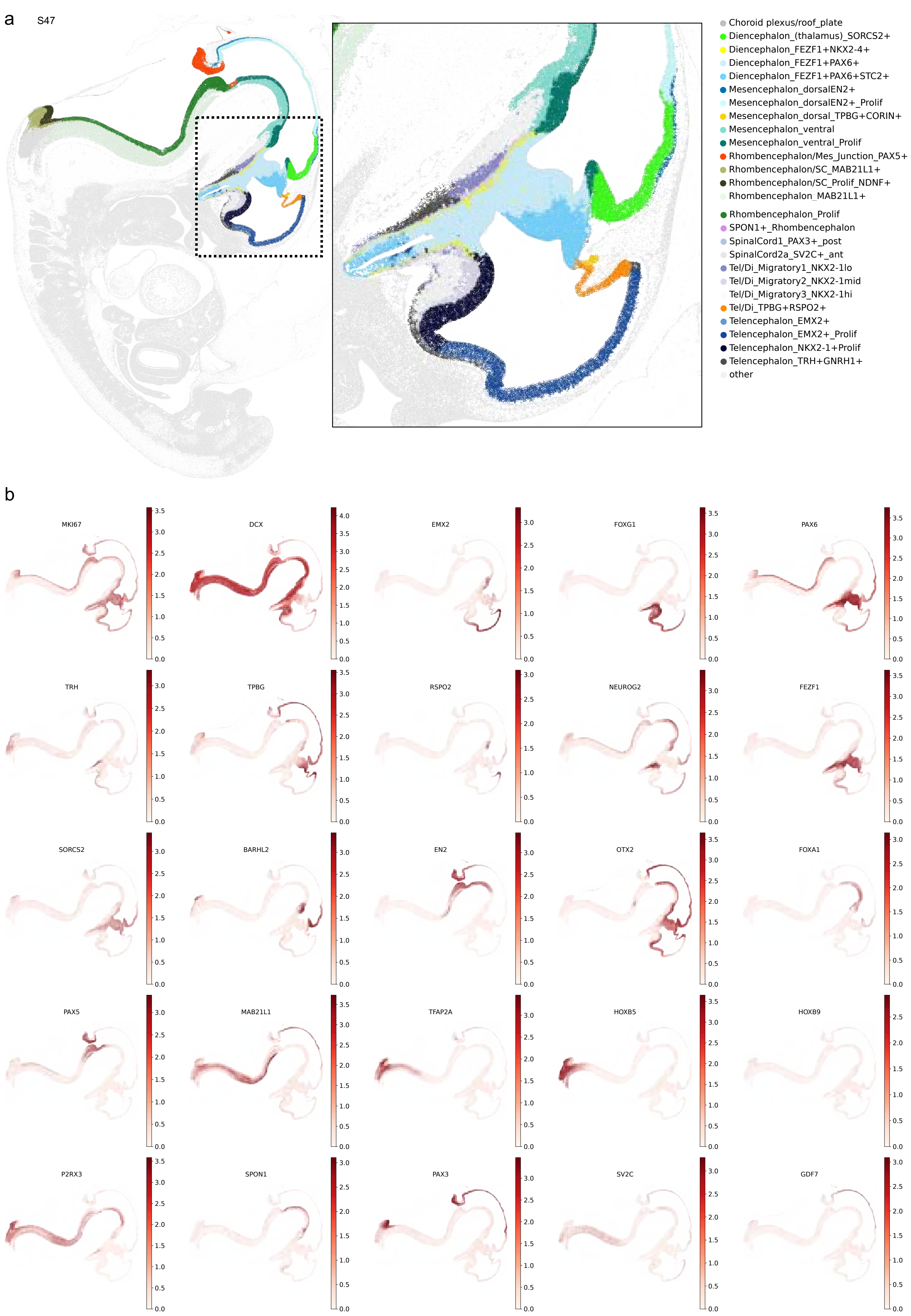

a
S47
Choroid plexus/roof_plate
Diencephalon_(thalamus)_SORCS2+
Diencephalon_FEZF1+NKX2-4+
Diencephalon_FEZF1+PAX6+
Diencephalon_FEZF1+PAX6+STC2+
Mesencephalon_dorsalEN2+
Mesencephalon_dorsalEN2+_Prolif
Mesencephalon_dorsal_TPBG+CORIN+
Mesencephalon_ventral
Mesencephalon_ventral_Prolif
Rhombencephalon/Mes_Junction_PAX5+
Rhombencephalon/SC_MAB21L1+
Rhombencephalon/SC_Prolif_NDNF+
Rhombencephalon_MAB21L1+
Rhombencephalon_Prolif
SPON1+_Rhombencephalon
SpinalCord1_PAX3+_post
SpinalCord2a_SV2C+_ant
Tel/Di_Migratory1_NKX2-1lo
Tel/Di_Migratory2_NKX2-1mid
Tel/Di_Migratory3_NKX2-1hi
Tel/Di_TPBG+RSPO2+
Telencephalon_EMX2+
Telencephalon_EMX2+_Prolif
Telencephalon_NKX2-1+Prolif
Telencephalon_TRH+GNRH1+
other
b
MKI67
DCX
EMX2
FOXG1
PAX6
TRH
TPBG
RSPO2
NEUROG2
FEZF1
SORCS2
BARHL2
EN2
OTX2
FOXA1
PAX5
MAB21L1
TFAP2A
HOXB5
HOXB9
P2RX3
SPON1
PAX3
SV2C
GDF7

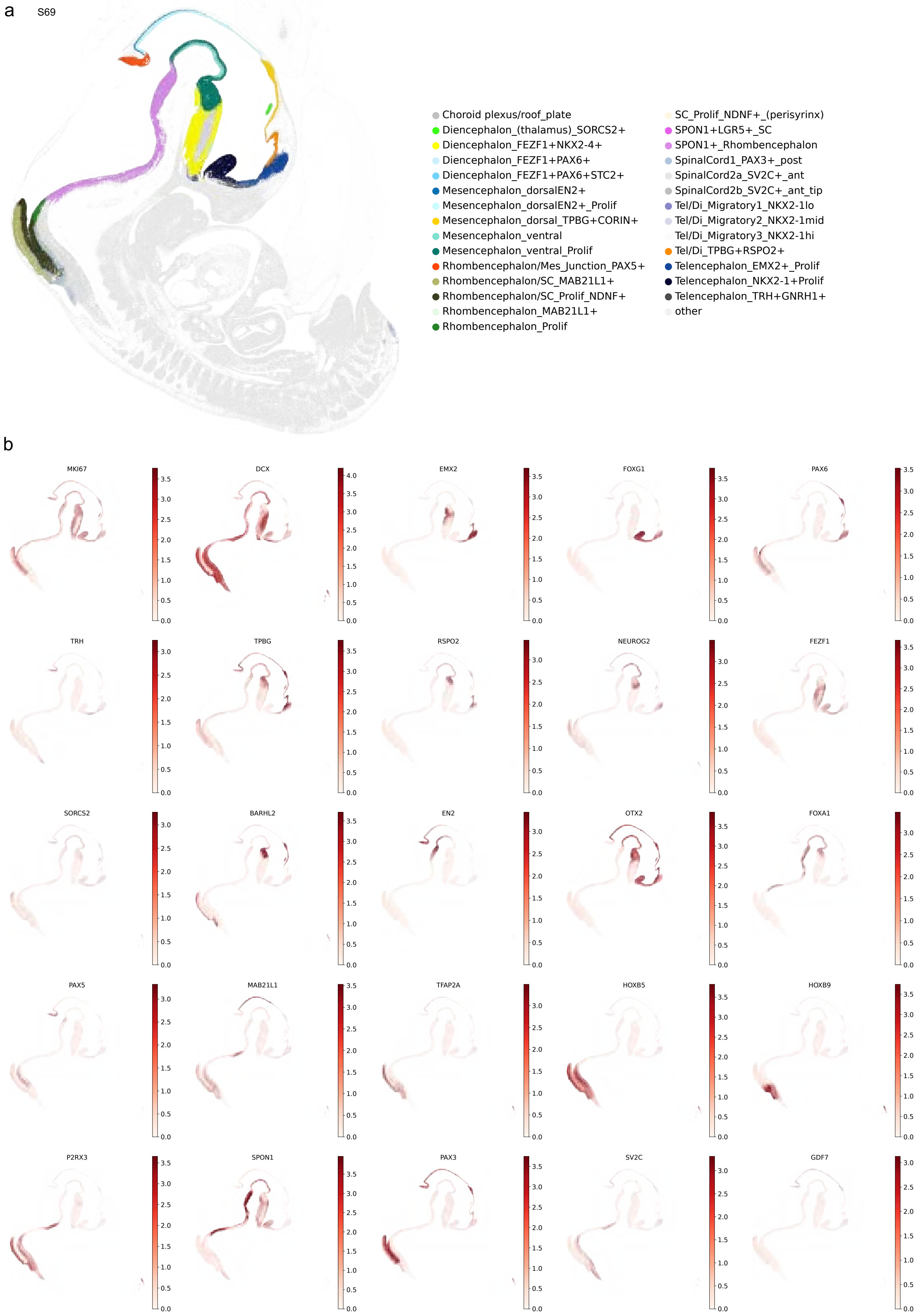
a
S69
Choroid plexus/roof_plate
Diencephalon_(thalamus)_SORCS2+
Diencephalon_FEZF1+NKX2-4+
Diencephalon_FEZF1+PAX6+
Diencephalon_FEZF1+PAX6+STC2+
Mesencephalon_dorsalEN2+
Mesencephalon_dorsalEN2+_Prolif
Mesencephalon_dorsal_TPBG+CORIN+
Mesencephalon_ventral
Mesencephalon_ventral_Prolif
Rhombencephalon/Mes_Junction_PAX5+
Rhombencephalon/SC_MAB21L1+
Rhombencephalon/SC_Prolif_NDNF+
Rhombencephalon_MAB21L1+
Rhombencephalon_Prolif
SC_Prolif_NDNF+_(perisyrinx)
SPON1+LGR5+_SC
SPON1+_Rhombencephalon
SpinalCord1_PAX3+_post
SpinalCord2a_SV2C+_ant
SpinalCord2b_SV2C+_ant_tip
Tel/Di_Migratory1_NKX2-1lo
Tel/Di_Migratory2_NKX2-1mid
Tel/Di_Migratory3_NKX2-1hi
Tel/Di_TPBG+RSPO2+
Telencephalon_EMX2+_Prolif
Telencephalon_NKX2-1+Prolif
Telencephalon_TRH+GNRH1+
other
b
MKI67
DCX
EMX2
FOXG1
PAX6
TRH
TPBG
RSPO2
NEUROG2
FEZF1
SORCS2
BARHL2
EN2
OTX2
FOXA1
PAX5
MAB21L1
TFAP2A
HOXB5
HOXB9
P2RX3
SPON1
PAX3
SV2C
GDF7

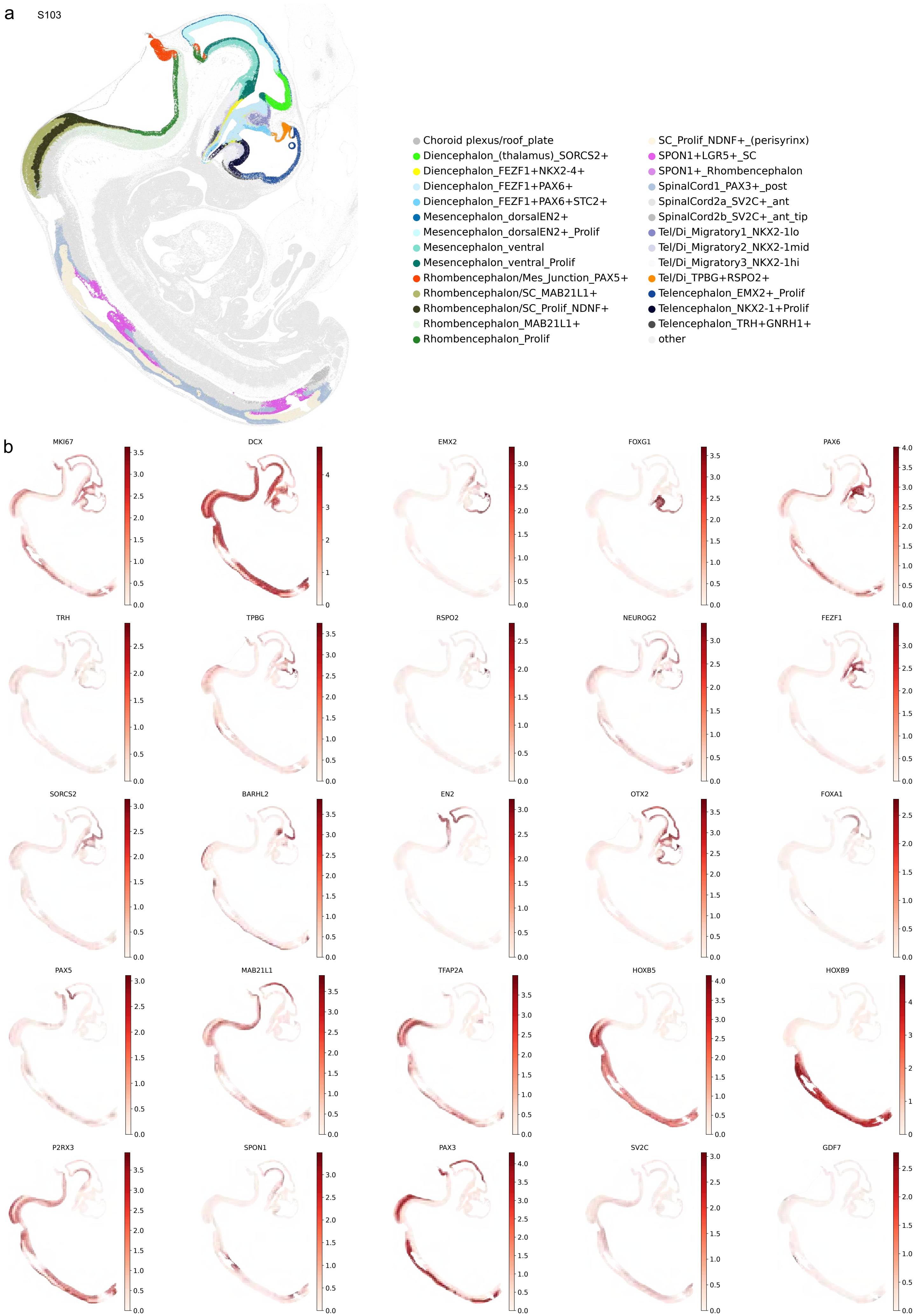

a
S103
Choroid plexus/roof_plate
Diencephalon_(thalamus)_SORCS2+
Diencephalon_FEZF1+NKX2-4+
Diencephalon_FEZF1+PAX6+
Diencephalon_FEZF1+PAX6+STC2+
Mesencephalon_dorsalEN2+
Mesencephalon_dorsalEN2+_Prolif
Mesencephalon_ventral
Mesencephalon_ventral_Prolif
Rhombencephalon/Mes_Junction_PAX5+
Rhombencephalon/SC_MAB21L1+
Rhombencephalon/SC_Prolif_NDNF+
Rhombencephalon_MAB21L1+
Rhombencephalon_Prolif
SC_Prolif_NDNF+_(perisyrinx)
SPON1+LGR5+_SC
SPON1+_Rhombencephalon
SpinalCord1_PAX3+_post
SpinalCord2a_SV2C+_ant
SpinalCord2b_SV2C+_ant_tip
Tel/Di_Migratory1_NKX2-1lo
Tel/Di_Migratory2_NKX2-1mid
Tel/Di_Migratory3_NKX2-1hi
Tel/Di_TPBG+RSPO2+
Telencephalon_EMX2+_Prolif
Telencephalon_NKX2-1+Prolif
Telencephalon_TRH+GNRH1+
other
b
MKI67
DCX
EMX2
FOXG1
PAX6
TRH
TPBG
RSPO2
NEUROG2
FEZF1
SORCS2
BARHL2
EN2
OTX2
FOXA1
PAX5
MAB21L1
TFAP2A
HOXB5
HOXB9
P2RX3
SPON1
PAX3
SV2C
GDF7
