## Appendix B for "An integrated single cell and spatial omics atlas of human prenatal development" (from another version of this preprint: version 1, 2026-04-01)

### Appendix B – Predominantly mesenchymal/stromal populations

**Appendix B Fig. 1 Gonad/adrenal/other.** a) UMAP embedding coloured by HDCA annotation (inlet: coloured by technology – scRNA-seq vs xenium 5k). b) UMAP embedding coloured by original author annotation (minimum 10 cells – number selected based on overall number of cells and number of overlapping labels in plot). c) Dot-plot of marker genes for cell populations in scRNA-seq and Xenium 5k. d) Whole embryo section profiled by Xenium 5k showing the location of each cell type.

**Appendix B Fig. 2 Mural.** a) UMAP embedding coloured by HDCA annotation (inlet: coloured by technology – scRNA-seq vs xenium 5k). b) UMAP embedding coloured by original author annotation (minimum 10 cells). c) Dot-plot of marker genes for cell populations in scRNA-seq and Xenium 5k. d) Whole embryo section profiled by Xenium 5k showing the location of each cell type.

**Appendix B Fig. 3 - Skin.** a) UMAP embedding coloured by HDCA annotation (inlet: coloured by technology – scRNA-seq vs xenium 5k). b) UMAP embedding coloured by original author annotation (minimum 100 cells). c) Dot-plot of marker genes for cell populations in scRNA-seq and Xenium 5k. d) Whole embryo section profiled by Xenium 5k showing the location of each cell type.

**Appendix B Fig. 4 - Cardiac.** a) UMAP embedding coloured by HDCA annotation (inlet: coloured by technology – scRNA-seq vs xenium 5k). b) UMAP embedding coloured by original author annotation (minimum 100 cells). c) Dot-plot of marker genes for cell populations in scRNA-seq and Xenium 5k. d) Whole embryo section profiled by Xenium 5k showing the location of each cell type.

**Appendix B Fig. 5 - Muscle.** a) UMAP embedding coloured by HDCA annotation (inlet: coloured by technology – scRNA-seq vs xenium 5k). b) UMAP embedding coloured by original author annotation (minimum 20 cells). c) Dot-plot of marker genes for cell populations in scRNA-seq and Xenium 5k. d) Whole embryo section profiled by Xenium 5k showing the location of each cell type.

**Appendix B Fig. 6 - Spine.** a) UMAP embedding coloured by HDCA annotation (inlet: coloured by technology – scRNA-seq vs xenium 5k). b) UMAP embedding coloured by original author annotation (minimum 100 cells). c) Dot-plot of marker genes for cell populations in scRNA-seq and Xenium 5k. d) Whole embryo section profiled by Xenium 5k showing the location of each cell type.

**Appendix B Fig. 7 - Skeletal.** a) UMAP embedding coloured by HDCA annotation (inlet: coloured by technology – scRNA-seq vs xenium 5k). b) UMAP embedding coloured by original author annotation (minimum 100 cells). c) Dot-plot of marker genes for cell populations in scRNA-seq and Xenium 5k. d) Whole embryo section profiled by Xenium 5k showing the location of each cell type.

**Appendix B Fig. 8 - Meninges.** a) UMAP embedding coloured by HDCA annotation (inlet: coloured by technology – scRNA-seq vs xenium 5k). b) UMAP embedding coloured by original author annotation (minimum 200 cells).

**Appendix B Fig. 9 - Eye.** a) UMAP embedding coloured by HDCA annotation. b) UMAP embedding coloured by original author annotation (minimum 10 cells). c) Dot-plot of marker genes for cell populations in scRNA-seq

(Fibroblast progenitors and mesothelial populations are included in Supplementary Figures)

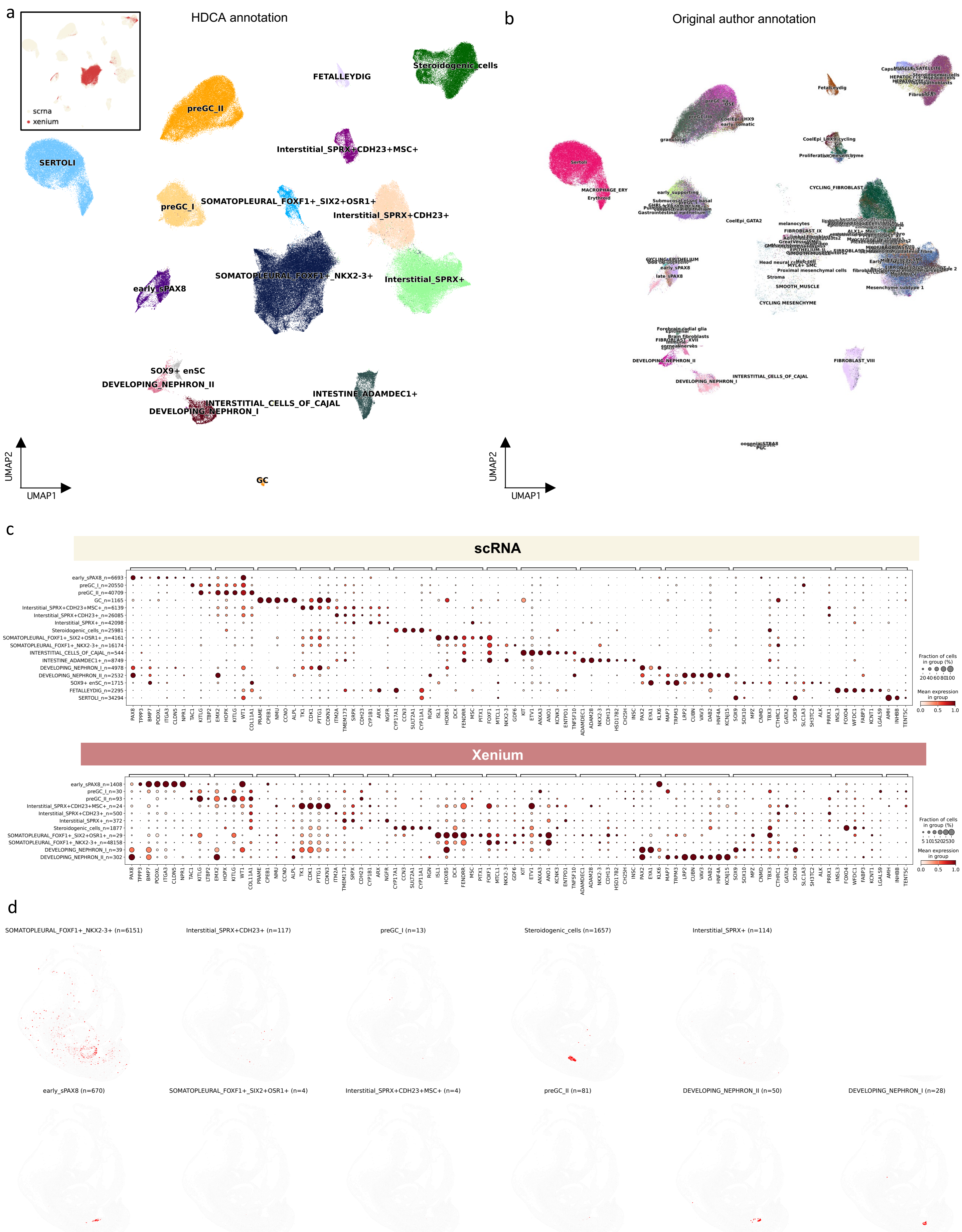
a
HDCA annotation
scrna
xenium
Steroidogenic_cells
FETALLEYDIG
preGC_II
Interstitial_SPRX+CDH23+MSC+
SERTOLI
preGC_I
SOMATOPLEURAL_FOXF1+_SIX2+OSR1+
Interstitial_SPRX+CDH23+
SOMATOPLEURAL_FOXF1+_NKX2-3+
Interstitial_SPRX+
early_sPAX8
SOX9+ enSC
DEVELOPING_NEPHRON_II
INTESTINE_ADAMDEC1+
INTERSTITIAL_CELLS_OF_CAJAL
DEVELOPING_NEPHRON_I
GC
UMAP2
UMAP1
b
Original author annotation
UMAP2
UMAP1
c
scRNA
early_sPAX8_n=6693
preGC_I_n=20550
preGC_II_n=40709
GC_n=1165
Interstitial_SPRX+CDH23+MSC+_n=6139
Interstitial_SPRX+CDH23+_n=26085
Interstitial_SPRX+_n=42098
Steroidogenic_cells_n=25981
SOMATOPLEURAL_FOXF1+_SIX2+OSR1+_n=4161
SOMATOPLEURAL_FOXF1+_NKX2-3+_n=16174
INTERSTITIAL_CELLS_OF_CAJAL_n=544
INTESTINE_ADAMDEC1+_n=8749
DEVELOPING_NEPHRON_I_n=4978
DEVELOPING_NEPHRON_II_n=2532
SOX9+ enSC_n=1715
FETALLEYDIG_n=2295
SERTOLI_n=34294
Fraction of cells in group (%)
20 40 60 80 100
Mean expression in group
0.0 0.5 1.0
Xenium
early_sPAX8_n=1408
preGC_I_n=30
preGC_II_n=93
Interstitial_SPRX+CDH23+MSC+_n=24
Interstitial_SPRX+CDH23+_n=500
Interstitial_SPRX+_n=372
Steroidogenic_cells_n=1877
SOMATOPLEURAL_FOXF1+_SIX2+OSR1+_n=29
SOMATOPLEURAL_FOXF1+_NKX2-3+_n=48158
DEVELOPING_NEPHRON_I_n=39
DEVELOPING_NEPHRON_II_n=302
Fraction of cells in group (%)
5 10 15 20 25 30
Mean expression in group
0.0 0.5 1.0
d
SOMATOPLEURAL_FOXF1+_NKX2-3+ (n=6151)
Interstitial_SPRX+CDH23+ (n=117)
preGC_I (n=13)
Steroidogenic_cells (n=1657)
Interstitial_SPRX+ (n=114)
early_sPAX8 (n=670)
SOMATOPLEURAL_FOXF1+_SIX2+OSR1+ (n=4)
Interstitial_SPRX+CDH23+MSC+ (n=4)
preGC_II (n=81)
DEVELOPING_NEPHRON_II (n=50)
DEVELOPING_NEPHRON_I (n=28)

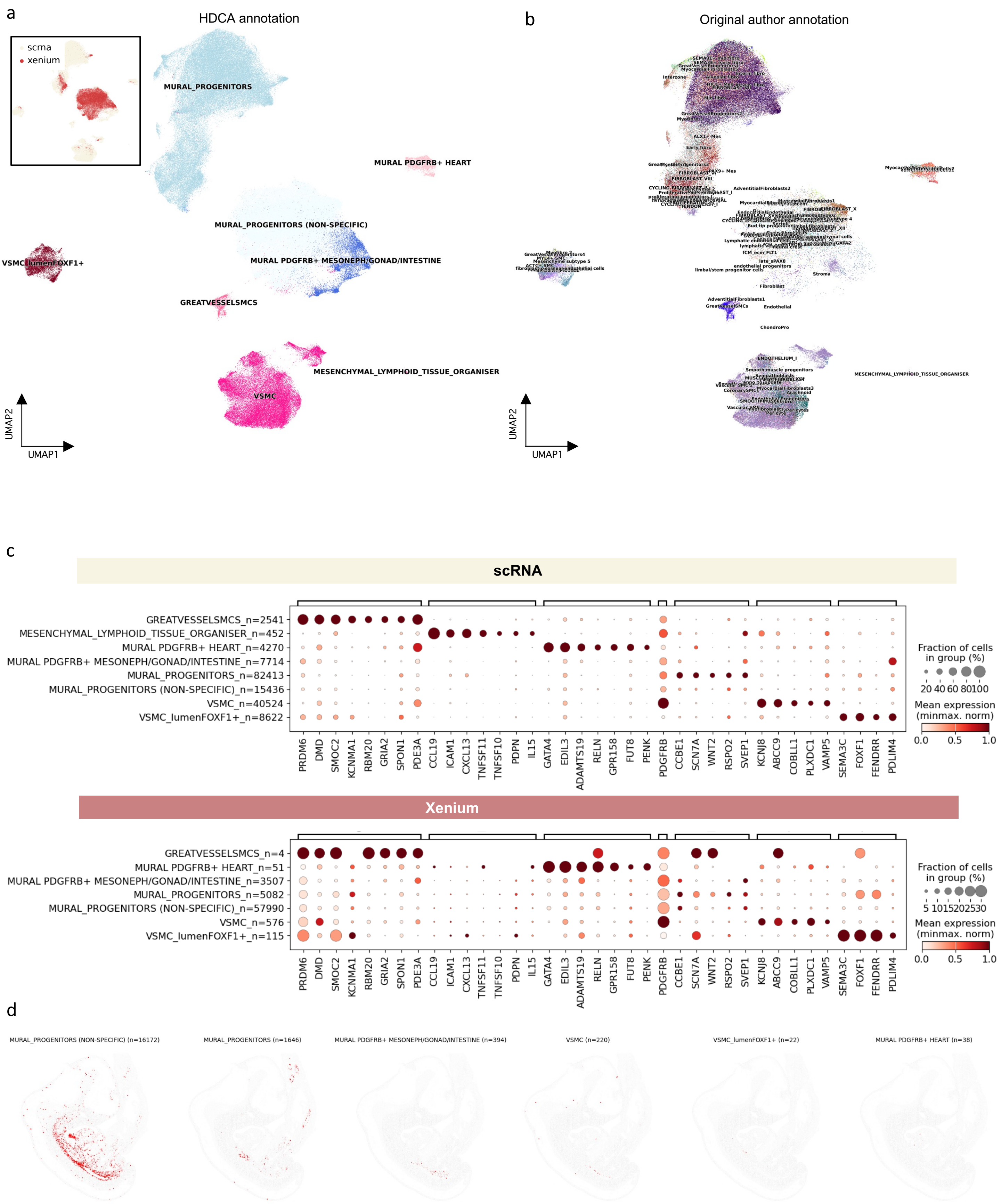
a
HDCA annotation
scrna
xenium
MURAL_PROGENITORS
MURAL PDGFRB+ HEART
MURAL_PROGENITORS (NON-SPECIFIC)
MURAL PDGFRB+ MESONEPH/GONAD/INTESTINE
VSMC_lumenFOXF1+
GREATVESSELSMCS
MESENCHYMAL_LYMPHOID_TISSUE_ORGANISER
VSMC
UMAP2
UMAP1
b
Original author annotation
UMAP2
UMAP1
c
scRNA
GREATVESSELSMCS_n=2541
MESENCHYMAL_LYMPHOID_TISSUE_ORGANISER_n=452
MURAL PDGFRB+ HEART_n=4270
MURAL PDGFRB+ MESONEPH/GONAD/INTESTINE_n=7714
MURAL_PROGENITORS_n=82413
MURAL_PROGENITORS (NON-SPECIFIC)_n=15436
VSMC_n=40524
VSMC_lumenFOXF1+_n=8622
PRDM6 DMD SMOC2 KCNMA1 RBM20 GRIA2 SPON1 PDE3A CCL19 ICAM1 CXCL13 TNFSF11 TNFSF10 PDPN IL15 GATA4 EDIL3 ADAMTS19 RELN GPR158 FUT8 PENK PDGFRB CCBE1 SCN7A WNT2 RSPO2 SVEP1 KCNJ8 ABCC9 COBLL1 PLXDC1 VAMP5 SEMA3C FOXF1 FENDRR PDLIM4
Fraction of cells in group (%)
20 40 60 80 100
Mean expression (minmax. norm)
0.0 0.5 1.0
Xenium
GREATVESSELSMCS_n=4
MURAL PDGFRB+ HEART_n=51
MURAL PDGFRB+ MESONEPH/GONAD/INTESTINE_n=3507
MURAL_PROGENITORS_n=5082
MURAL_PROGENITORS (NON-SPECIFIC)_n=57990
VSMC_n=576
VSMC_lumenFOXF1+_n=115
PRDM6 DMD SMOC2 KCNMA1 RBM20 GRIA2 SPON1 PDE3A CCL19 ICAM1 CXCL13 TNFSF11 TNFSF10 PDPN IL15 GATA4 EDIL3 ADAMTS19 RELN GPR158 FUT8 PENK PDGFRB CCBE1 SCN7A WNT2 RSPO2 SVEP1 KCNJ8 ABCC9 COBLL1 PLXDC1 VAMP5 SEMA3C FOXF1 FENDRR PDLIM4
Fraction of cells in group (%)
5 10 15 20 25 30
Mean expression (minmax. norm)
0.0 0.5 1.0
d
MURAL_PROGENITORS (NON-SPECIFIC) (n=16172)
MURAL_PROGENITORS (n=1646)
MURAL PDGFRB+ MESONEPH/GONAD/INTESTINE (n=394)
VSMC (n=220)
VSMC_lumenFOXF1+ (n=22)
MURAL PDGFRB+ HEART (n=38)

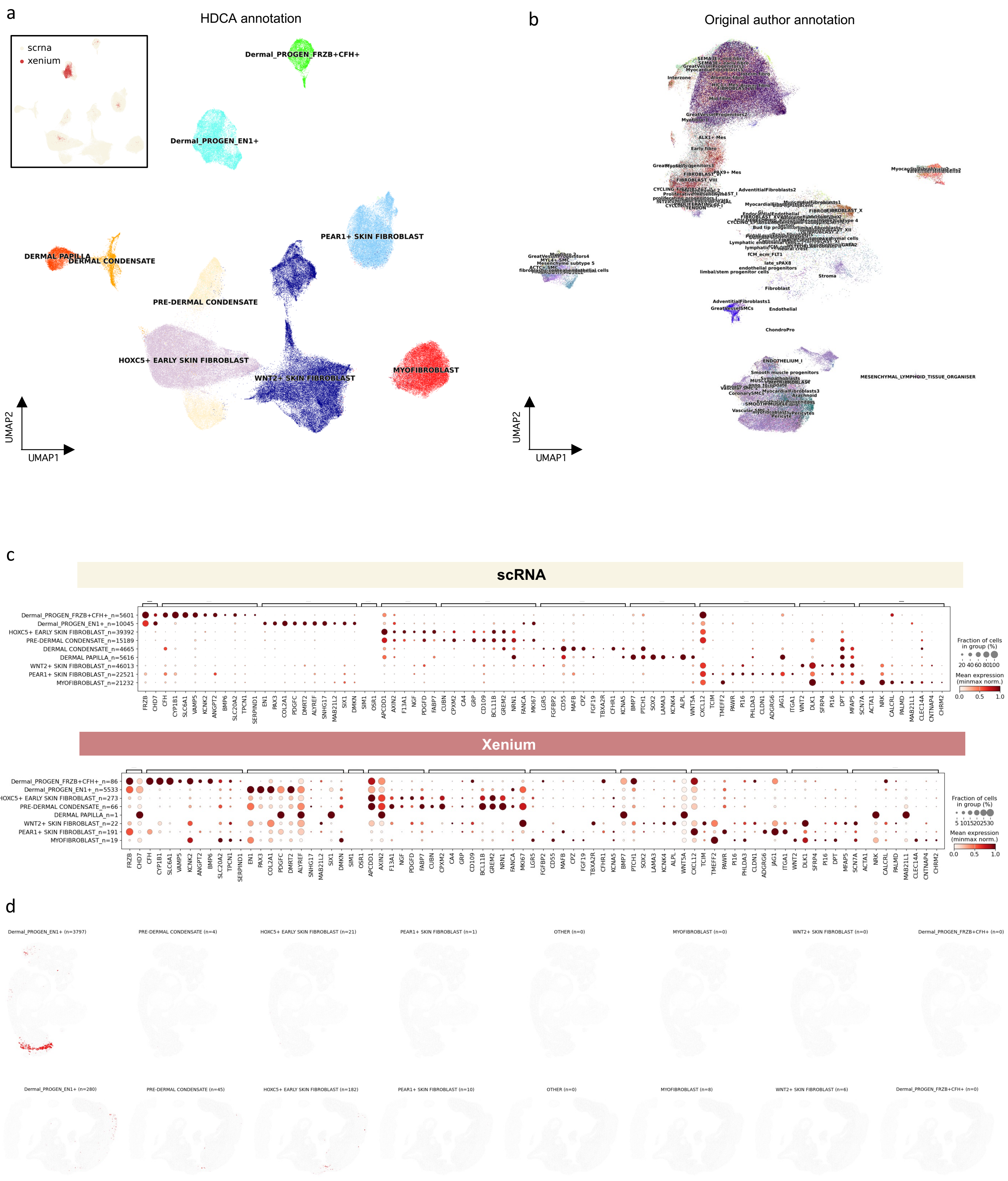
a
HDCA annotation
scrna
xenium
Dermal_PROGEN_FRZB+CFH+
Dermal_PROGEN_EN1+
PEAR1+ SKIN FIBROBLAST
DERMAL PAPILLA
DERMAL CONDENSATE
PRE-DERMAL CONDENSATE
HOXC5+ EARLY SKIN FIBROBLAST
WNT2+ SKIN FIBROBLAST
MYOFIBROBLAST
UMAP2
UMAP1
b
Original author annotation
UMAP2
UMAP1
c
scRNA
Dermal_PROGEN_FRZB+CFH+_n=5601
Dermal_PROGEN_EN1+_n=10045
HOXC5+ EARLY SKIN FIBROBLAST_n=39392
PRE-DERMAL CONDENSATE_n=15189
DERMAL CONDENSATE_n=4665
DERMAL PAPILLA_n=5616
WNT2+ SKIN FIBROBLAST_n=46013
PEAR1+ SKIN FIBROBLAST_n=22521
MYOFIBROBLAST_n=21232
Fraction of cells in group (%)
20 40 60 80 100
Mean expression (minmax norm.)
0.0 0.5 1.0
Xenium
Dermal_PROGEN_FRZB+CFH+_n=86
Dermal_PROGEN_EN1+_n=5533
HOXC5+ EARLY SKIN FIBROBLAST_n=273
PRE-DERMAL CONDENSATE_n=66
DERMAL PAPILLA_n=1
WNT2+ SKIN FIBROBLAST_n=22
PEAR1+ SKIN FIBROBLAST_n=191
MYOFIBROBLAST_n=19
Fraction of cells in group (%)
5 10 15 20 25 30
Mean expression (minmax norm.)
0.0 0.5 1.0
d
Dermal_PROGEN_EN1+ (n=3797)
PRE-DERMAL CONDENSATE (n=4)
HOXC5+ EARLY SKIN FIBROBLAST (n=21)
PEAR1+ SKIN FIBROBLAST (n=1)
OTHER (n=0)
MYOFIBROBLAST (n=0)
WNT2+ SKIN FIBROBLAST (n=0)
Dermal_PROGEN_FRZB+CFH+ (n=0)
Dermal_PROGEN_EN1+ (n=280)
PRE-DERMAL CONDENSATE (n=45)
HOXC5+ EARLY SKIN FIBROBLAST (n=182)
PEAR1+ SKIN FIBROBLAST (n=10)
OTHER (n=0)
MYOFIBROBLAST (n=8)
WNT2+ SKIN FIBROBLAST (n=6)
Dermal_PROGEN_FRZB+CFH+ (n=0)

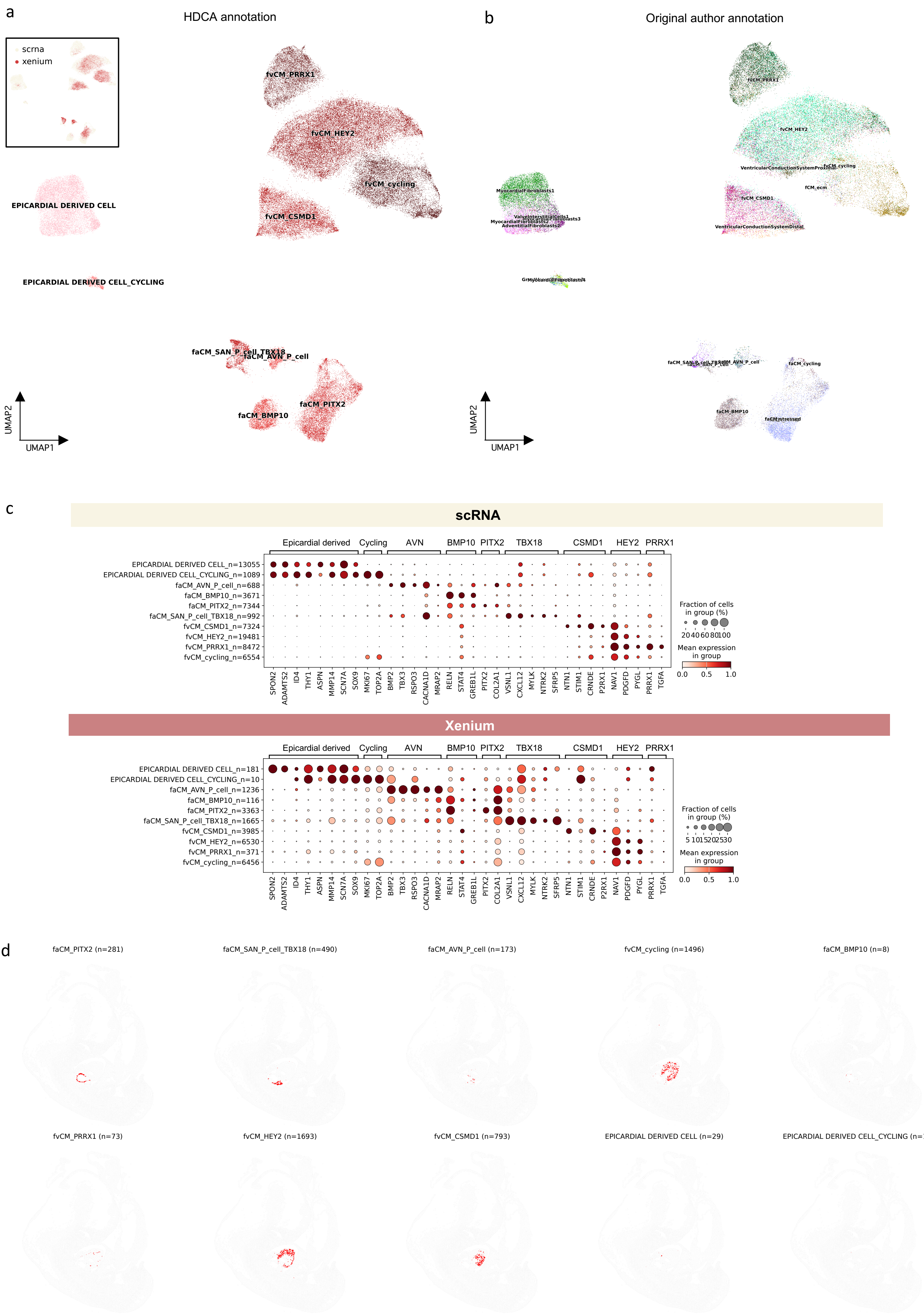

a
HDCA annotation
scrna
xenium
fvCM_PRRX1
fvCM_HEY2
fvCM_cycling
fvCM_CSMD1
EPICARDIAL DERIVED CELL
EPICARDIAL DERIVED CELL_CYCLING
faCM_SAN_P_cell_TBX18
faCM_AVN_P_cell
faCM_PITX2
faCM_BMP10
UMAP2
UMAP1
b
Original author annotation
fvCM_PRRX1
fvCM_HEY2
VentricularConductionSystemProximal
fvCM_cycling
fCM_ecm
fvCM_CSMD1
VentricularConductionSystemDistal
MyocardialFibroblasts1
AdventitialFibroblasts2
faCM_cycling
faCM_BMP10
UMAP2
UMAP1
c
scRNA
Epicardial derived
Cycling
AVN
BMP10
PITX2
TBX18
CSMD1
HEY2
PRRX1
EPICARDIAL DERIVED CELL_n=13055
EPICARDIAL DERIVED CELL_CYCLING_n=1089
faCM_AVN_P_cell_n=688
faCM_BMP10_n=3671
faCM_PITX2_n=7344
faCM_SAN_P_cell_TBX18_n=992
fvCM_CSMD1_n=7324
fvCM_HEY2_n=19481
fvCM_PRRX1_n=8472
fvCM_cycling_n=6554
SPON2
ADAMTS2
ID4
THY1
ASPN
MMP14
SCN7A
SOX9
MKI67
TOP2A
BMP2
TBX3
RSPO3
CACNA1D
MRAP2
RELN
STAT4
GREB1L
PITX2
COL2A1
VSNL1
CXCL12
MYLK
NTRK2
SFRP5
NTN1
STIM1
CRNDE
P2RX1
NAV1
PDGFD
PYGL
PRRX1
TGFA
Fraction of cells in group (%)
20 40 60 80 100
Mean expression in group
0.0
0.5
1.0
Xenium
Epicardial derived
Cycling
AVN
BMP10
PITX2
TBX18
CSMD1
HEY2
PRRX1
EPICARDIAL DERIVED CELL_n=181
EPICARDIAL DERIVED CELL_CYCLING_n=10
faCM_AVN_P_cell_n=1236
faCM_BMP10_n=116
faCM_PITX2_n=3363
faCM_SAN_P_cell_TBX18_n=1665
fvCM_CSMD1_n=3985
fvCM_HEY2_n=6530
fvCM_PRRX1_n=371
fvCM_cycling_n=6456
SPON2
ADAMTS2
ID4
THY1
ASPN
MMP14
SCN7A
SOX9
MKI67
TOP2A
BMP2
TBX3
RSPO3
CACNA1D
MRAP2
RELN
STAT4
GREB1L
PITX2
COL2A1
VSNL1
CXCL12
MYLK
NTRK2
SFRP5
NTN1
STIM1
CRNDE
P2RX1
NAV1
PDGFD
PYGL
PRRX1
TGFA
Fraction of cells in group (%)
5 10 15 20 25 30
Mean expression in group
0.0
0.5
1.0
d
faCM_PITX2 (n=281)
faCM_SAN_P_cell_TBX18 (n=490)
faCM_AVN_P_cell (n=173)
fvCM_cycling (n=1496)
faCM_BMP10 (n=8)
fvCM_PRRX1 (n=73)
fvCM_HEY2 (n=1693)
fvCM_CSMD1 (n=793)
EPICARDIAL DERIVED CELL (n=29)

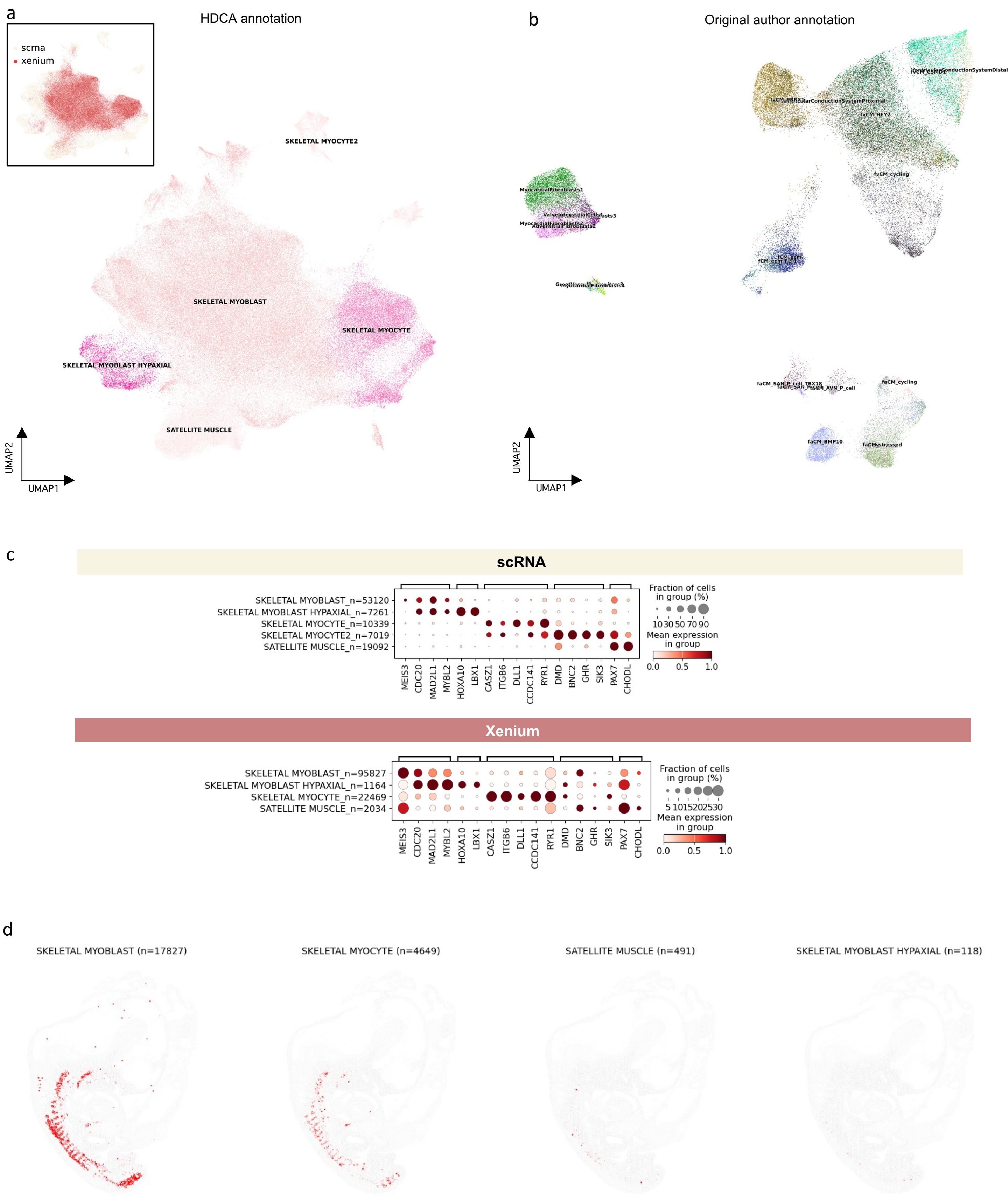
a
HDCA annotation
scrna
xenium
SKELETAL MYOCYTE2
SKELETAL MYOBLAST
SKELETAL MYOCYTE
SKELETAL MYOBLAST HYPAXIAL
SATELLITE MUSCLE
UMAP2
UMAP1
b
Original author annotation
fvCM_PRRX1
fvCM_HEY2
fvCM_cycling
MyocardialFibroblasts1
faCM_cycling
faCM_BMP10
UMAP2
UMAP1
c
scRNA
SKELETAL MYOBLAST_n=53120
SKELETAL MYOBLAST HYPAXIAL_n=7261
SKELETAL MYOCYTE_n=10339
SKELETAL MYOCYTE2_n=7019
SATELLITE MUSCLE_n=19092
MEIS3
CDC20
MAD2L1
MYBL2
HOXA10
LBX1
CASZ1
ITGB6
DLL1
CCDC141
RYR1
DMD
BNC2
GHR
SIK3
PAX7
CHODL
Fraction of cells in group (%)
10 30 50 70 90
Mean expression in group
0.0 0.5 1.0
Xenium
SKELETAL MYOBLAST_n=95827
SKELETAL MYOBLAST HYPAXIAL_n=1164
SKELETAL MYOCYTE_n=22469
SATELLITE MUSCLE_n=2034
MEIS3
CDC20
MAD2L1
MYBL2
HOXA10
LBX1
CASZ1
ITGB6
DLL1
CCDC141
RYR1
DMD
BNC2
GHR
SIK3
PAX7
CHODL
Fraction of cells in group (%)
5 10 15 20 25 30
Mean expression in group
0.0 0.5 1.0
d
SKELETAL MYOBLAST (n=17827)
SKELETAL MYOCYTE (n=4649)
SATELLITE MUSCLE (n=491)
SKELETAL MYOBLAST HYPAXIAL (n=118)

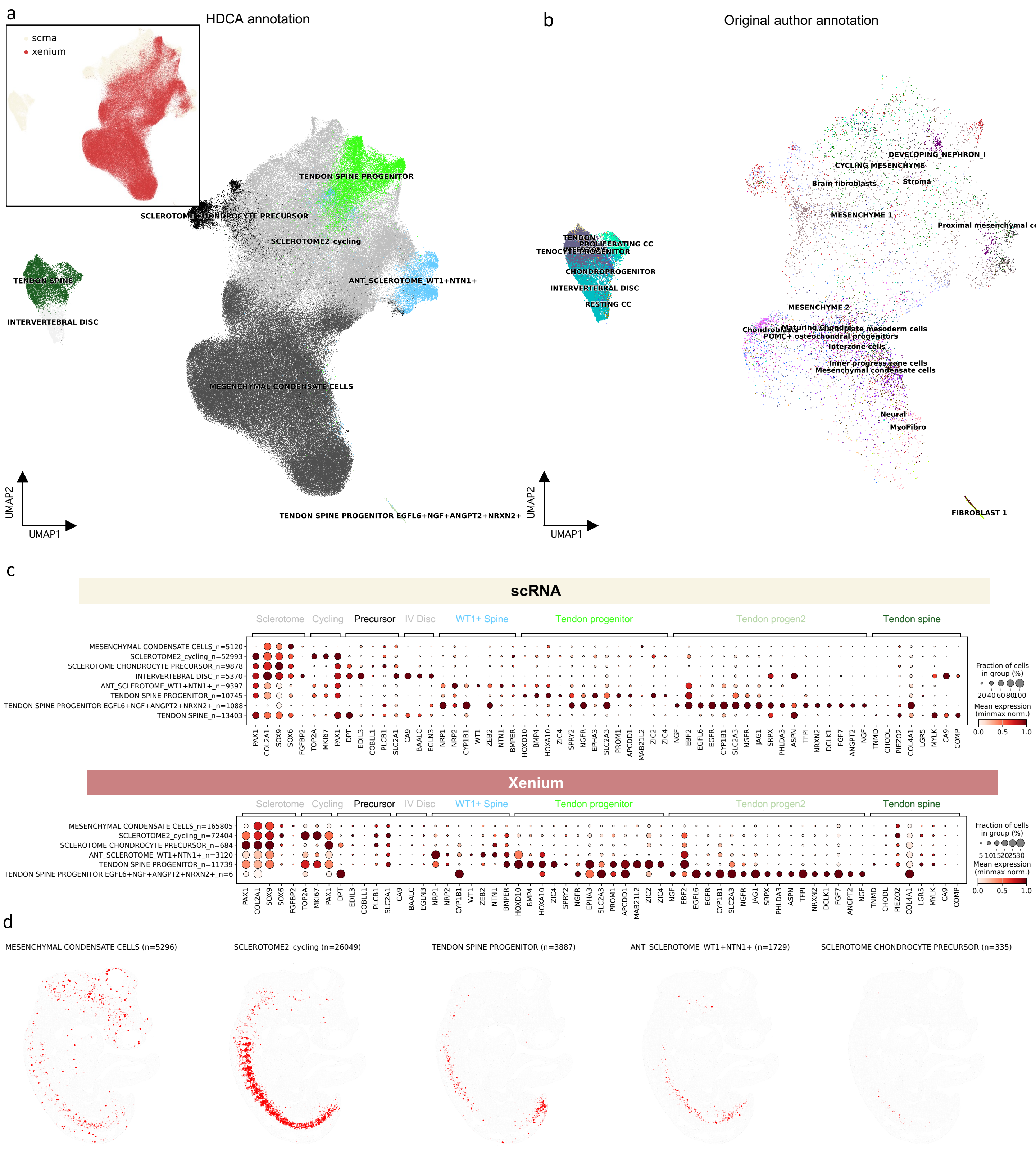
a
HDCA annotation
scrna
xenium
TENDON SPINE PROGENITOR
SCLEROTOME CHONDROCYTE PRECURSOR
SCLEROTOME2_cycling
ANT_SCLEROTOME_WT1+NTN1+
TENDON SPINE
INTERVERTEBRAL DISC
MESENCHYMAL CONDENSATE CELLS
TENDON SPINE PROGENITOR EGFL6+NGF+ANGPT2+NRXN2+
UMAP2
UMAP1
b
Original author annotation
DEVELOPING_NEPHRON_I
CYCLING MESENCHYME
Brain fibroblasts
Stroma
MESENCHYME 1
Proximal mesenchymal ce
TENDON
PROLIFERATING CC
TENOCYTE PROGENITOR
CHONDROPROGENITOR
INTERVERTEBRAL DISC
RESTING CC
MESENCHYME 2
Chondroblasts
Maturing Chondrocytes
Paraxial mesoderm cells
POMC+ osteochondral progenitors
Interzone cells
Inner progress zone cells
Mesenchymal condensate cells
Neural
MyoFibro
FIBROBLAST 1
UMAP2
UMAP1
c
scRNA
Sclerotome
Cycling
Precursor
IV Disc
WT1+ Spine
Tendon progenitor
Tendon progen2
Tendon spine
MESENCHYMAL CONDENSATE CELLS_n=5120
SCLEROTOME2_cycling_n=52993
SCLEROTOME CHONDROCYTE PRECURSOR_n=9878
INTERVERTEBRAL DISC_n=5370
ANT_SCLEROTOME_WT1+NTN1+_n=9397
TENDON SPINE PROGENITOR_n=10745
TENDON SPINE PROGENITOR EGFL6+NGF+ANGPT2+NRXN2+_n=1088
TENDON SPINE_n=13403
PAX1 COL2A1 SOX9 SOX6 FGFBP2 TOP2A MKI67 PAX1 DPT EDIL3 COBLL1 PLCB1 SLC2A1 CA9 BAALC EGLN3 NRP1 NRP2 CYP1B1 WT1 ZEB2 NTN1 BMPER HOXD10 BMP4 HOXA10 ZIC4 SPRY2 NGFR EPHA3 SLC2A3 PROM1 APCDD1 MAB21L2 ZIC2 ZIC4 NGF EBF2 EGFL6 EGFR CYP1B1 SLC2A3 NGFR JAG1 SRPX PHLDA3 ASPN TFPI NRXN2 DCLK1 FGF7 ANGPT2 NGF TNMD CHODL PIEZO2 COL4A1 LGR5 MYLK CA9 COMP
Fraction of cells in group (%)
20 40 60 80 100
Mean expression (minmax norm.)
0.0 0.5 1.0
Xenium
Sclerotome
Cycling
Precursor
IV Disc
WT1+ Spine
Tendon progenitor
Tendon progen2
Tendon spine
MESENCHYMAL CONDENSATE CELLS_n=165805
SCLEROTOME2_cycling_n=72404
SCLEROTOME CHONDROCYTE PRECURSOR_n=684
ANT_SCLEROTOME_WT1+NTN1+_n=3120
TENDON SPINE PROGENITOR_n=11739
TENDON SPINE PROGENITOR EGFL6+NGF+ANGPT2+NRXN2+_n=6
PAX1 COL2A1 SOX9 SOX6 FGFBP2 TOP2A MKI67 PAX1 DPT EDIL3 COBLL1 PLCB1 SLC2A1 CA9 BAALC EGLN3 NRP1 NRP2 CYP1B1 WT1 ZEB2 NTN1 BMPER HOXD10 BMP4 HOXA10 ZIC4 SPRY2 NGFR EPHA3 SLC2A3 PROM1 APCDD1 MAB21L2 ZIC2 ZIC4 NGF EBF2 EGFL6 EGFR CYP1B1 SLC2A3 NGFR JAG1 SRPX PHLDA3 ASPN TFPI NRXN2 DCLK1 FGF7 ANGPT2 NGF TNMD CHODL PIEZO2 COL4A1 LGR5 MYLK CA9 COMP
Fraction of cells in group (%)
5 10 15 20 25 30
Mean expression (minmax norm.)
0.0 0.5 1.0
d
MESENCHYMAL CONDENSATE CELLS (n=5296)
SCLEROTOME2_cycling (n=26049)
TENDON SPINE PROGENITOR (n=3887)
ANT_SCLEROTOME_WT1+NTN1+ (n=1729)
SCLEROTOME CHONDROCYTE PRECURSOR (n=335)

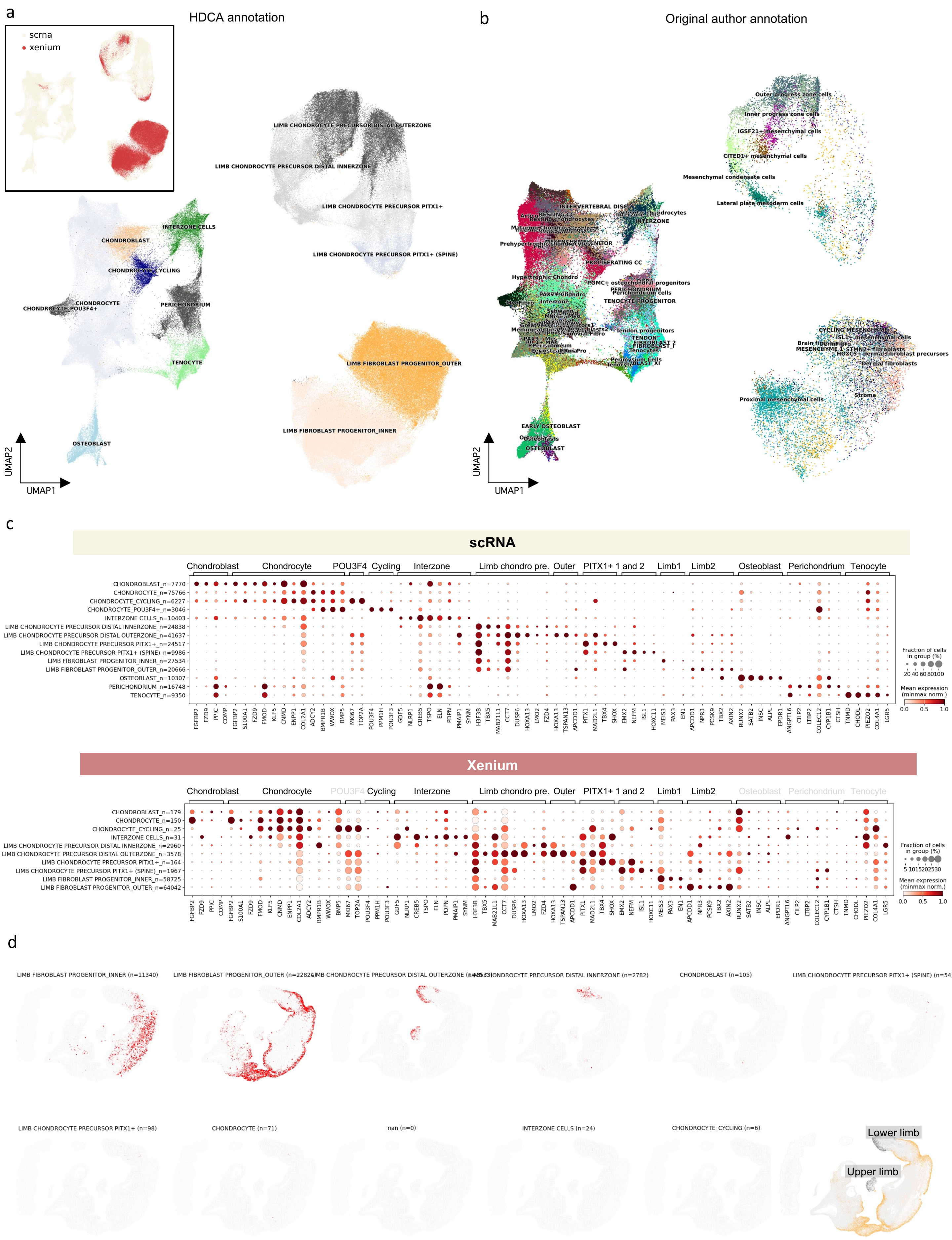
a
HDCA annotation
scrna
xenium
LIMB CHONDROCYTE PRECURSOR DISTAL OUTERZONE
LIMB CHONDROCYTE PRECURSOR DISTAL INNERZONE
LIMB CHONDROCYTE PRECURSOR PITX1+
LIMB CHONDROCYTE PRECURSOR PITX1+ (SPINE)
INTERZONE CELLS
CHONDROBLAST
CHONDROCYTE_CYCLING
CHONDROCYTE
CHONDROCYTE_POU3F4+
PERICHONDRIUM
TENOCYTE
OSTEOBLAST
LIMB FIBROBLAST PROGENITOR_OUTER
LIMB FIBROBLAST PROGENITOR_INNER
UMAP2
UMAP1
b
Original author annotation
Outer progress zone cells
Inner progress zone cells
IGSF21+ mesenchymal cells
CITED1+ mesenchymal cells
Mesenchymal condensate cells
Lateral plate mesoderm cells
INTERVERTEBRAL DISC
INTERZONE
Resting chondrocytes
Prehypertrophic chondrocytes
PROLIFERATING CC
Hypertrophic Chondro
POMC+ osteochondral progenitors
Perichondrium cells
TENOCYTE PROGENITOR
Interzone
Tendon progenitors
Tenocytes
Perimysium Cells
CYCLING MESENCHYME
ISL1+ mesenchymal cells
Brain fibroblast
HOXC5+ dermal fibroblast precursors
Dermal fibroblasts
Stroma
Proximal mesenchymal cells
EARLY OSTEOBLAST
Osteoblasts
OSTEOBLAST
UMAP2
UMAP1
c
scRNA
Chondroblast
Chondrocyte
POU3F4
Cycling
Interzone
Limb chondro pre.
Outer
PITX1+ 1 and 2
Limb1
Limb2
Osteoblast
Perichondrium
Tenocyte
CHONDROBLAST_n=7770
CHONDROCYTE_n=75766
CHONDROCYTE_CYCLING_n=6227
CHONDROCYTE_POU3F4+_n=3046
INTERZONE CELLS_n=10403
LIMB CHONDROCYTE PRECURSOR DISTAL INNERZONE_n=24838
LIMB CHONDROCYTE PRECURSOR DISTAL OUTERZONE_n=41637
LIMB CHONDROCYTE PRECURSOR PITX1+_n=24517
LIMB CHONDROCYTE PRECURSOR PITX1+ (SPINE)_n=9986
LIMB FIBROBLAST PROGENITOR_INNER_n=27534
LIMB FIBROBLAST PROGENITOR_OUTER_n=20666
OSTEOBLAST_n=10307
PERICHONDRIUM_n=16748
TENOCYTE_n=9350
Fraction of cells in group (%)
20 40 60 80 100
Mean expression (minmax norm.)
0.0 0.5 1.0
FGFBP2 FZD9 PPIC COMP FGFBP2 S100A1 FZD9 FMOD KLF5 CNMD ENPP1 COL2A1 ADCY2 BMPR1B WWOX BMP5 MKI67 TOP2A POU3F4 PPM1H POU3F3 GDF5 NLRP1 CREB5 TSPO ELN PDPN PMAIP1 SYNM H3F3B TBX5 MAB21L1 CCT7 DUSP6 HOXA13 LMO2 FZD4 HOXA13 TSPAN13 APCDD1 PITX1 MAD2L1 TBX4 SHOX EMX2 NEFM ISL1 HOXC11 MEIS3 PAX3 EN1 APCDD1 NPR3 PCSK9 TBX2 AXIN2 RUNX2 SATB2 INSC ALPL EPOR1 ANGPTL6 CILP2 LTBP2 COLEC12 CYP1B1 CTSH TNMD CHODL PIEZO2 COL4A1 LGR5
Xenium
CHONDROBLAST_n=179
CHONDROCYTE_n=150
CHONDROCYTE_CYCLING_n=25
INTERZONE CELLS_n=31
LIMB CHONDROCYTE PRECURSOR DISTAL INNERZONE_n=2960
LIMB CHONDROCYTE PRECURSOR DISTAL OUTERZONE_n=3578
LIMB CHONDROCYTE PRECURSOR PITX1+_n=164
LIMB CHONDROCYTE PRECURSOR PITX1+ (SPINE)_n=1967
LIMB FIBROBLAST PROGENITOR_INNER_n=58725
LIMB FIBROBLAST PROGENITOR_OUTER_n=64042
Fraction of cells in group (%)
5 10 15 20 25 30
Mean expression (minmax norm.)
0.0 0.5 1.0
d
LIMB FIBROBLAST PROGENITOR_INNER (n=11340)
LIMB FIBROBLAST PROGENITOR_OUTER (n=22824)
LIMB CHONDROCYTE PRECURSOR DISTAL OUTERZONE
LIMB CHONDROCYTE PRECURSOR DISTAL INNERZONE (n=2782)
CHONDROBLAST (n=105)
LIMB CHONDROCYTE PRECURSOR PITX1+ (SPINE)
LIMB CHONDROCYTE PRECURSOR PITX1+ (n=98)
CHONDROCYTE (n=71)
nan (n=0)
INTERZONE CELLS (n=24)
CHONDROCYTE_CYCLING (n=6)
Lower limb
Upper limb

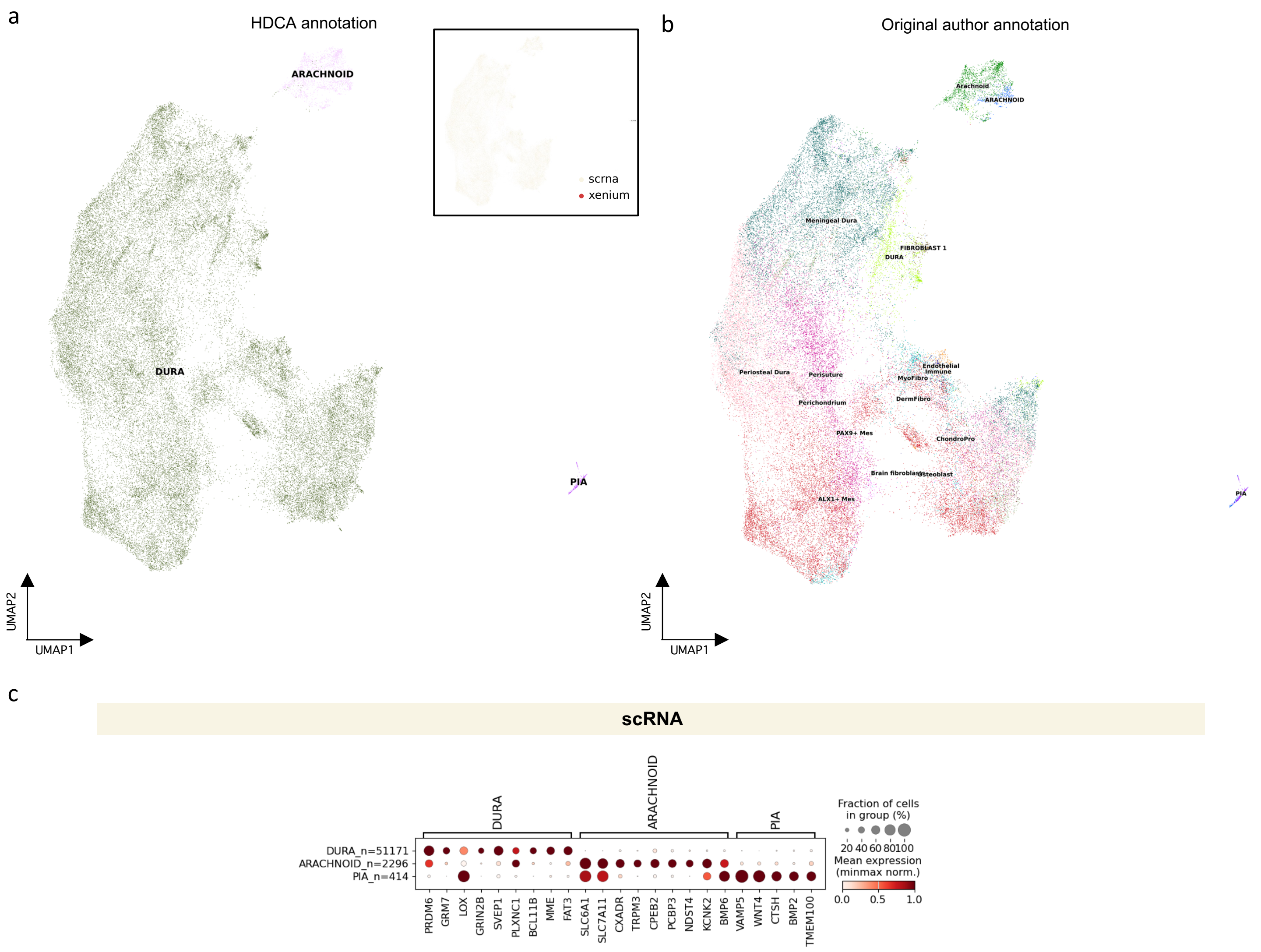
a
HDCA annotation
ARACHNOID
DURA
PIA
scrna
xenium
UMAP2
UMAP1
b
Original author annotation
Arachnoid
ARACHNOID
Meningeal Dura
FIBROBLAST 1
DURA
Periosteal Dura
Perisuture
Endothelial
Immune
MyoFibro
DermFibro
Perichondrium
PAX9+ Mes
ChondroPro
Brain fibroblast
Osteoblast
ALX1+ Mes
PIA
UMAP2
UMAP1
c
scRNA
DURA
ARACHNOID
PIA
DURA_n=51171
ARACHNOID_n=2296
PIA_n=414
PRDM6
GRM7
LOX
GRIN2B
SVEP1
PLXNC1
BCL11B
MME
FAT3
SLC6A1
SLC7A11
CXADR
TRPM3
CPEB2
PCBP3
NDST4
KCNK2
BMP6
VAMP5
WNT4
CTSH
BMP2
TMEM100
Fraction of cells
in group (%)
20 40 60 80 100
Mean expression
(minmax norm.)
0.0 0.5 1.0

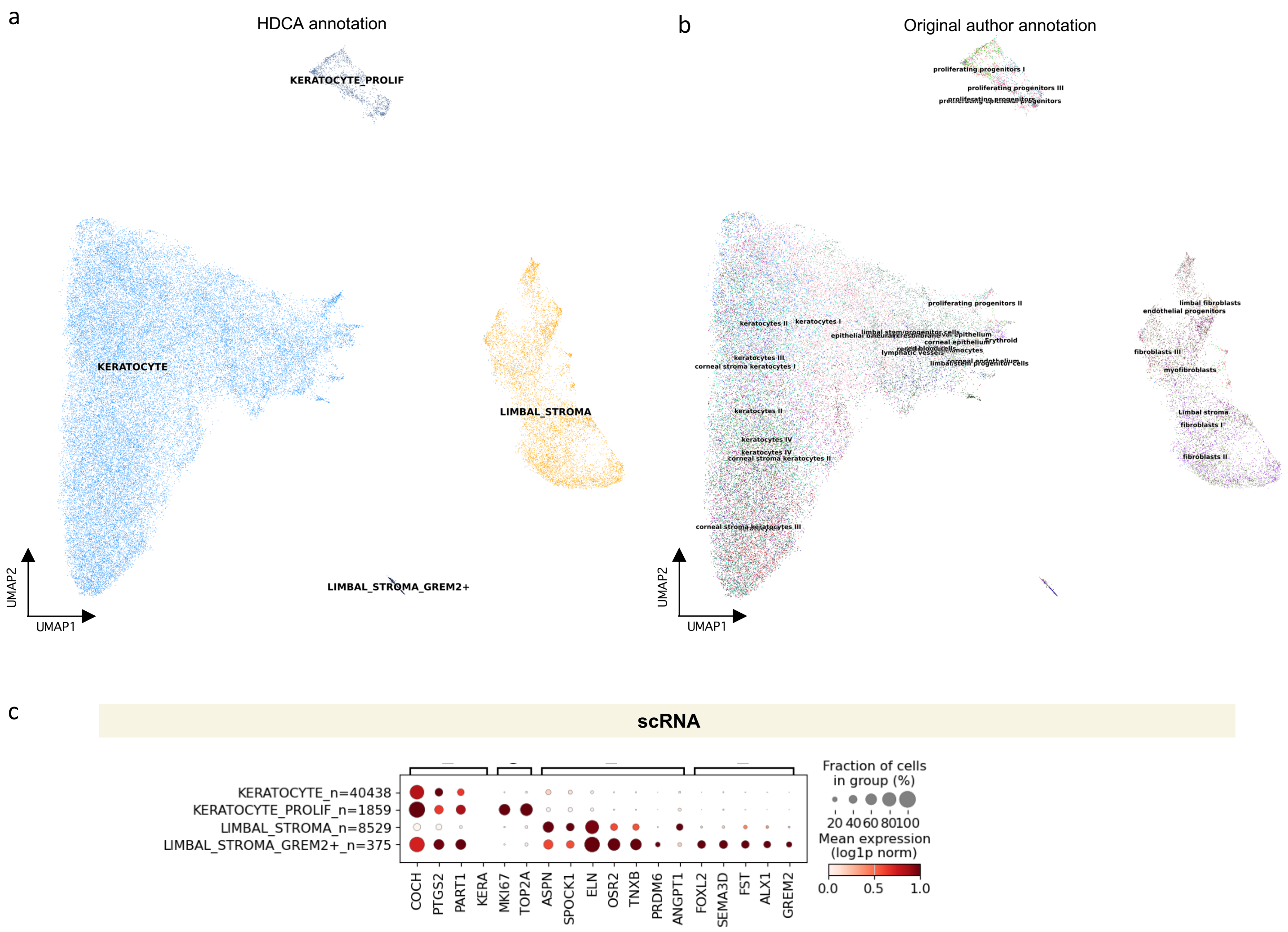
a
HDCA annotation
KERATOCYTE_PROLIF
KERATOCYTE
LIMBAL_STROMA
LIMBAL_STROMA_GREM2+
UMAP2
UMAP1
b
Original author annotation
proliferating progenitors I
proliferating progenitors III
proliferating progenitors II
keratocytes II
keratocytes I
keratocytes III
corneal stroma keratocytes I
keratocytes IV
corneal stroma keratocytes II
corneal stroma keratocytes III
corneal epithelium
erythroid
lymphatic vessels
limbal fibroblasts
endothelial progenitors
fibroblasts III
myofibroblasts
Limbal stroma
fibroblasts I
fibroblasts II
UMAP2
UMAP1
c
scRNA
KERATOCYTE_n=40438
KERATOCYTE_PROLIF_n=1859
LIMBAL_STROMA_n=8529
LIMBAL_STROMA_GREM2+_n=375
COCH
PTGS2
PART1
KERA
MKI67
TOP2A
ASPN
SPOCK1
ELN
OSR2
TNXB
PRDM6
ANGPT1
FOXL2
SEMA3D
FST
ALX1
GREM2
Fraction of cells in group (%)
20 40 60 80 100
Mean expression (log1p norm)
0.0 0.5 1.0
