## Supplementary Figures 1-6 for "An integrated single cell and spatial omics atlas of human prenatal development" (from another version of this preprint: version 1, 2026-04-01)

### Figure S1: Integrated single cell and spatial omics atlas of human prenatal development

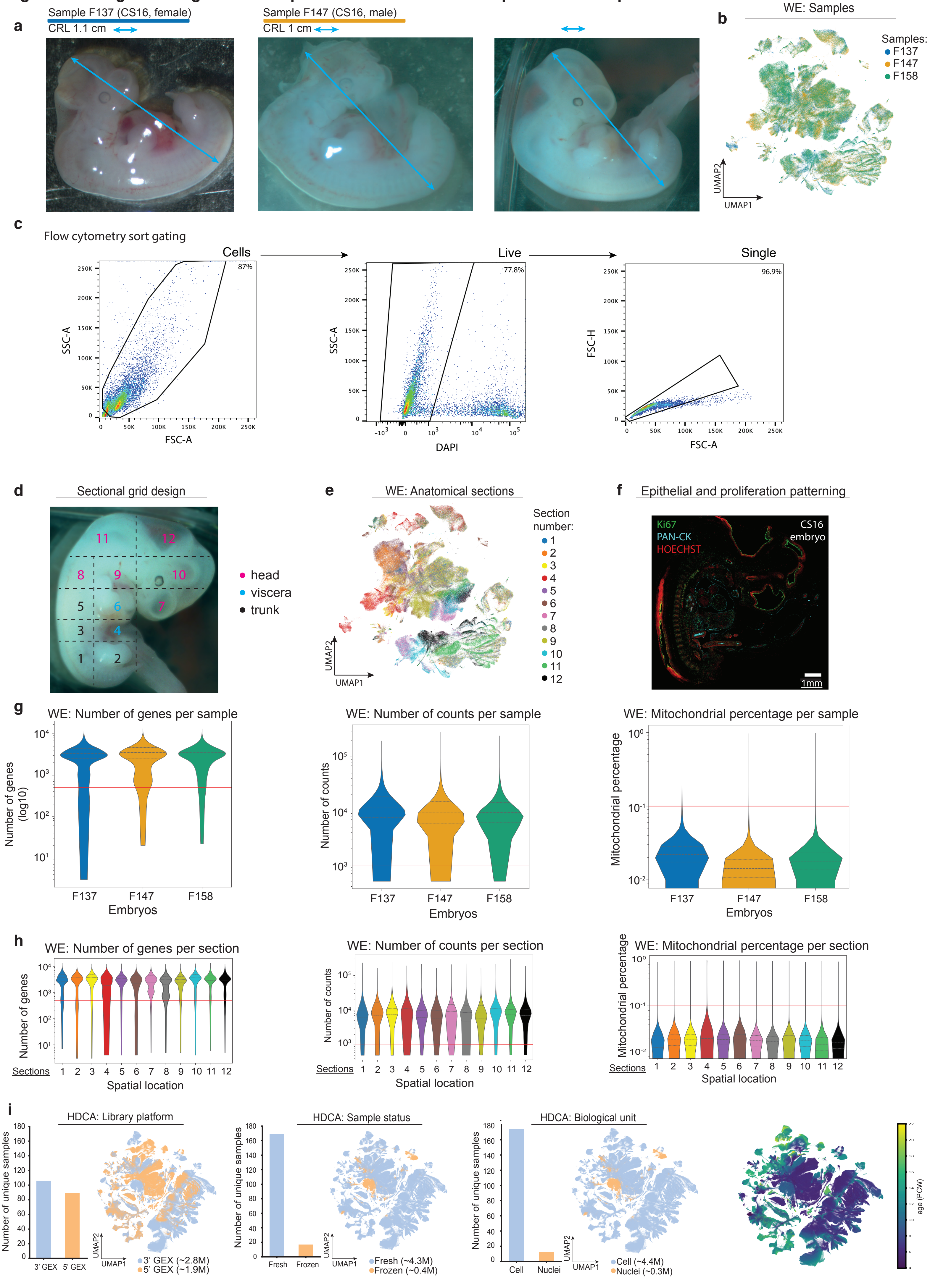

**Figure S1: Integrated single cell and spatial omics atlas of human prenatal development**

**a)** Images of intact embryos F137 (CS16, female), F147 (CS16, male) and F158 (CS17, female) with an attached yolk sac for F158. Blue arrows indicate the crown-rump length (CRL) measurements, which were 1.1cm, 1cm and 1.1cm respectively. **b)** UMAP visualisation of whole human embryo (WE) scRNA-seq data (n=3; k=1,029,326) where dot colour indicates sample. **c)** Gating strategy used to isolate cells for the scRNAseq experiments. This is a representative strategy for all samples and sections, taken from sample F158 section 3**. d)** F137 image in sagittal plane and overlaid with 12 sectional grid design depicting the representative tissue dissections undertaken prior to scRNA-seq data generation. Pink = Head sections, Blue = Viscera sections, Black = Trunk sections. **e)** UMAP visualisation of WE scRNA-seq data where dot colour indicates anatomical section number outlined in part C. **f)** RareCyte immunofluorescence imaging of a CS16 embryo with Hoescht staining (red) and antibodies for Ki67 (green) and Pan-Cytokeratin (blue). Scale bar=1mm. **g)** Violin plots for whole embryo scRNAseq data, stratified by sample (top) and section (bottom) for quality control measures on a log10 scale. Left: number of genes, middle: number of counts, right: mitochondrial percentage. Red lines across plots indicate threshold applied being ≥ 500 genes, ≥ 1,000 counts and ≤ 10% respectively. **h)** The same QC metrics as in (g), but per anatomical pseudospatial section, as in panel (d). **i)** Barplots and UMAP visualisations depicting the distribution of Library platform (3’ or 5’ chemistry), Sample status (Fresh or Frozen), Biological unit (Cell or Nuclei), or Age, respectively within the HDCA data. Numbers for each category rounded to nearest decimal point in million, actual numbers in **Supplementary Table 7**.

### Figure S2 : Integrated single cell and spatial omics atlas of human prenatal development

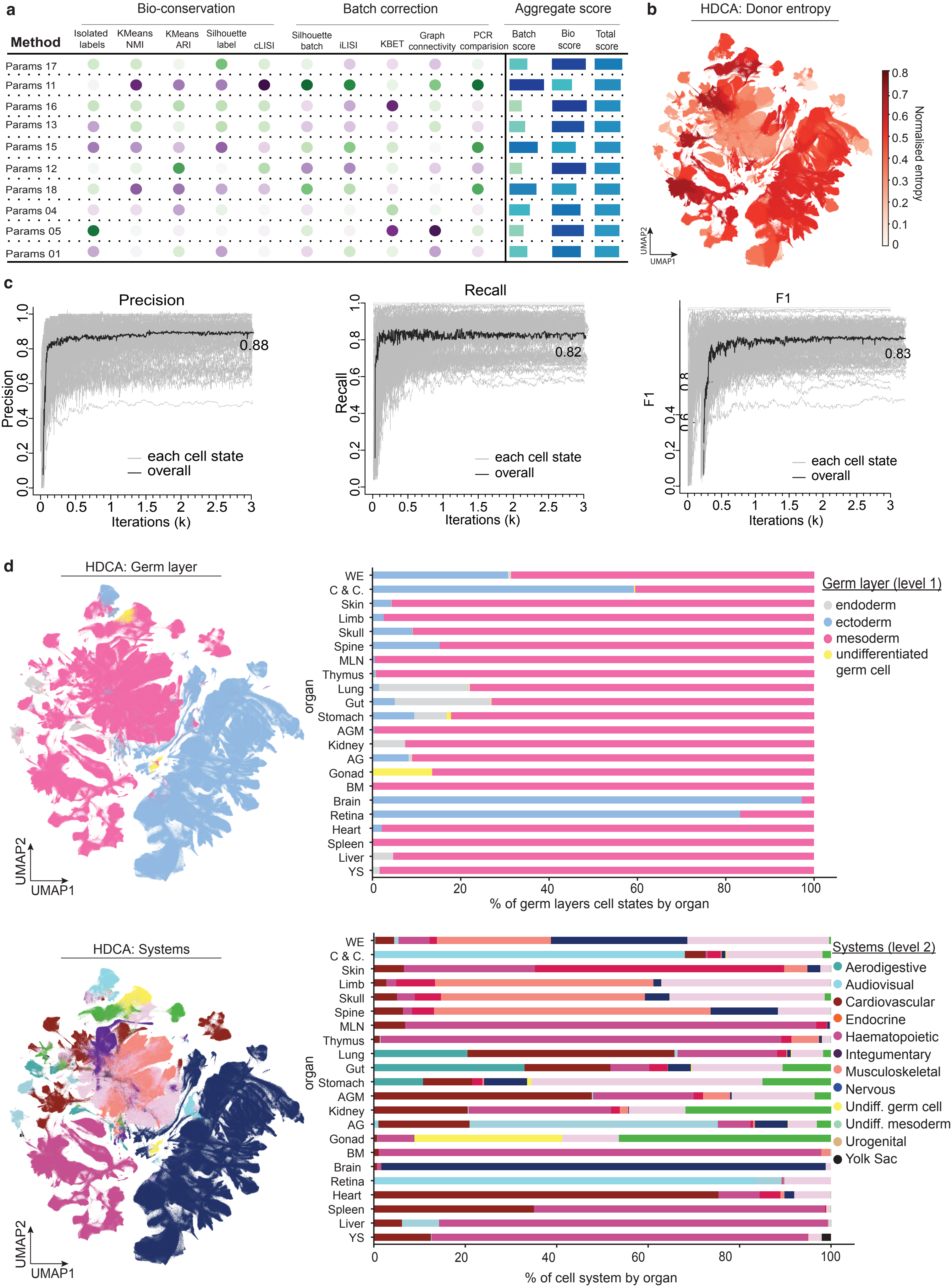

**Figure S2: Integrated single cell and spatial omics atlas of human prenatal development**

**a)** Summary table of HDCA integration benchmarking results using scib-metrics showing the top 10 best performing models. Batch correction score and biological conservation score each show the mean across metrics of that type, with metric scores scaled between range 0 and 1. Remaining models and quantitative score values stored in **Supplementary Table 8**. **b)** UMAP visualisation of sample mixture (entropy) within the HDCA integrated landscape across leiden clusters (scVI latents, KNN=30, leiden resolution=3). Entropy scores per cluster normalized between range 0 and 1 with a vmax=0.8 applied for visualisation. **c)** Line charts showing the performance of a global CellTypist model trained on public single cell/nuclei data outlined in **Fig. S1** for precision (left), recall (middle) and F1 scores (right) for each iteration of training. Black lines represent the median scores averaged across individual scores of all predicted cell types. **d)** Top: UMAP visualization of HDCA data coloured by Germ layer (left, level 1) and barplot distribution of Germ layer across individual organs and whole human embryo datasets within the HDCA data (right). Bottom: UMAP visualization of HDCA data coloured by system (left, level 2) and barplot distribution of systems across individual organs and whole human embryo datasets within the HDCA data (right).

### Figure S3: Benchmarking atlas integration with trajectory fidelity

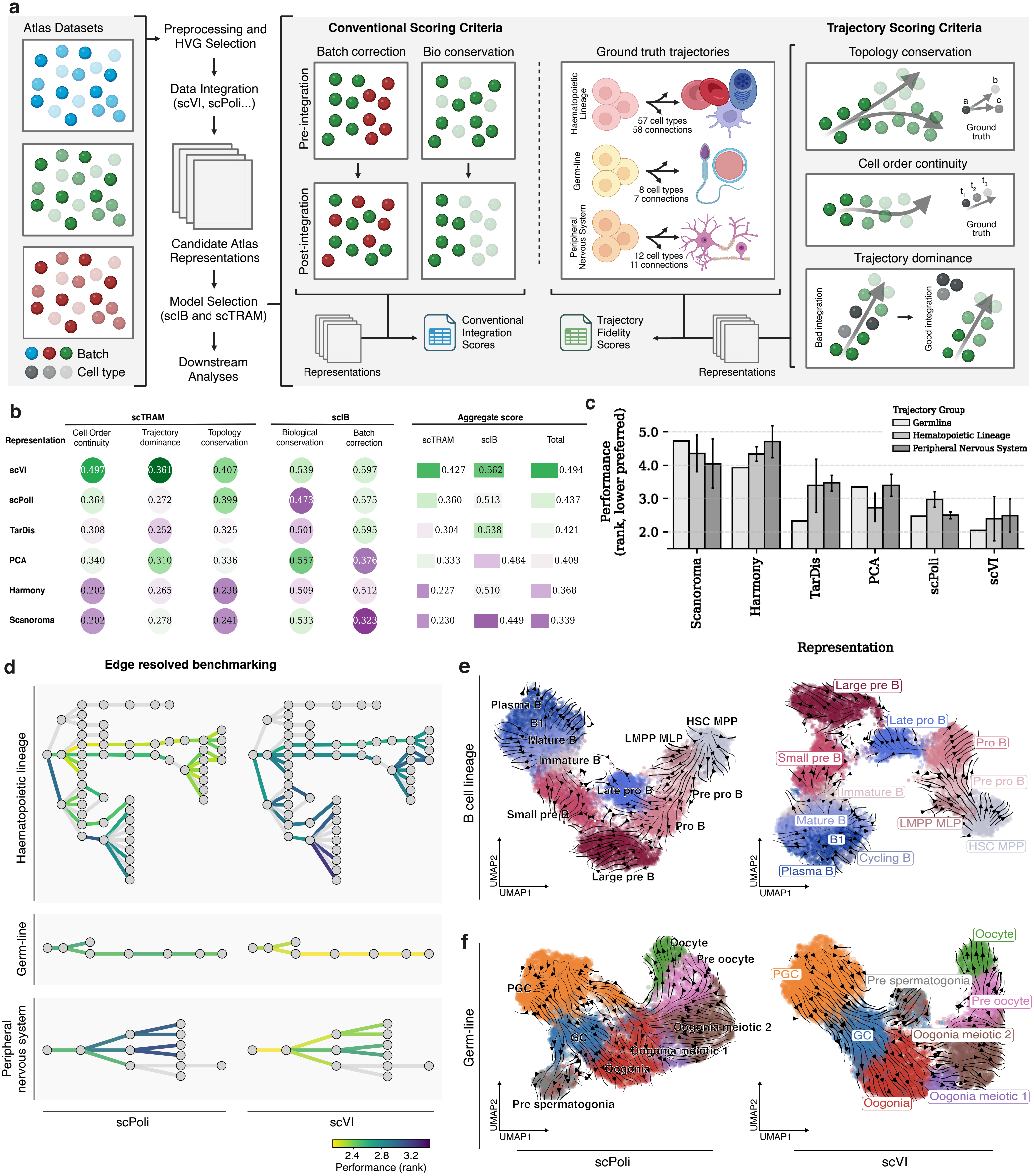

**Figure S3: Benchmarking atlas integration with trajectory fidelity**

**a)** Schematic of the benchmarking workflow combining scTRAM with the scIB protocol to evaluate developmental atlas integrations. Multiple single cell datasets were pre-processed and integrated (for example with scVI or scPoli) to generate candidate atlas embeddings. Conventional scIB metrics quantify batch correction and biological conservation. In parallel, scTRAM measures conservation of developmental trajectories against curated ground-truth lineages—haematopoietic, germline and peripheral nervous system—using 3 criteria: topology conservation, cell-order continuity and trajectory dominance. The resulting scores guide model selection for downstream analyses. **b)** Quantitative benchmarking of atlas integration methods using scTRAM and scIB. Aggregate columns show the mean of the 3 scTRAM metrics, the 2 scIB metrics and the overall mean across all 5 metrics (higher is better). scVI attains the highest combined score, indicating the best compromise between lineage preservation and batch integration. **c)** scTRAM performance stratified by trajectory group across integration methods. Bars show the mean trajectory-fidelity score (normalized 0–1; higher is better) for the 3 ground-truth lineages reference graphs for each method, with error bars indicating variability across scTRAM metrics. **d)** Edge-resolved trajectory fidelity for 2 atlas integrations. Reference lineage graphs for the 3 ground-truth lineage trajectories are shown with edges coloured by the scTRAM per-edge composite score (mean of topology conservation, cell-order continuity and trajectory dominance; scale 0.4–0.7, higher is better). Grey edges denote segments where scPoli and scVI are statistically indistinguishable (non-significant two-sided paired comparisons and Benjamini–Hochberg at 5% multiple-testing correction). Across the remaining edges, scVI attains higher fidelity on most segments and branch points, indicating graded preservation of developmental progression. **e)** UMAP visualisation of the B-cell differentiation trajectory in 2 atlas embeddings. Cells are coloured by pseudotime from early progenitors (pink/red) to terminal B states (blue); black streamlines/arrows depict directional flow inferred with the CellRank pseudotime kernel. Relative to scPoli, the scVI embedding shows a more coherent progression from HSC/MPP and LMPP/MLP through pro-B and pre-B stages to mature and plasma B cells. **f)** UMAP visualisation of the germline lineage in the García component dataset versus the integrated atlas. Cells are coloured by curated cell-type annotations; black streamlines indicate directional flow inferred with the CellRank pseudotime kernel. The atlas yields a more continuous progression from primordial germ cells through oogonia and pre-oocyte to oocyte, and better resolves branching towards pre-spermatogonia and granulosa cells.

### Figure S4: A comprehensive, adaptable human developmental resource

a

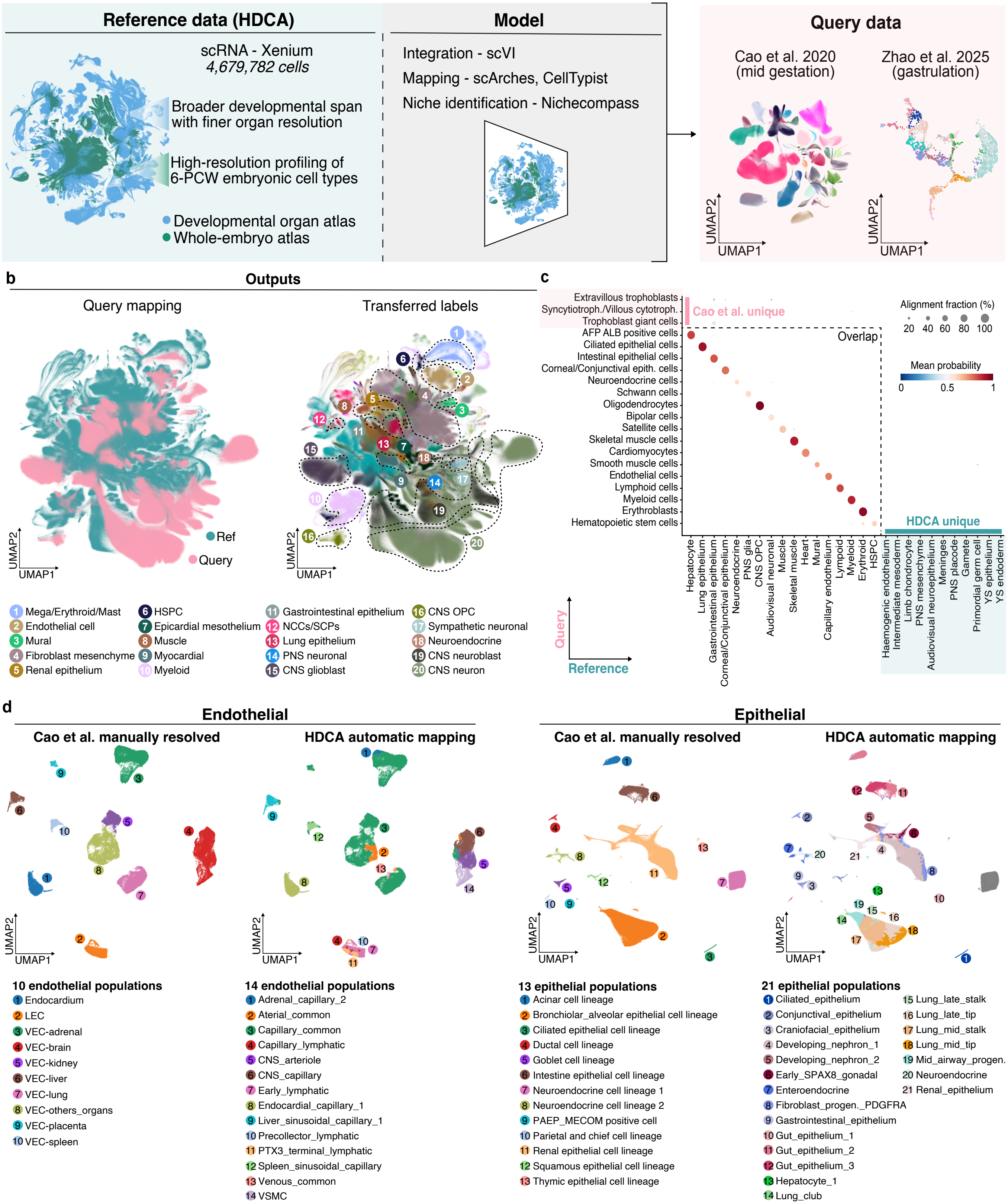


| | # Cells | # Total cell types | # Genes expressed | # UMIs |
|---|---|---|---|---|
| Cao *et al.* | 89k | 10 | 466 | 648 |
| HDCA | 106k | 14 (40) | 3233 | 12818 |

| | # Cells | # Total cell types | # Genes expressed | # UMIs |
|---|---|---|---|---|
| Cao *et al.* | 282k | 13 | 629 | 991 |
| HDCA | 222k | 21 (40) | 2980 | 11464 |

**Figure S4: A comprehensive, adaptable human developmental resource**

**a)** Schematic overview of reference model construction and query data mapping. Two complementary models, including a deep learning-based scArches model and an expression-based CellTypist model, are used to map two query datasets. A NicheCompass-based model for niche detection is also provided. **b)** Left: UMAP visualisation of reference and query cells following scArches-based mapping. Right: Transferred cell type labels overlaid onto the UMAP colored by selected major cell types. **c)** Scatter plot showing cell type label transfer from the reference to the query dataset[47], highlighting shared and non-overlapping populations between datasets. Color and size of the dot denote the mean prediction probability and cell type alignment fraction, respectively. **d**) UMAPs of endothelial, and epithelial cells from Cao et al., colored by author-refined annotations (left) and by cell type labels transferred using the HDCA reference mapping model (right). Tables below summarize the number of cells, total annotated cell types (the number in parentheses denotes the total number of cell types present in the HDCA), and average expressed genes and UMI counts between the two datasets.

**Figure S5: Cell type label transfer from the HDCA reference to the mid-gestation query data**

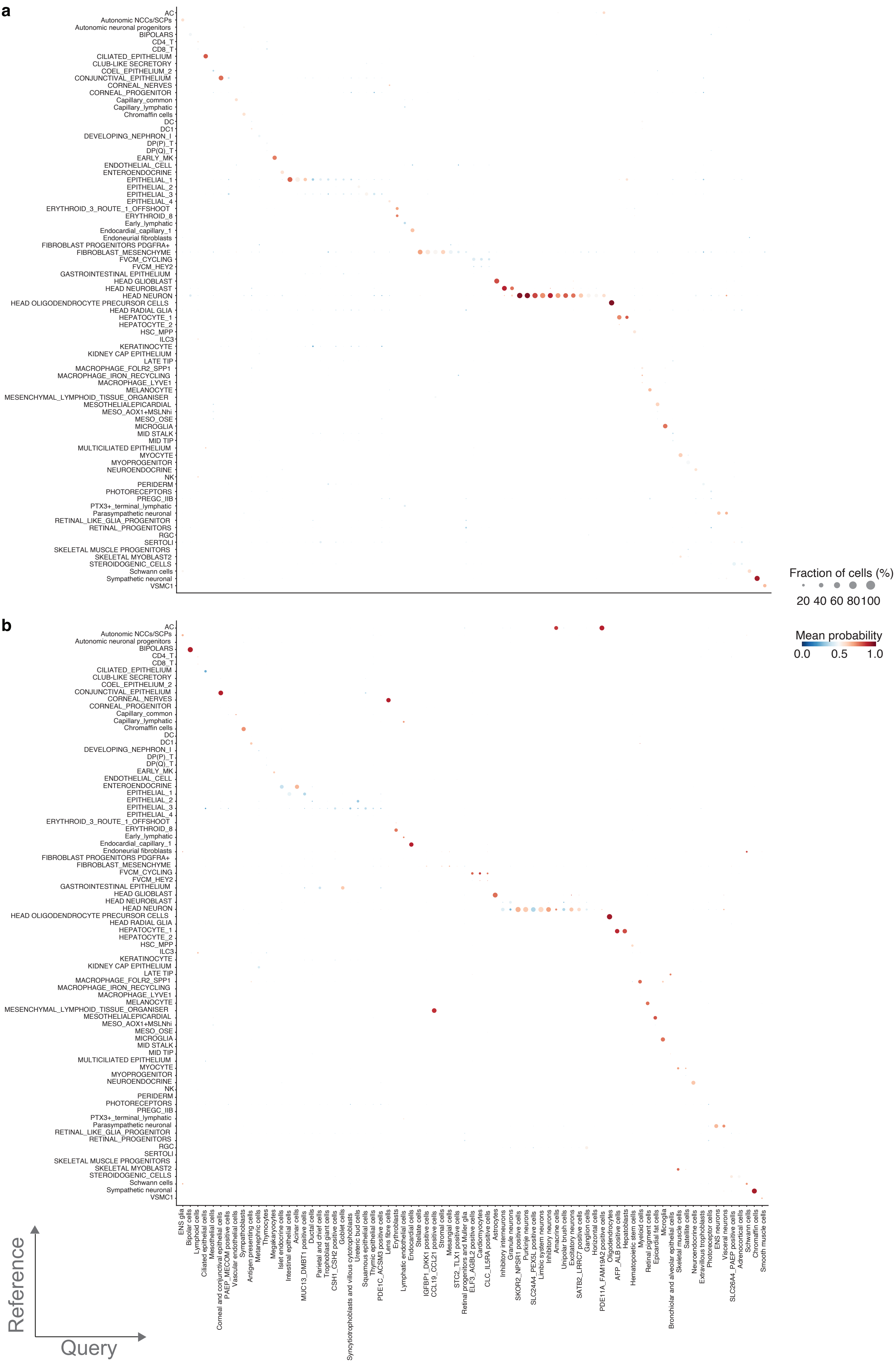

**Figure S5: Cell type label transfer from the HDCA reference to the mid-gestation query data**

**a)** Scatter plot showing cell type label transfer from the reference to the query dataset[47] based on the scArches model. Color and size of the dot denote the mean prediction probability and cell type alignment fraction, respectively. **b)** As with a), but based on the Celltypist model.

#### Figure S6: Reference mapping of early human embryogenesis data

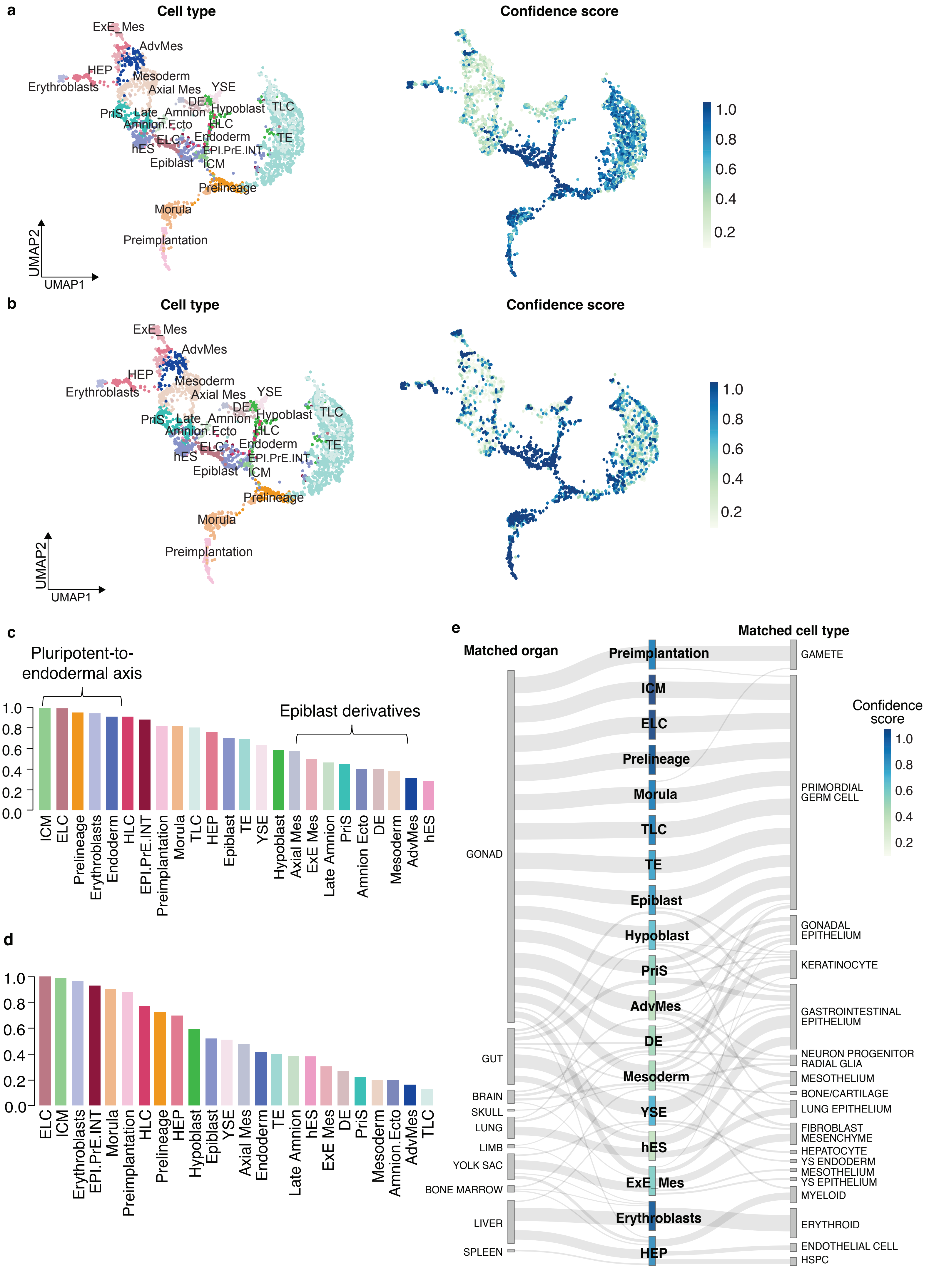

**Figure S6: Reference mapping of early human embryogenesis data**

**a)** Left: UMAP visualisation of the zygote-to-gastrula query dataset coloured by original author cell annotations. Right: Mapping confidence scores overlaid onto the UMAP following reference mapping by the scArches model. **b)** As with a), but by the CellTypist model. **c)** Bar plot summarising mean mapping confidence scores across cell types based on the scArches model, highlighting high confidence along the pluripotent-to-endodermal axis and lower confidence among epiblast-derived populations. **d)** As with c), but based on the CellTypist model. **e)** Sankey diagram depicting correspondence between query cell types (middle) and matched reference cell types (upper) and organs (lower).
