## Supplementary Figure 7 for "An integrated single cell and spatial omics atlas of human prenatal development" (from another version of this preprint: version 1, 2026-04-01)

Figure S7: An unsupervised systematic decomposition of human embryos into niches

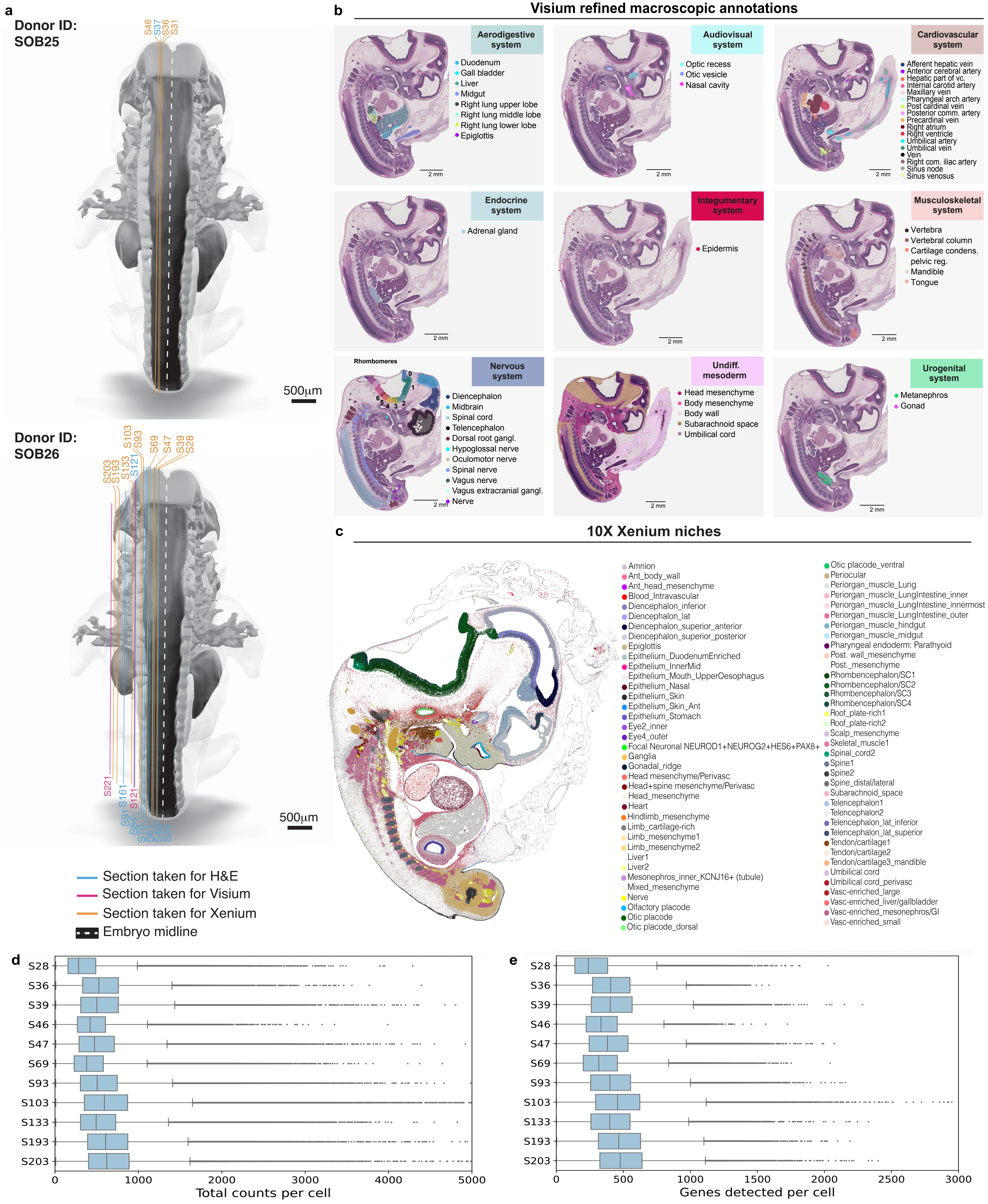

**Figure S7: An unsupervised systematic decomposition of human embryos into niches**

**a)** Adapted schematic taken from https://hdbratlas.org/carnegie-stages/CS17.html showing an example of the posterior side of a CS17 human embryo. The lines indicate the estimated locations of where FFPE embryo sections (from samples SOB25, SOB26, see **Supplementary Table 5** for full sample metadata) were sliced for analysis for H&E, and Xenium and Visium 10x Genomics platforms. The scale bar provided is an estimate based on the average width of the CS17 embryo (4mm). The dotted lines were drawn based on the distance from the midline (distances provided in **Supplementary Table 5**). **b)** Grid layout of broad anatomical systems within the 10X Visium data. For each broad anatomical system, the Visium shown highlights refined labels for each system respectively on H&E of a sagittal slice of an embryo. The scale bar for all images is 2mm. **c)** Xenium niches from **Fig. 2b**, section 28, showing labels for all legends. **d-e)** Distribution (box plot) of total counts per cell (d) and number of genes per cell (e) for each Xenium 5k section. Box-plot elements: center line, median; box limits, upper and lower quartiles; whiskers, 1.5x interquartile range; points, outliers.
