## Supplementary Figure 8 for "An integrated single cell and spatial omics atlas of human prenatal development" (from another version of this preprint: version 1, 2026-04-01)

Figure S8: An unsupervised systematic decomposition of human embryos into niches and comparison to expert histological annotations

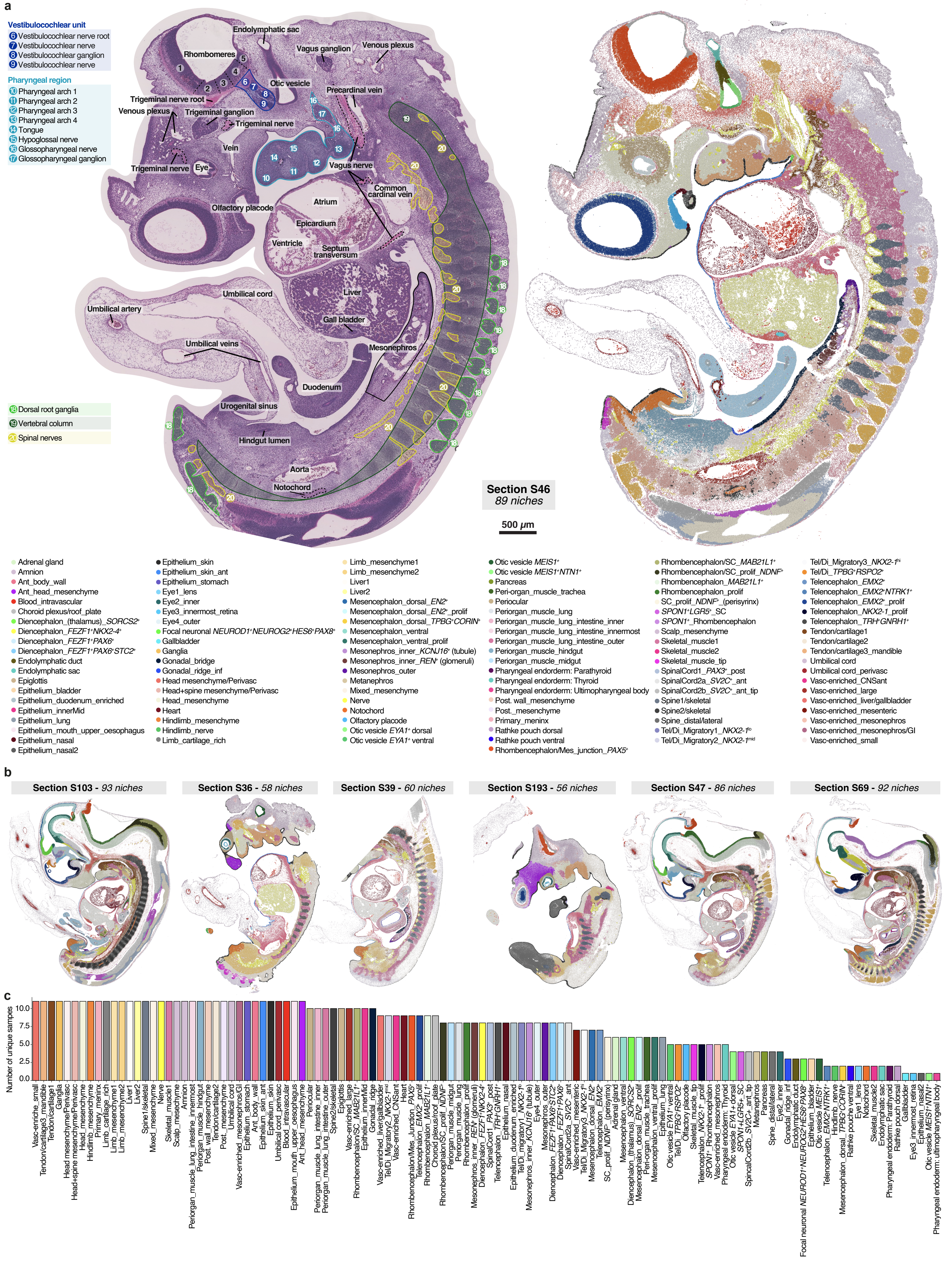

**Figure S8: An unsupervised systematic decomposition of human embryos into niches and comparison to expert histological annotations.**

**a)** Left: Expert histological annotations of human embryo section S46. Right: Tissue section profiled by Xenium 5k section S46, coloured by 89 of the 114 niches (see **Supplementary Table 13** for niche distribution across all sections). **b)** Remaining 6 whole embryo Xenium 5k reference sections (S103, S36, S39, S193, S47, S69; not shown in **Fig. 2b** (S28) or panel a above (S46)) coloured by niche. Colours as per panel a. **c)** Bar plot showing the number of unique sections containing the niche (minimum of 10 cells) - reference sections only. Some niches were only captured in 1 section, such as the gallbladder and ultimopharyngeal body, highlighting the value of comprehensive sagittal sections.
