## Supplementary Figures 9-10 for "An integrated single cell and spatial omics atlas of human prenatal development" (from another version of this preprint: version 1, 2026-04-01)

**Figure S9: Focal and distributed niche locations**

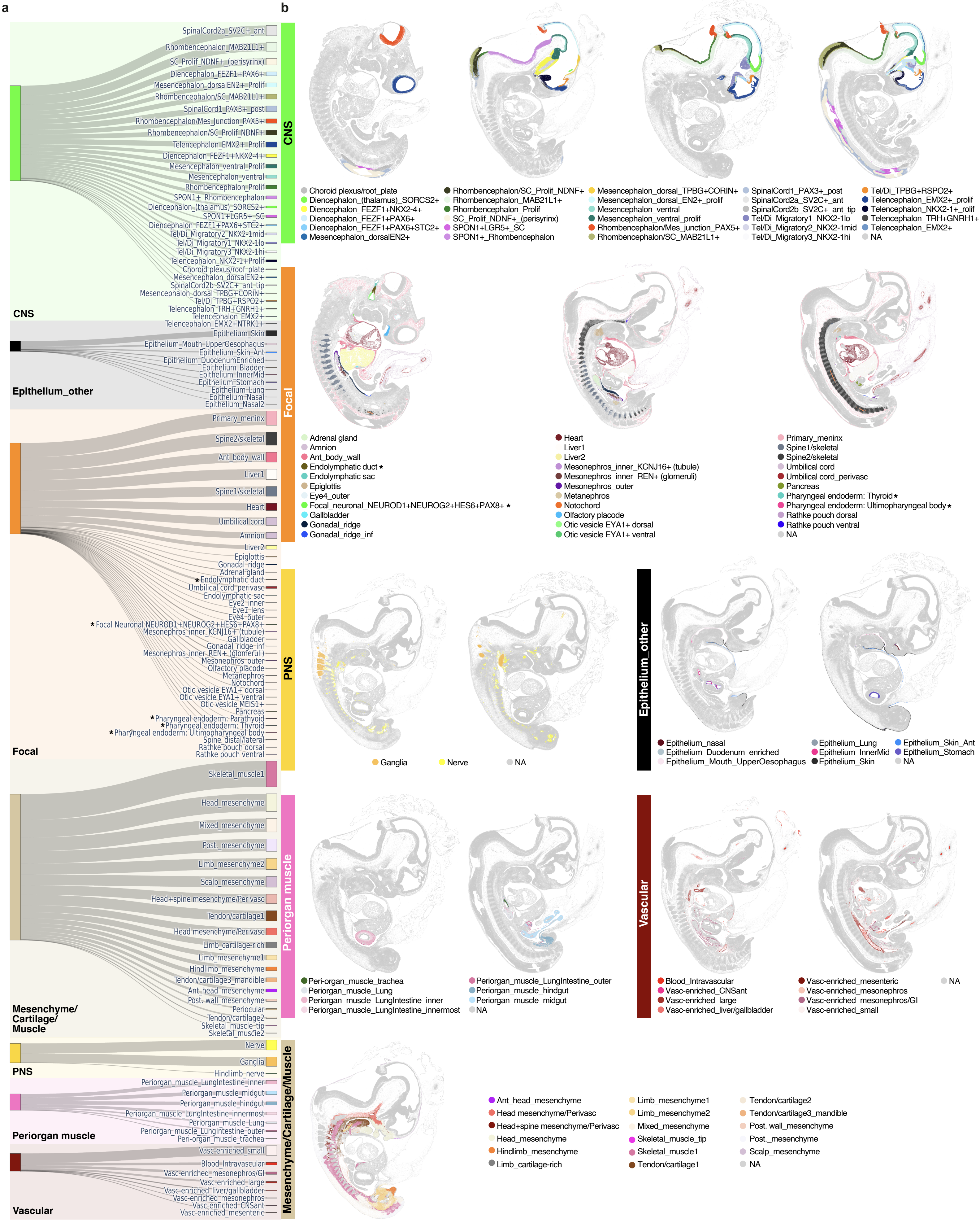

**Figure S9: Focal and distributed niche locations.**

**a)** Sankey plot showing the niches contained within each broad group (focal, vascular-enriched, mesenchyme/cartilage/tendon. epithelium_other, central nervous system (CNS), peripheral nervous system (PNS), peri-organ muscle). * indicates focal niche not annotated from H&E. **b)** Whole embryo sections coloured by niches within each broad group to more clearly illustrate the distribution of niches within these groups. Focal annotations corresponded to well-defined structures. (Sections used: focal S46, S69, and S103; PNS S28 and S102; CNS S28, S46, S69, and S103; Mesenchyme/cartilage/muscle S47; Epithelium_other S28 and S69; Periorgan muscle S28 and S47; Vascular rich S69 and S103). Note that due to showing limited sections, not all niches are shown.

Figure S10: Known niches - molecular drivers and patterning

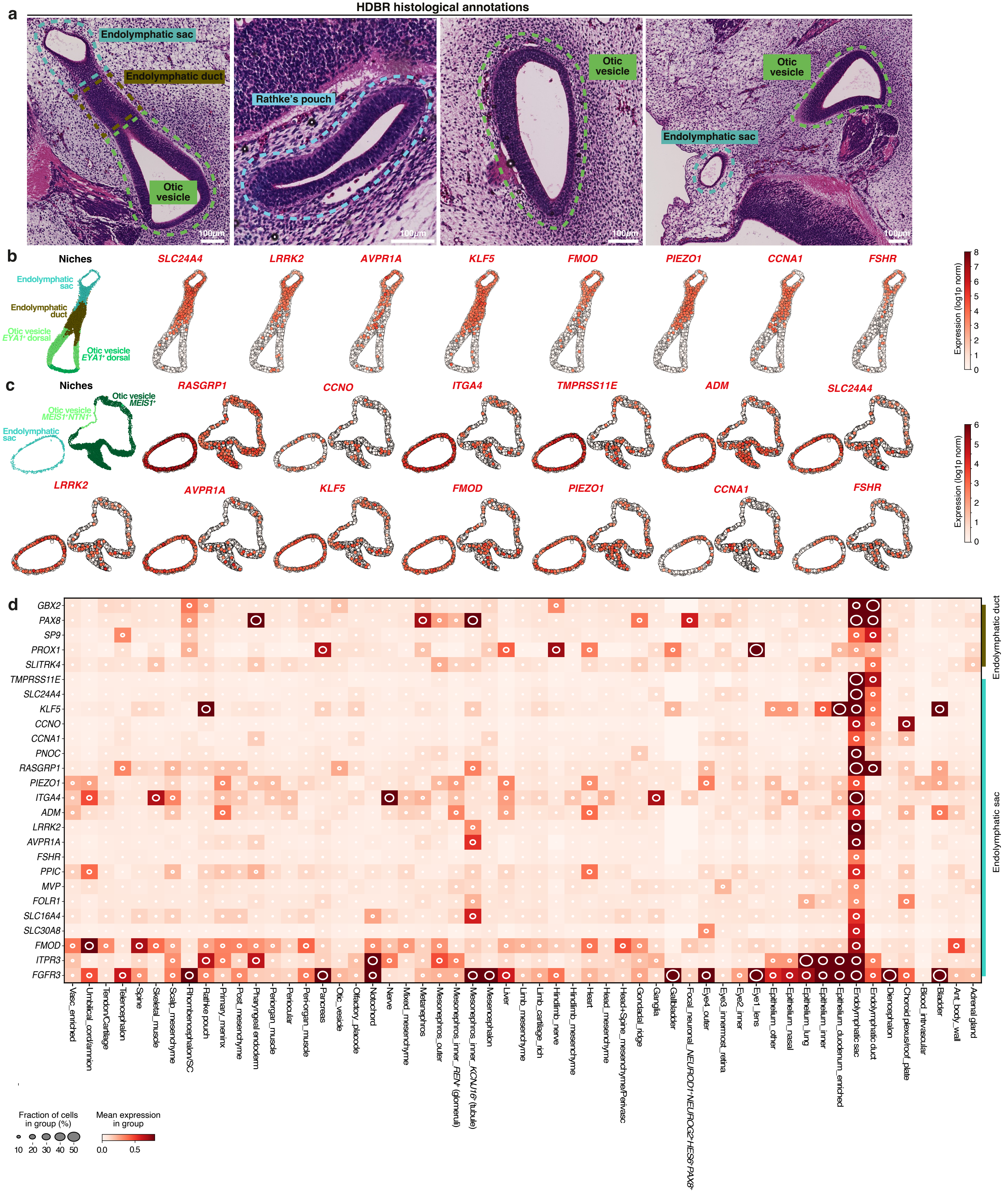

**Figure S10: Known niches - molecular drivers and patterning**

**a)** Expert histological annotations of human embryo sections, showing the endolymphatic sac adjacent to the otic vesicle (left). Rathke's pouch (middle) and otic vesicle (right), but without evidence of patterning from H&E alone. **b)** Left: Xenium 5k capture of the otic vesicle adjacent to the endolymphatic sac from S46. Right: gene expression of differential expressed genes within the endolymphatic sac. **c)** Left: Xenium 5k capture of the otic vesicle and endolymphatic sac in a query section for validation (S203). Right: gene expression of differential expressed genes within the endolymphatic sac. **d)** Combined heatmap-dotplot showing expression for endolymphatic sac-associated genes across all niches, with some niches collapsed into a single group (e.g. vascular-enriched, Rathke's pouch).
