## Supplementary Figures 11-12 for "An integrated single cell and spatial omics atlas of human prenatal development" (from another version of this preprint: version 1, 2026-04-01)

### Figure S11: New detail within known histological structures

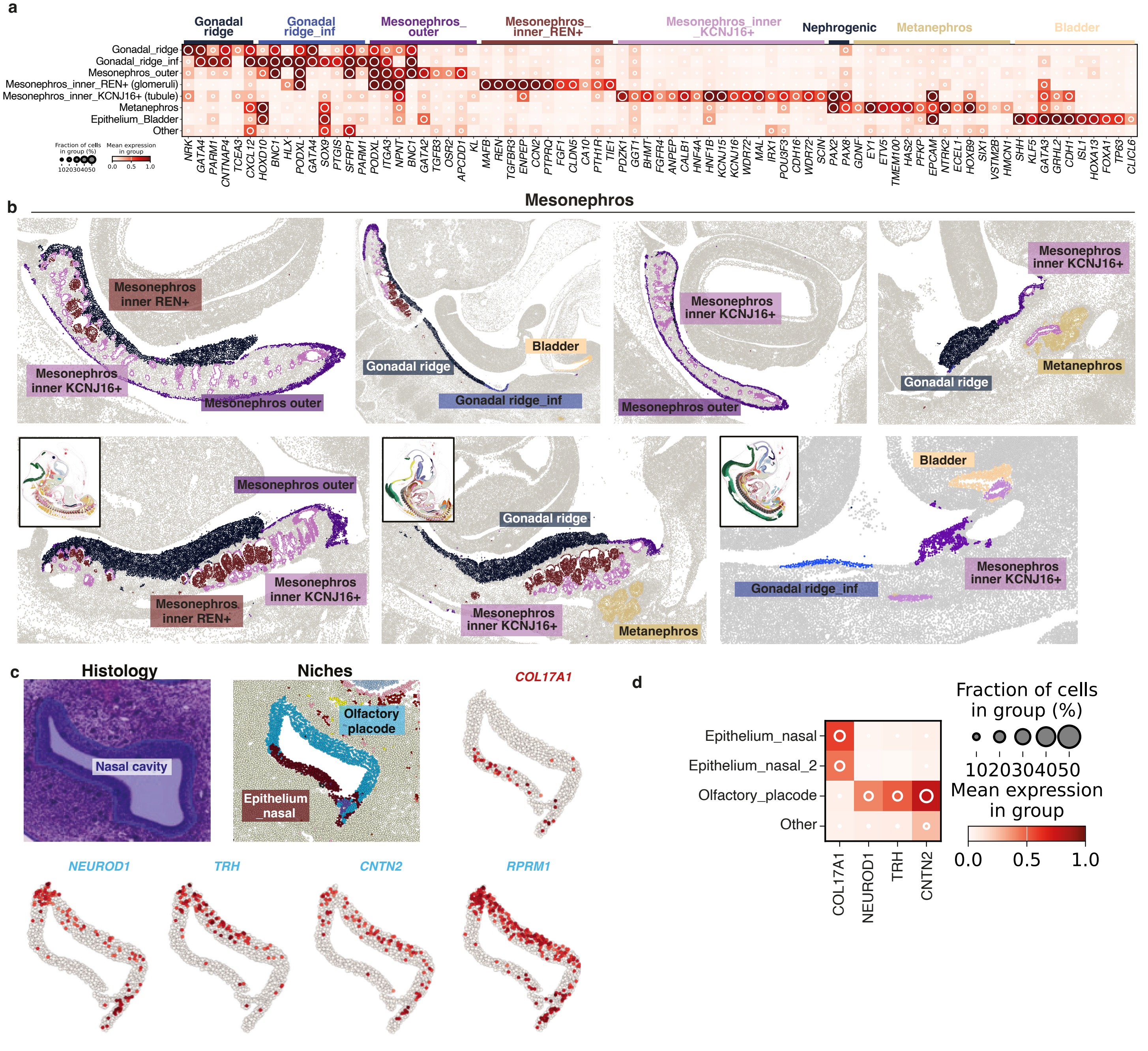

**Figure S11: New detail within known histological structures**

**a)** Combined heatmap-dotplot showing niche differential gene expression for gonad and nephric niches. **b)** Mesonephros/metanephros region captured in embryo sections profiled by Xenium 5k, coloured by niche for mesonephros/metanephros/bladder niches. Top row: S39, S46, S47, S133. Bottom row: S93, S69, S103. Inlet: whole embryo section coloured by all niches. **c)** Expert histological annotations of nasal cavity and nearby region coloured by Xenium 5k niche (in section S28), showing olfactory placode within the nasal cavity. Expression of niche differentially expressed genes for nasal epithelium (*COL17A1*) and olfactory placode (*NEUROD1*, *TRH*, *CNTN2*, *RPRM1*). **d)** Combined heatmap-dotplot of expression of niche differentially expressed genes for nasal epithelium (*COL17A1*) and olfactory placode (*NEUROD1*, *TRH*, *CNTN2*, *RPRM1*) in all sections (log1p norm expression).

### Figure S12: Novel (unannotated) niches within human embryo

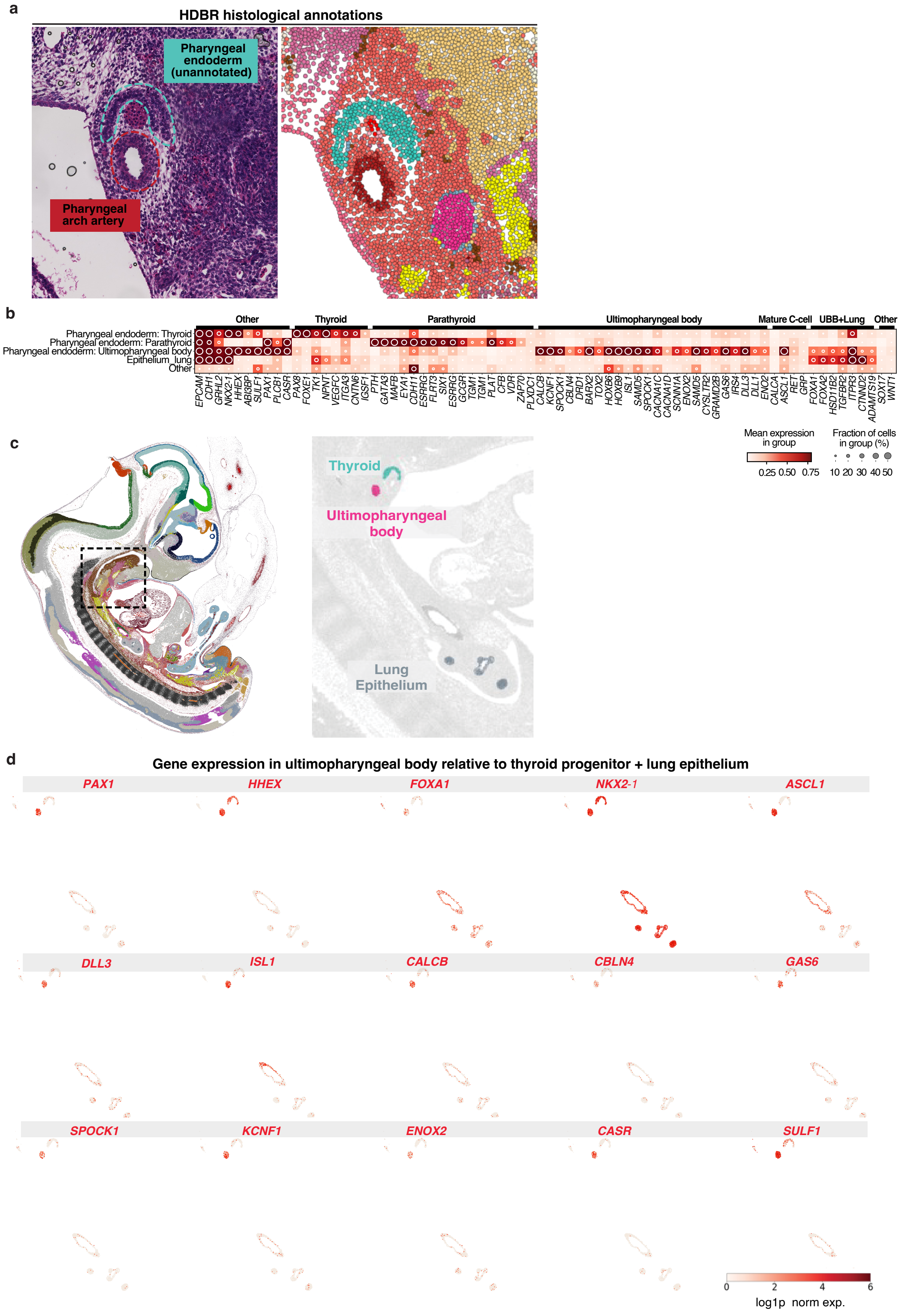

**Figure S12: Novel (unannotated) niches within human embryo**

**a)** Expert histological annotations of pharyngeal artery (left, section 91), without annotated pharyngeal endoderm. Nearby tissue region coloured by Xenium 5k niche (right, section 103). **b)** Combined heatmap-dotplot showing niche differential gene expression for pharyngeal endoderm niches and lung epithelium. Log1p norm expression. **c)** Xenium 5k whole embryo section coloured by niche, with zoomed in region showing pharyngeal endoderm and lung epithelium niches from section 103 (top row). **d)** Expression of selected genes in the pharyngeal endoderm and lung epithelium niches.
