## Supplementary Figures 13-18 for "An integrated single cell and spatial omics atlas of human prenatal development" (from another version of this preprint: version 1, 2026-04-01)

### Figure S13: Reference-query mapping to allow automated annotation and identify novel niches not present in the reference and niche-level investigation of disease-associated genes

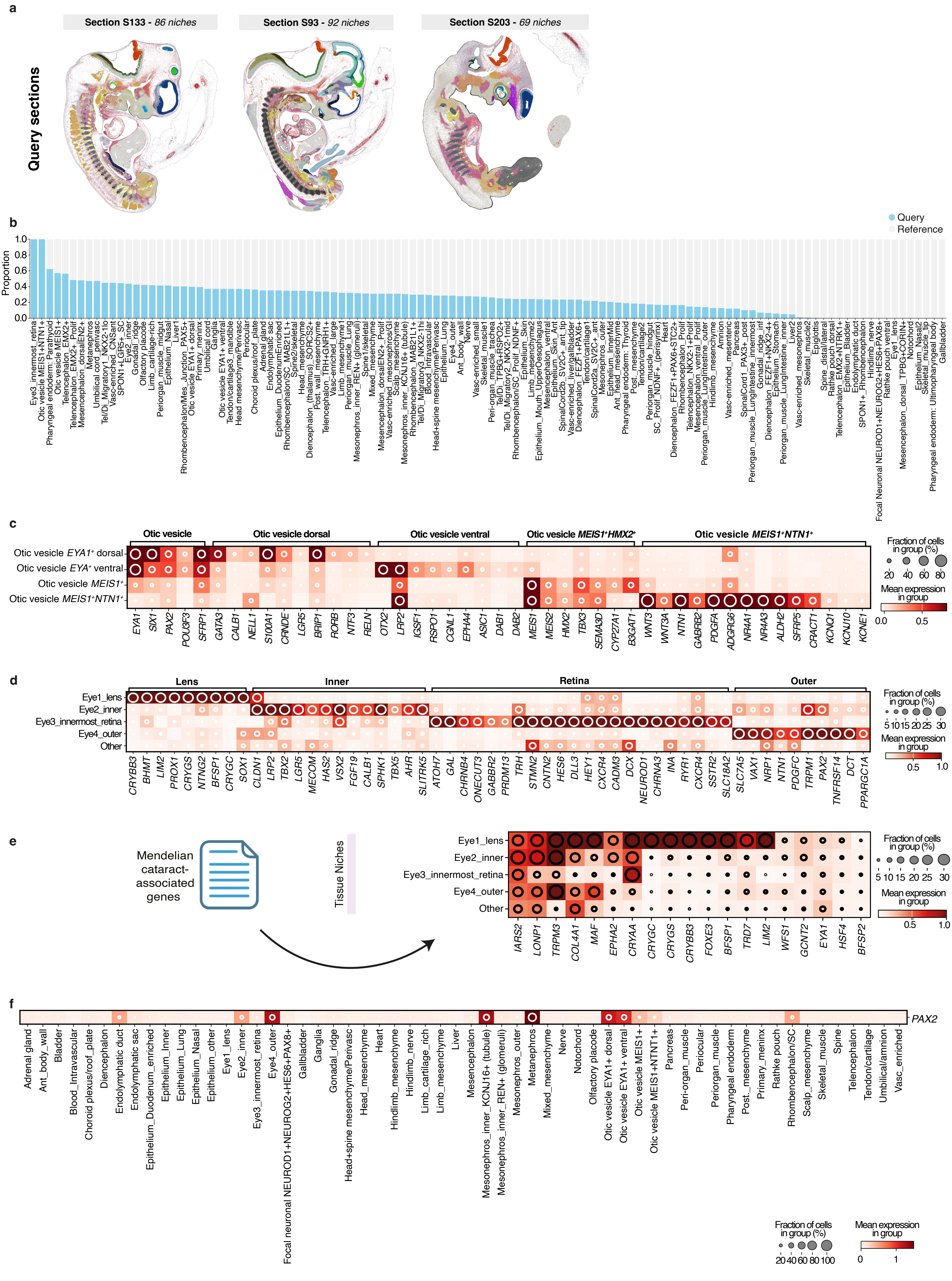

**Figure S13: Reference-query mapping to allow automated annotation and identify novel niches not present in the reference and niche-level investigation of disease-associated genes**

**a)** Xenium 5k whole embryo section for 3 query sections after reference-query mapping (S133, S93, S203), coloured by some of the 114 niches (colours as in **Fig. S8a;** see **Supplementary Table 13** for niche distribution across all sections). **b)** Proportion of each niche from reference and query samples, showing identification of a novel retinal niche in the query samples. **c**) Combined heatmap-dotplot of expression of niche differentially expressed genes for otic vesicle niches. **d)** Combined heatmap-dotplot of expression of niche differentially expressed genes for eye niches. **e)** Combined heatmap-dotplot of expression of genes associated with congenital cataracts in the eye and other niches. **f)** Combined heatmap-dotplot of expression of *PAX2* across niches, with some niches collapsed into a single group (e.g. vascular-enriched, Rathke's pouch).

#### Figure S14: Undefined progenitor populations

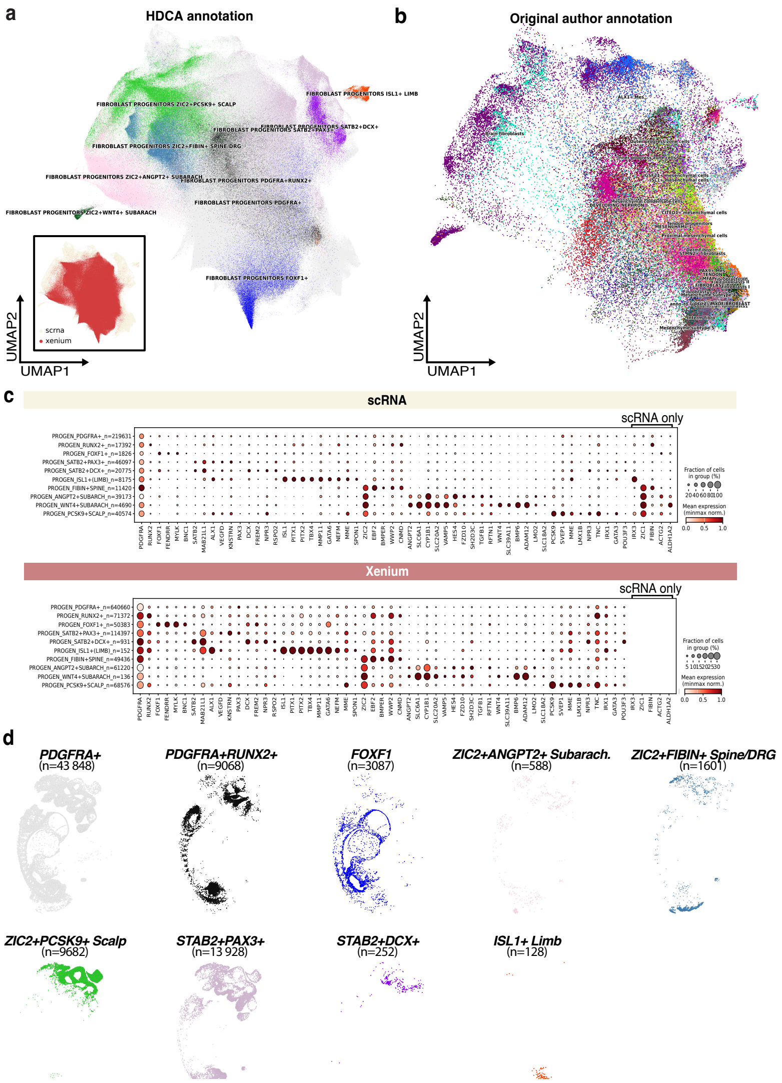

**Figure S14: Undefined progenitor populations.**

**a)** UMAP embedding of originally undefined progenitor cells, coloured by cell type. Inset: coloured by technology. **b)** as in panel a, but coloured by original author annotation (minimum of 1000 cells). **c)** Dot plot of marker gene expression for undefined progenitors shown in scRNA-seq and Xenium 5k. **d)** Xenium-5k whole embryo showing the location of each undefined progenitor population (section S46).

### Figure S15: Lateral plate mesoderm and mesothelial populations

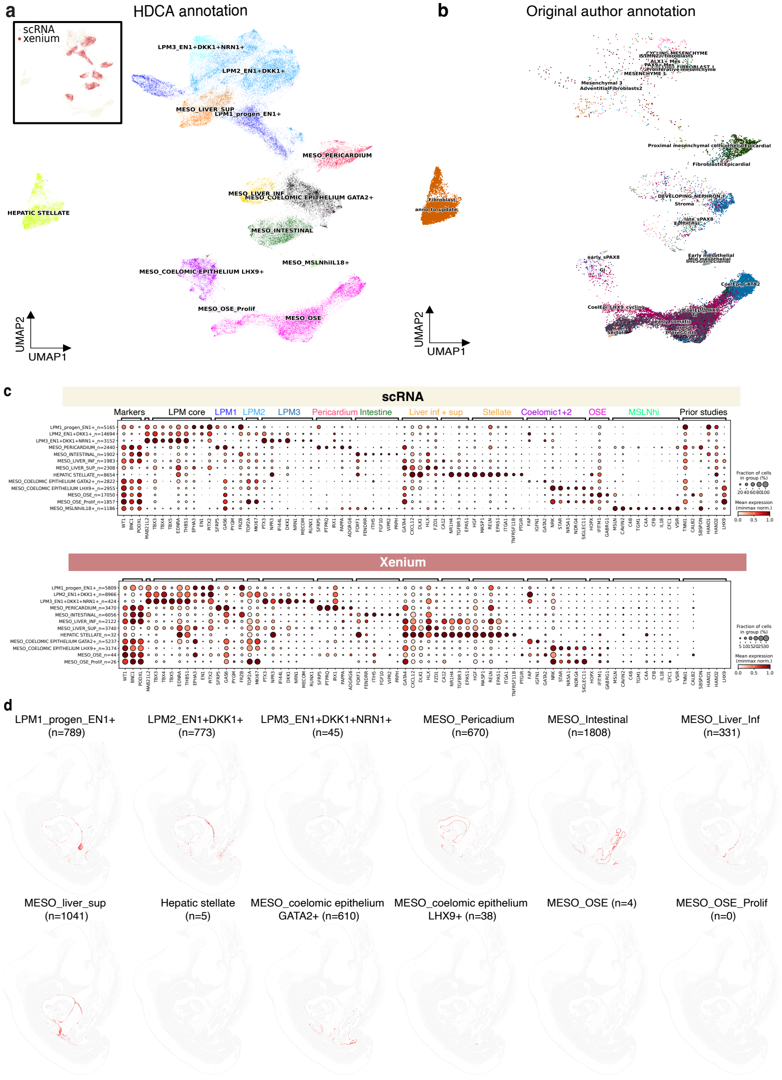

**Figure S15: Lateral plate mesoderm and mesothelial populations.**

**a)** UMAP embedding of mesothelial group, coloured by cell type. Inlet: coloured by technology. **b)** UMAP embedding of mesothelial group, coloured by original author annotation (minimum of 20 cells). **c)** Dot plot of marker gene expression for mesothelial cells shown in scRNA-seq and Xenium 5k. **d)** Xenium-5k whole embryo showing the location of each mesothelial population (section S103).

Figure S16: Extra-embryonic umbilical cord and amnion populations

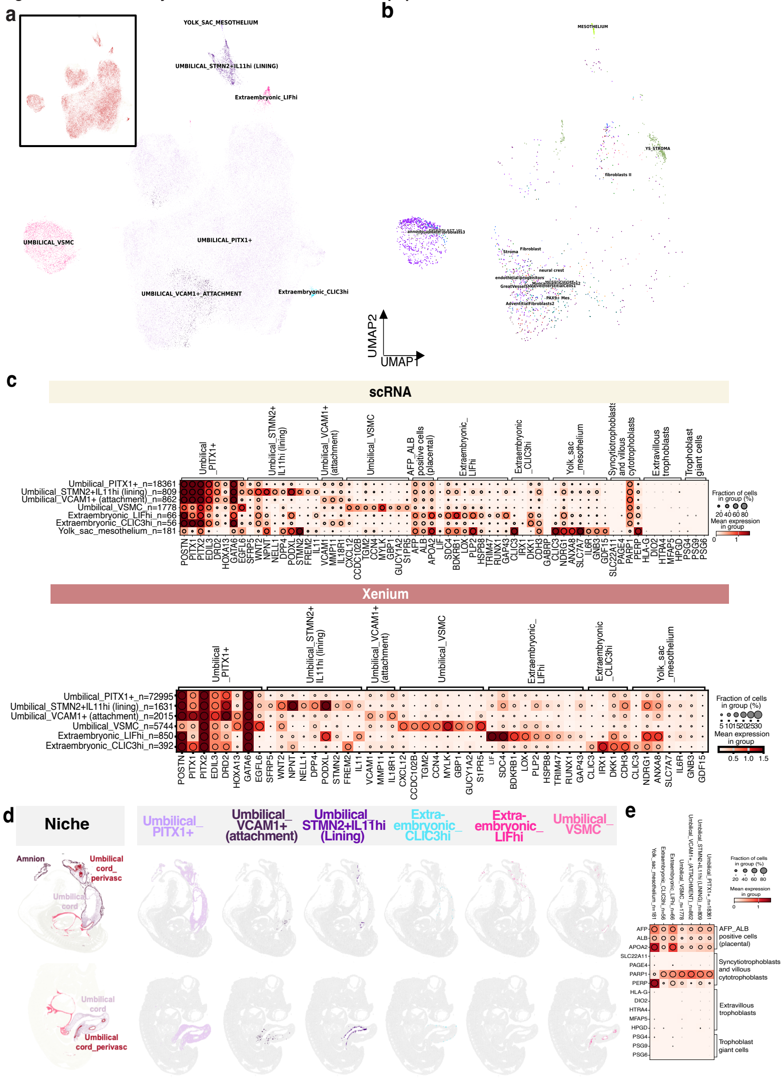

**Figure S16: Extra-embryonic umbilical cord and amnion populations.**

**a)** UMAP embedding of umbilical/extraembryonic group, coloured by cell type. Inlet: coloured by technology. **b)** UMAP embedding of umbilical/extraembryonic group, coloured by original author annotation (minimum of 30 cells). **c)** Dot plot of marker gene expression for umbilical and extraembryonic cells shown in scRNA-seq and Xenium 5k. **d)** Left: 2 whole embryo sections profiled by Xenium 5k, showing location of amnion, umbilical cord, and umbilical cord_perivascular niche locations (sections S69 top and S46 bottom). Right: Same whole embryo sections showing the location of umbilical/extraembryonic populations. **e)** Dot plot of marker gene expression for trophoblast expression from Cao et al[47] (scRNA-seq data).

### Figure S17: Undefined progenitor cell distributions and origins

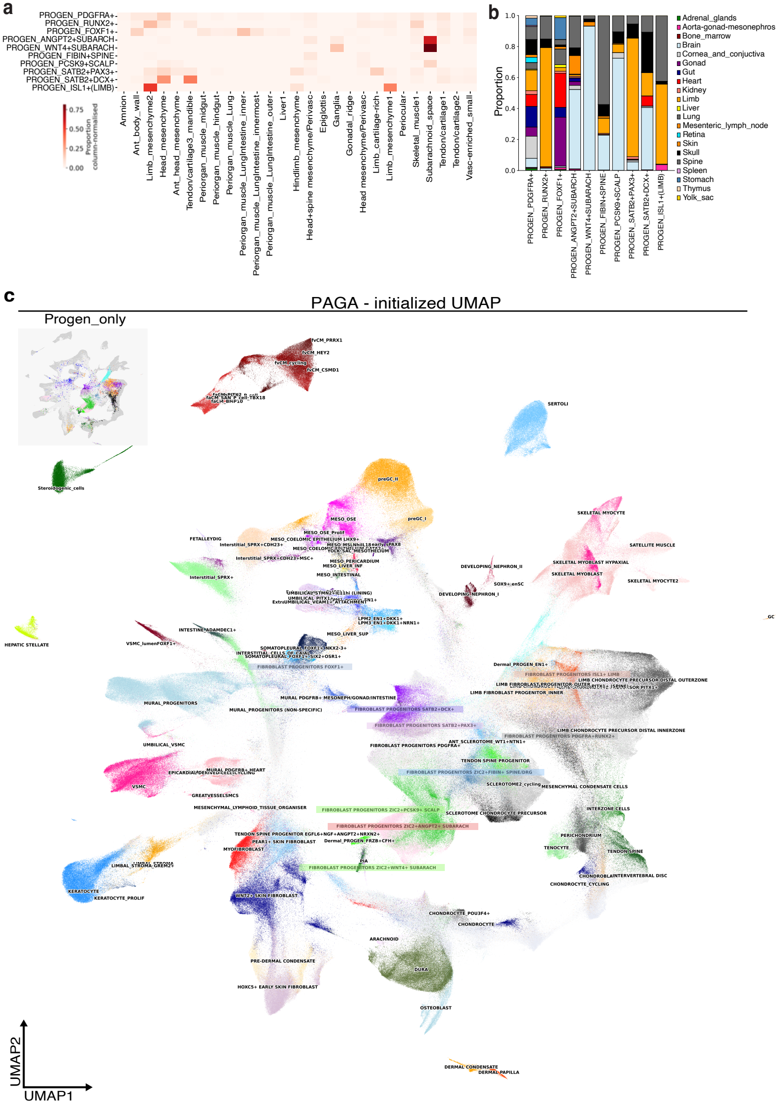

**Figure S17: Undefined progenitor cell distributions and origins.**

**a)** Heatmap showing distribution of undefined progenitor cells by niche. **b)** Bar plot showing proportion of undefined progenitor cells by organ of origin in scRNA-seq data (from integrated atlas). **c)** UMAP embedding of scRNA-seq data, coloured by cell type, with PAGA initialisation. Fibroblast progenitors $ISL1^{+}$ (limb) were located with limb fibroblast populations. $ZIC2^{+}FIBIN^{+}$ (Spine/DRG) was in the spinal cluster. $ZIC2^{+}WNT4^{+}$ was located with meningeal populations. Fibroblast progenitors $FOXF1^{+}$ were located with somatopleural populations.

### Figure S18: Lateral plate mesoderm heterogeneity and mesothelium formation

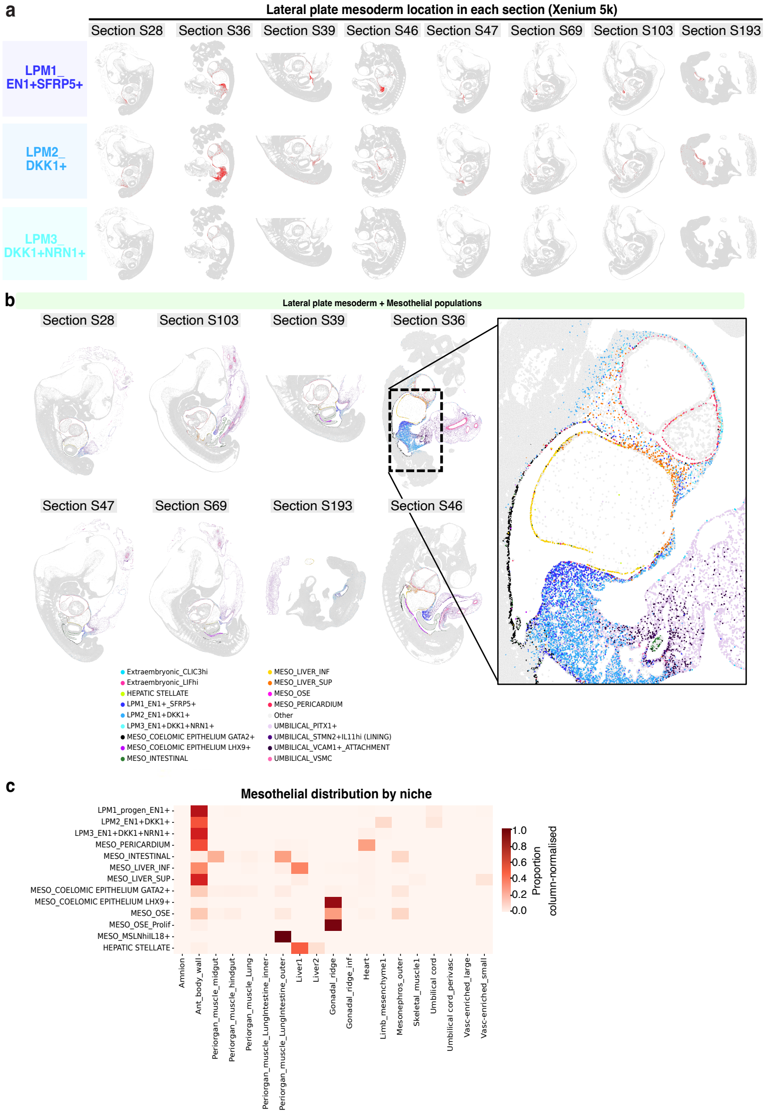

**Figure S18: Lateral plate mesoderm heterogeneity and mesothelium formation**

**a)** Location of LPM subtypes across all 8 reference whole embryo sections (Xenium-5k) (left to right, S28, S36, S39, S46, S47, S69, S103, S193). **b)** Location of LPM, mesothelial, and umbilical cell types in all reference embryo sections. Top row (left to right): sections S28, S103, S39, S36. Bottom row: S47, S69, S193, S46. **c)** Heatmap showing distribution of mesothelial cells by niche.
