## Supplementary Figures 19-23 for "An integrated single cell and spatial omics atlas of human prenatal development" (from another version of this preprint: version 1, 2026-04-01)

### Figure S19: Decoding cranial embryo mesoderm niches

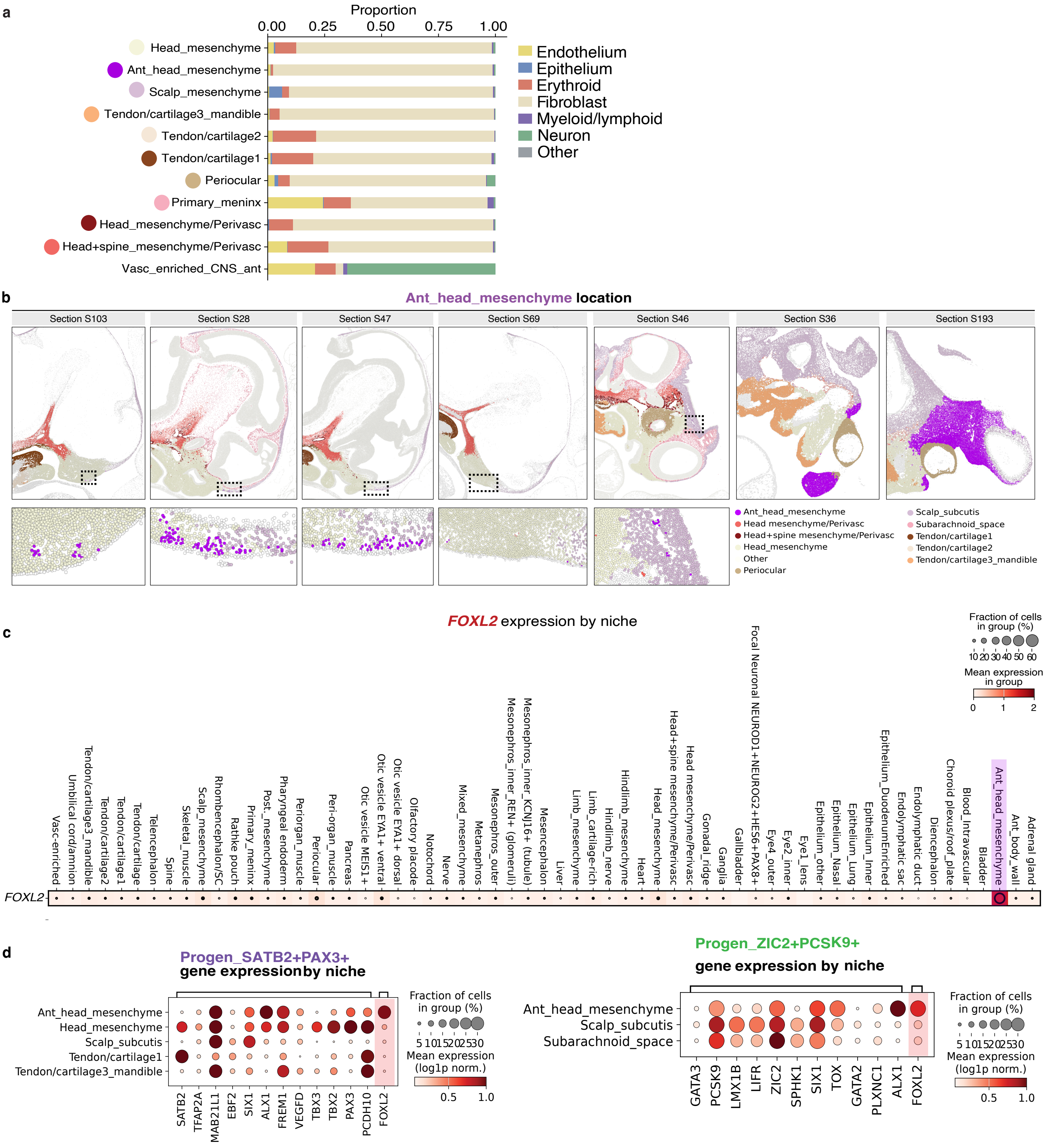

**Figure S19: Decoding cranial embryo mesoderm niches**

**a)** Barplot showing proportion of cells in head region niches that were predominantly formed of fibroblast cells. **b)** Xenium 5k profiling of head region in whole embryo reference sections, excluding section S39 reference section which lacks head (left to right: S103, S28, S47, S69, S46, S36, S193) coloured by head mesenchyme niches. Dotted boxes highlight the 'Ant_head_mesenchyme niche', with zoomed in regions shown below the dotted box. **c)** Combined heatmap-dotplot of *FOXL2* expression across all niches. **d)** Dotplot showing core marker genes and *FOXL2* expression in fibroblast progenitors that were most common in the Ant_head_mesenchyme niche (Progen SATB2+PAX3+ and Progen_ZIC2+PCSK9+).

### Figure S20: Novel mesenchymal progenitors are recapitulated in human organoid model

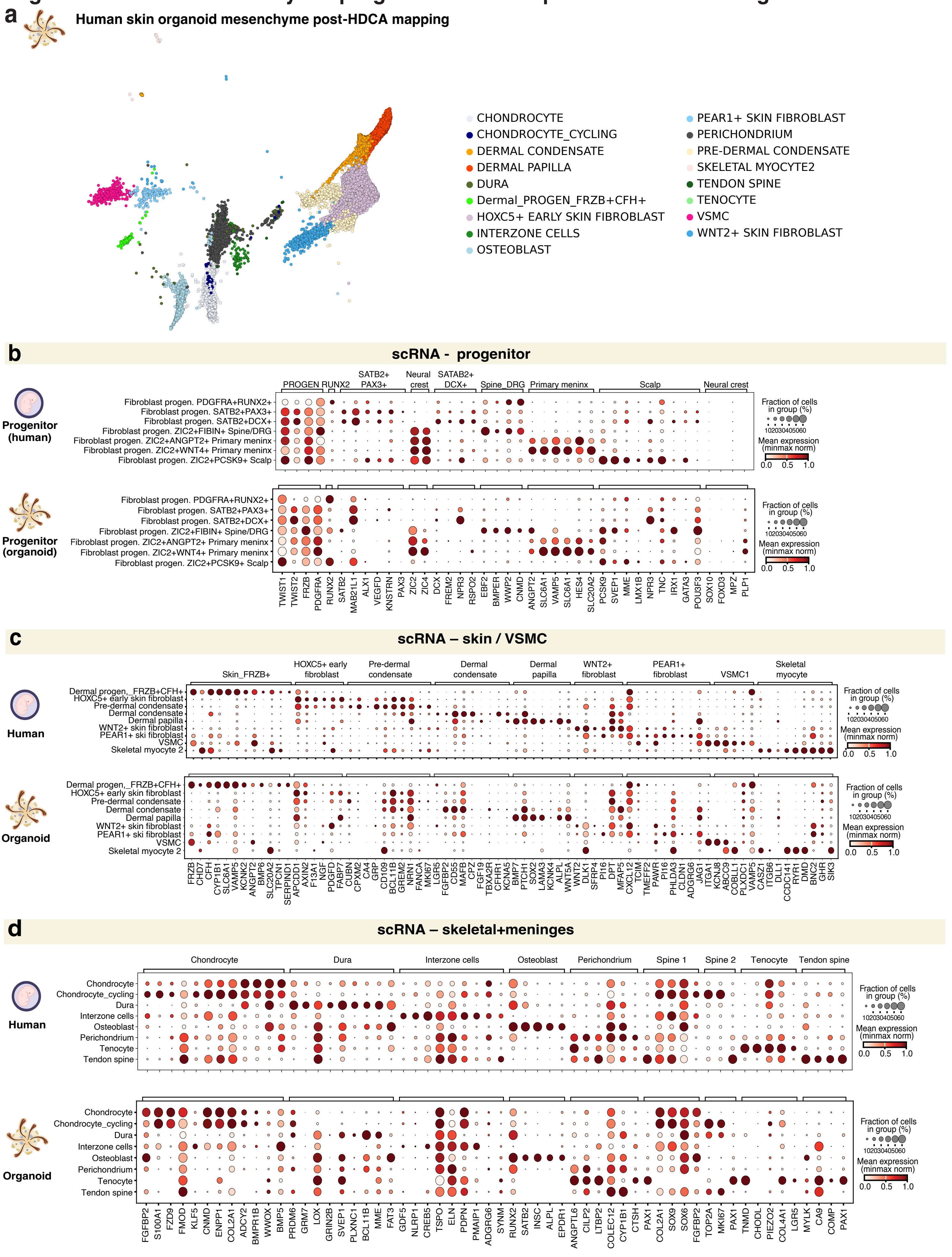

**Figure S20: Novel mesenchymal progenitors are recapitulated in human organoid model**

**a)** UMAP embedding of non-undefined progenitor cells in human skin organoid after reference-query mapping, coloured by cell type. **b)** Dot plots of marker gene expression for undefined progenitors in scRNA-seq data in human developmental cell atlas (top) and human skin organoid (bottom). **c)** Dot plots of marker gene expression for skin populations and vascular smooth muscle cells in scRNA-seq data in human developmental cell atlas (top) and human skin organoid (bottom). **d)** Dot plots of marker gene expression for skeletal and meningeal populations in scRNA-seq data in human developmental cell atlas (top) and human skin organoid (bottom).

### Figure S21: Specialisation of organ-specific endothelial cells during human development

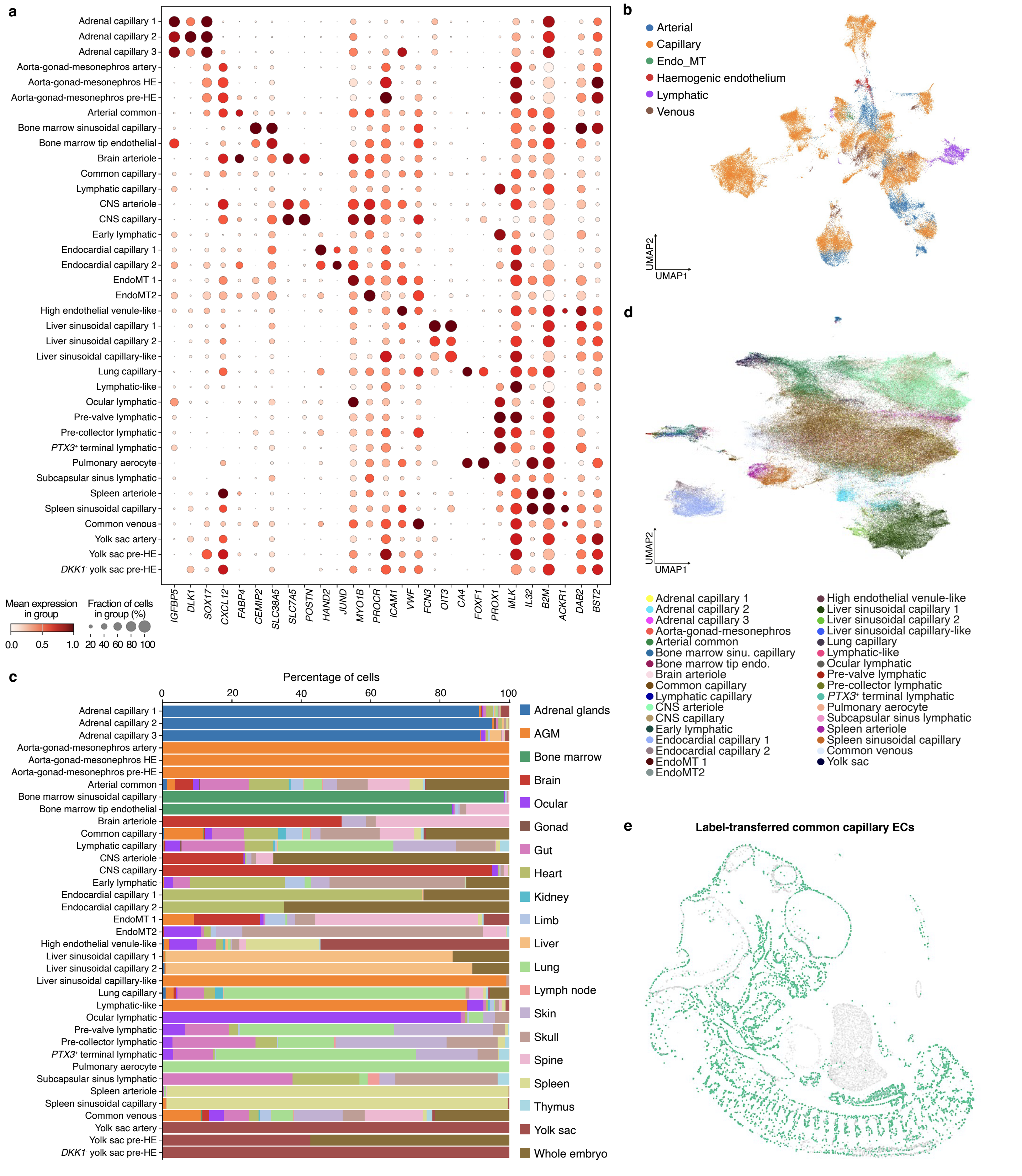

**Figure S21: Specialisation of organ-specific endothelial cells during human development**

**a)** Dotplot of canonical marker genes for each of the annotated EC types within the HDCA EC data. **b)** Broad annotations across the HDCA EC types, grouping cells in arterial, capillary, endothelial-mesenchymal transition-associated, hemogenic endothelial, lymphatic and venous identity. **c)** Bar plots for the percentage of cells within each EC type that is represented by each of the different organ systems within the HDCA. **d)** UMAP showing scVI integrated scRNA-seq data of ECs from the HDCA with 163,973 ECs isolated from the same 11 Xenium spatial transcriptomic sections used for niche annotation and analysis. scANVI was used to transfer labels from scRNA-seq to Xenium ECs. **e)** Label-transfer of the 'common capillary' annotation from scRNA-seq data to Xenium, with the spatial graph showing the generalised distribution of these cells, plotted using section S133 and representative of three sections analysed.

### Figure S22: Identification, characterisation and validation of lymphatic endothelial heterogeneity

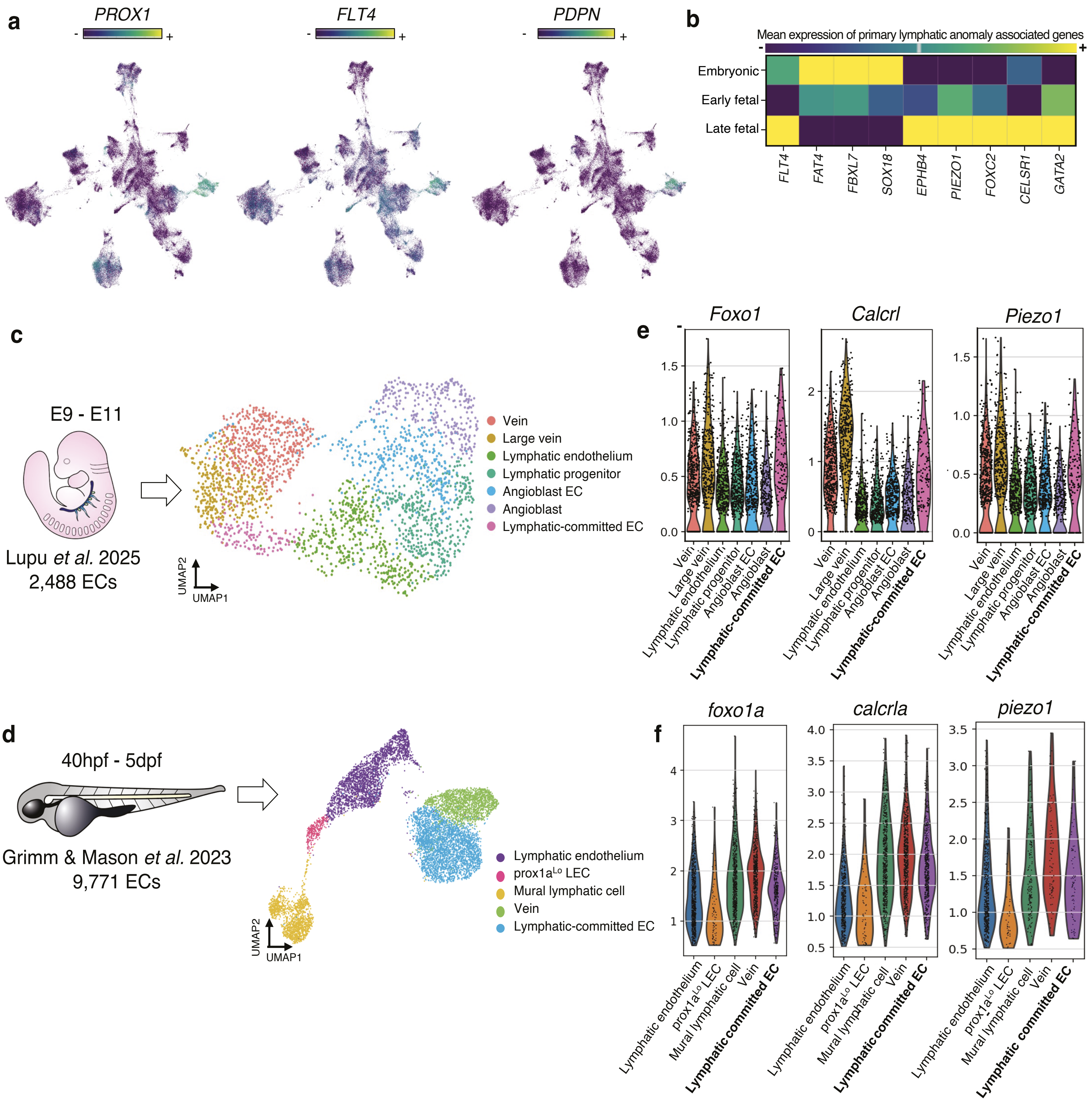

**Figure S22: Identification, characterisation and validation of lymphatic endothelial heterogeneity**

**a)** Feature plots showing expression of key marker genes for LECs within the HDCA EC dataset including *PROX1*, *VEGFR3* and *PDPN*, highlighting a distinct cluster of LECs. **b)** Heatmap showing LEC expression of genes associated with primary lymphatic anomalies across developmental time, grouped into embryonic, early fetal or late fetal periods. **c)** A previously published scRNA-seq dataset of mouse lymphatic development was profiled, using cell type annotations from the original analyses. **d)** A previously published scRNA-seq dataset of zebrafish lymphatic development was profiled, using cell type annotations from the original analyses. **e)** Expression of mouse orthologues from markers of early LECs, *Foxo1, Calcrl* and *Piezo1,* within different cell types within the mouse dataset. **f)** Expression of zebrafish orthologues from markers of early LECs, *foxo1a, calcrla* and *piezo1,* within different cell types within the mouse dataset.

**Figure S23: Lineage specification, maturation and regulatory dynamics in the developing human PNS and organoid model**

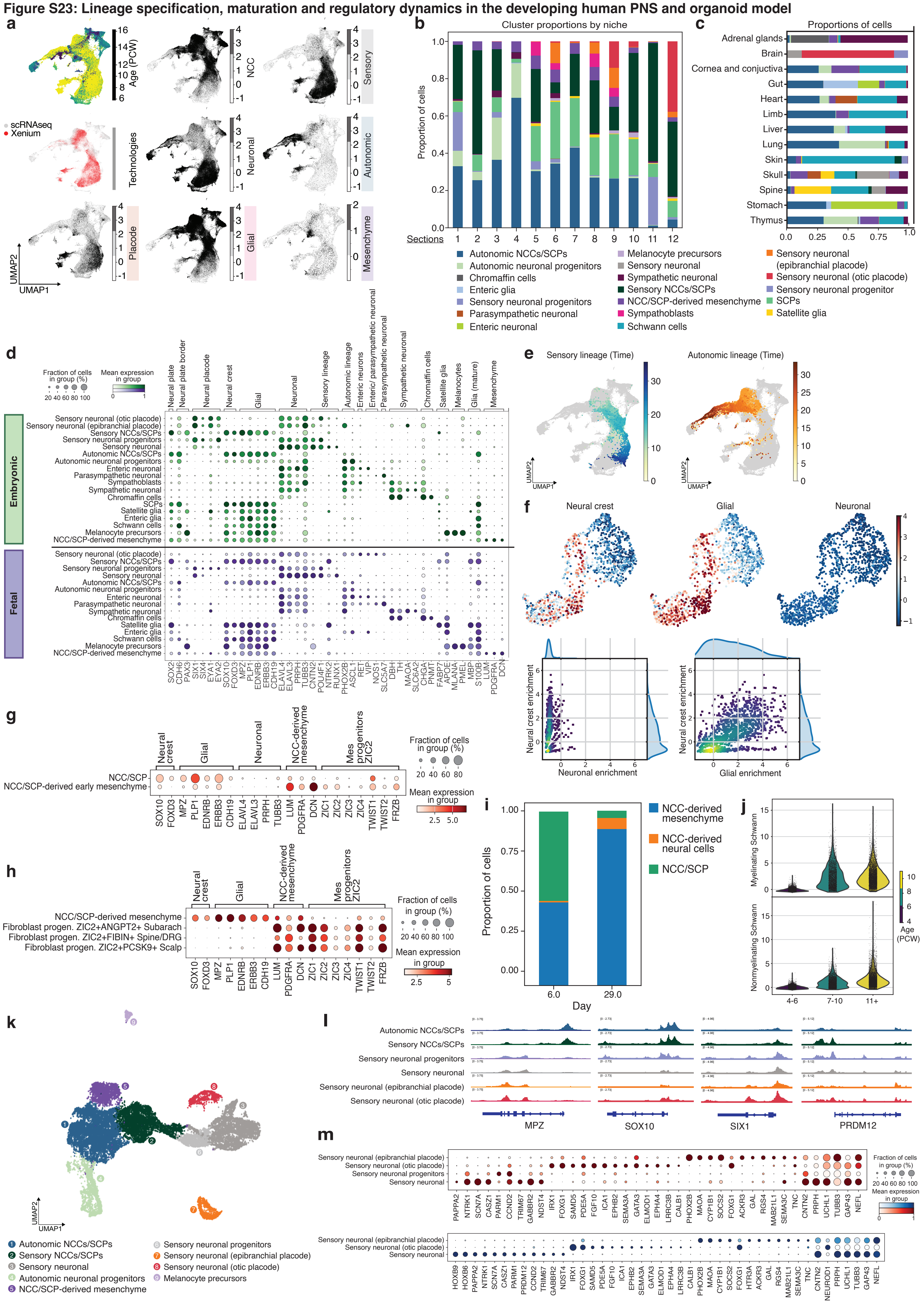

**Figure S23: Lineage specification, maturation and regulatory dynamics in the developing human PNS and organoid model**

**a)** UMAPs show the enrichment scores of the main PNS lineages across all the cells in the HDCA PNS dataset, as in **Fig. 5a**. **b)** Barplot depicting the cell type proportions per section origin only from scRNA-seq derived from the whole embryo (WE) dataset. **c)** As in panel b, but from public datasets. Only organs with > 300 cells are displayed. **d)** Dotplot of marker gene expression for PNS populations in **Fig .5a** divided by embryonic (< PCW 8) and fetal (> PCW 8) stage. **e)** UMAP visualisation of PNS population autonomic and sensory trajectories calculated with Cell2fate for the WE dataset. **f)** UMAPs (top) display the enrichment scores of 'Neural crest', 'Glial', and 'Neuronal' signatures of day 6 cranial neural crest cells derived from skin organoids[89]. Scatterplots (bottom) compare single cell enrichment scores of the 'Neural crest' signature (y-axis) with 'Glial' and 'Neuronal' signature scores (x-axis) across cells in UMAP space. **g**) Dotplot showing expression of genes associated with the 'Neural crest', 'Glial', 'Neuronal', 'NCC-derived mesenchyme' and '*ZIC*$^{hi}$ Mesenchymal progenitors' signatures in day 6 cranial neural crest cells and NCC/SCP-derived early mesenchyme from skin organoids[89]. **h**) Dotplot depicts gene expression of genes associated with the 'Neural crest', 'Glial', 'Neuronal', 'NCC-derived mesenchyme' and 'Mesenchymal progenitors *ZIC*$^{hi}$' signatures in NCC/SCP-derived mesenchyme populations in **Fig. 3 and 5**. **i)** Barplots depicting the proportion of NCC/SCP, NCC-derived neural cells (Sensory neuronal progenitors, SCPs, Satellite glia, Schwann cells and Sensory neuronal) and NCC-derived mesenchyme (NCC/SCP-derived mesenchyme, NCC/SCP-derived early mesenchyme) from skin organoids at day 6 and 29. **j)** Violin plots depicting the enrichment scores of myelinating and non-myelinating Schwann cells in the glial populations from **Fig. 5a**. **k)** Left: UMAP representation of the 6PCW WE single-nucleus multiome data. PNS cells are annotated into 9 distinct cell populations. Right: Dotplot shows the mean expression of known marker genes across PNS cell populations. **l)** Chromatin accessibility (ATAC) profiles of known marker genes across NCC/SCP and placode-derived cell populations. Profiles are shown as aggregated peak signals for each population. Panels for each gene were autoscaled to the maximum peak value to enable comparison across cell populations. **m)** Dotplot showing the differentially expressed genes between the different sensory neuronal populations in the scRNA-seq (top) and the Xenium (bottom).
