## Supplementary Figure 24 for "An integrated single cell and spatial omics atlas of human prenatal development" (from another version of this preprint: version 1, 2026-04-01)

### Figure S24: PNS distribution and interactions across tissue niches

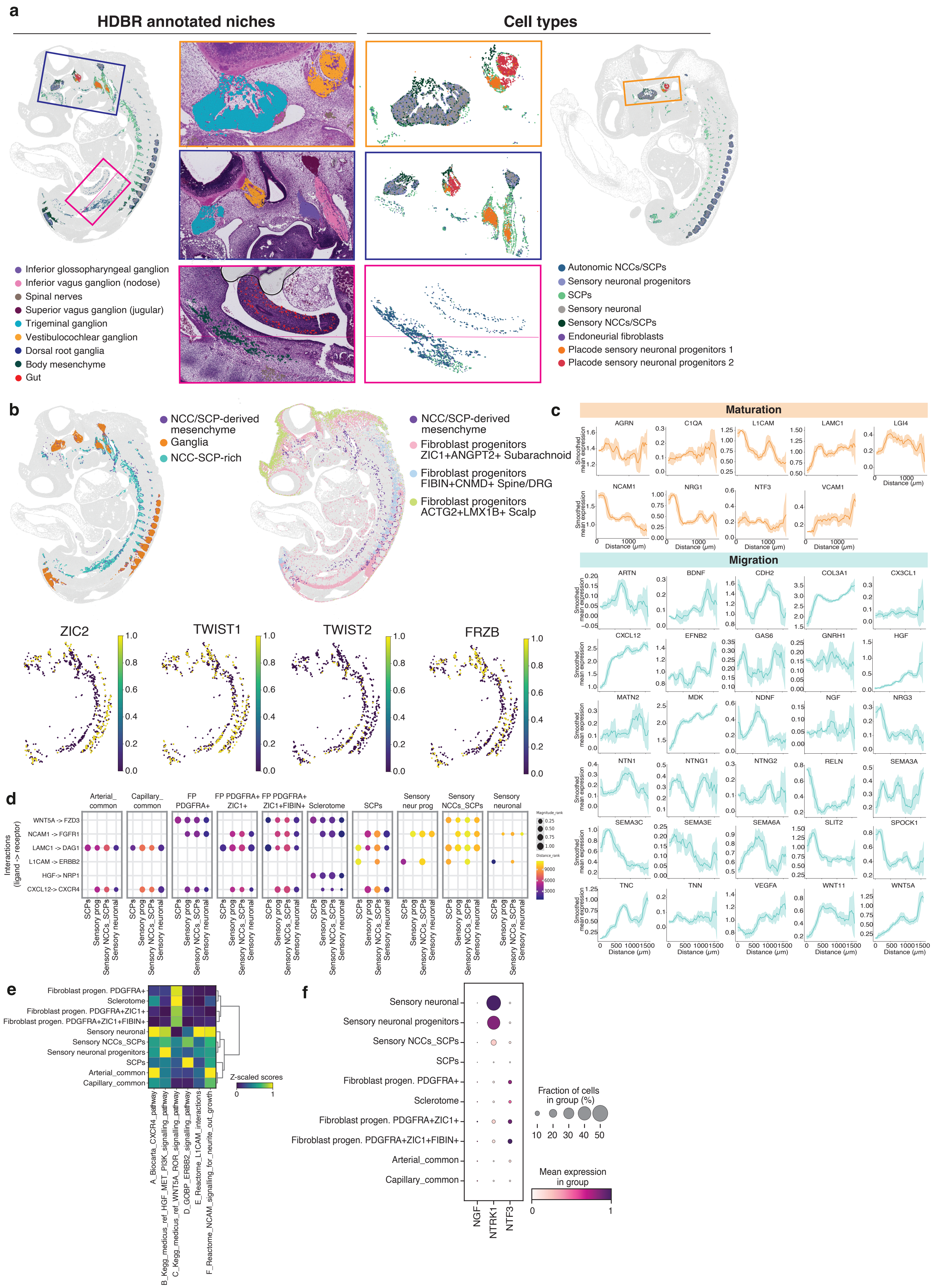

**Figure S24: PNS distribution and interactions across tissue niches**

**a)** Spatial location of the PNS populations of an embryo at ~6PCW (left) in 2 independent sections (S46 left, S113 right). Zoom into specific PNS areas (right) into head ganglia region, DRG and inferior hypogastric plexus. H&E, HDBR annotated niches and cell types. **b)** Spatial location of NCC/SCP-derived mesenchyme overlaid on PNS niches (left), mesenchymal progenitors *ZIC*$^{hi}$ populations (middle) and expression of genes associated with *ZIC*$^{hi}$ mesenchymal lineages (right; *ZIC2*, *TWIST1*, *TWIST2*, *FRZB*) in a sagittal slice S46 of an embryo at ~6PCW. **c)** Expression of maturation (orange) and migration (turquoise) ligands curated from literature across the distance from the DRG (proximal to distal planes) using Visium 10X Genomics section 121. **d)** Dotplot showing the ligand-receptor interactions between the cell subtypes found in the different DRG niches. Dot colour represents the distance rank between cell types and dot size reflected interaction magnitude. **e)** Heatmap of the enrichment scores using downstream signalling gene signatures for the receptors in (d) in the cell subtypes found in the different DRG niches. **f)** Dotplot showing the expression of genes associated with neuropathies affecting cranial sensory ganglia in the cell type populations in the DRG using Xenium 10X Genomics data.
